# Identification of a Monovalent Pseudo-Natural Product Degrader Class Supercharging Degradation of IDO1 by its native E3 KLHDC3

**DOI:** 10.1101/2024.07.10.602857

**Authors:** Elisabeth Hennes, Belén Lucas, Natalie S. Scholes, Xiu-Fen Cheng, Daniel C. Scott, Matthias Bischoff, Katharina Reich, Raphael Gasper, María Lucas, Teng Teng Xu, Lisa-Marie Pulvermacher, Lara Dötsch, Hana Imrichova, Alexandra Brause, Kesava Reddy Naredla, Sonja Sievers, Kamal Kumar, Petra Janning, Malte Gersch, Peter J. Murray, Brenda A. Schulman, Georg E. Winter, Slava Ziegler, Herbert Waldmann

**Affiliations:** Max-Planck-Institut für Molekulare Physiologie, Abteilung Chemische Biologie, Otto-Hahn-Straße 11, 44227 Dortmund, Germany; Technische Universität Dortmund, Fakultät Chemie und Chemische Biologie, Otto-Hahn-Straße 6, 44221 Dortmund, Germany; CeMM Research Center for Molecular Medicine of the Austrian Academy of Sciences, Vienna, Austria; Department of Structural Biology, St. Jude Children’s Research Hospital, Memphis, TN 38105, USA; Compound Management and Screening Center Otto-Hahn-Str.11, 44227 Dortmund, Germany; Max-Planck-Institut für Molekulare Physiologie, Zentrale Einheit für Kristallographie und Biophysik, Otto-Hahn-Straße 11, 44227 Dortmund, Germany; Instituto de Biomedicina y Biotecnología de Cantabria, Universidad de Cantabria-CSIC, C/ Albert Einstein 22, PCTCAN, 39011 Santander, Spain; Immunoregulation Research Group, Max Planck Institute of Biochemistry, Martinsried, Germany; Chemical Genomics Centre, Max-Planck-Institut für Molekulare Physiologie, 44227 Dortmund, Germany; Department of Molecular Machines and Signaling, Max Planck Institute of Biochemistry, Martinsried, Germany

## Abstract

Targeted protein degradation (TPD) modulates protein function beyond inhibition of enzyme activity or protein-protein interactions. Most degrader drugs function by directly mediating proximity between a neosubstrate and hijacked E3 ligase. Here, we identified pseudo-natural products derived from (-)-myrtanol, termed iDegs that inhibit and induce degradation of the immunomodulatory enzyme indoleamine-2,3-dioxygenase 1 (IDO1) by a distinct mechanism. iDegs boost IDO1 ubiquitination and degradation by the cullin-RING E3 ligase CRL2^KLHDC3^, which we identified to natively mediate ubiquitin-mediated degradation of IDO1. Therefore, iDegs increase IDO1 turnover using the native proteolytic pathway. In contrast to clinically explored IDO1 inhibitors, iDegs reduce formation of kynurenine by both inhibition and induced degradation of the enzyme and, thus, would also modulate non-enzymatic functions of IDO1. This unique mechanism of action may open up new therapeutic opportunities for the treatment of cancer beyond classical inhibition of IDO1.

## Main

Small-molecule mediated targeted protein degradation (TPD) is a powerful approach to modulate protein functions beyond enzymatic activity and protein-protein interactions.^1^ Current strategies for TPD rely on small molecules (“degraders”) that induce molecular proximity between a neosubstrate and hijacked E3 ubiquitin ligase.^2^ Degraders can either be heterobifunctional, harboring distinct entities for engaging both target and E3 via dedicated ligands (“PROTACs”), or monovalent, binding to either the target or the ligase to adapt its protein surface and induce a highly cooperative tripartite assembly (Molecular Glue Degraders, MGDs). Both degrader types have been applied successfully for a variety of targets and have entered the clinics^3^ and they highlight the route to novel opportunities for degrader development. In particular, the discovery of new degrader chemotypes which may induce recruitment of new E3 ligases and new TPD strategies are in high demand.

Natural products (NPs) and analogs have yielded E3 ligase ligands and mono- and bivalent inducers of protein degradation^4^, raising the possibility that new degrader chemotypes could be derived from NPs. Pseudo-natural products (PNPs) combine natural-product fragments in arrangements and combinations not observed in NPs. They retain the biological relevance of NPs but open new chemical space and therefore may have unexpected and novel targets,^5,6^ such that exploration of their bioactivity in particular in unbiased cell-based and phenotypic screens^5,7^ may identify novel small-molecule degrader chemotypes and E3 ligases.

The heme-binding enzyme indolamine 2,3-dioxygenase 1 (IDO1) converts tryptophan (Trp) to kynurenine (Kyn). Trp shortage and Kyn elevation have numerous downstream effects including reduced Teff cell proliferation and promotion of Treg cell differentiation, generation of ligands for the aryl-hydrocarbon receptor and ferroptosis-suppressive kynurenine derivatives, all of which are linked to reduced anti-tumor immunity.^8–12^ Recently, the Epstein Barr virus (EBV) has been reported to induce IDO1 expression, which is related to EBV-associated lymphoma.^13^ Moreover, IDO1 and Kyn have been linked to neurodegeneration and increased IDO1 expression in amyloid and tau pathologies is associated with impaired hippocampal glucose metabolism, spatial memory and synaptic plasticity.^14,15^ In general, clinical exploration of different IDO1 inhibitors for cancer treatment has,^16–22^ despite encouraging preclinical data,^23,24^ met with limited success.^25,26^ Plausible reasons include that IDO1 may have non-enzymatic functions in immunosuppression and in glioblastoma ^27–34^. Nevertheless, there are eight active and two recruiting clinical trials with IDO1 inhibitors^35^ underscoring the potential and the continuous interest in IDO1 modulation for disease treatment. The limitations arising from the use of enzyme inhibition only^36^ may be overcome by IDO1 degradation, which would eliminate both enzymatic activity and signaling function. Initial investigations of IDO1-directed PROTACs^37,38^ support this notion, but small-molecule monovalent degraders or molecular glues targeting IDO1 have yet to be identified.

In the course of a cell-based screen for small molecules that reduce Kyn levels, we identified a class of pseudo-natural products derived from (-)-myrtanol, termed iDegs, that both inhibit IDO1 and induce IDO1 degradation. iDegs induce structural changes in IDO1 that cause enhanced ubiquitination and augmented degradation by CRL2^KLHDC3^, a ligase we identified to mediate also the ubiquitination and native degradation of IDO1. Our work defines a unique mechanism of action, a new type of degrader, and identifies a new E3-ligase previously not employed for small molecule-mediated protein degradation.

### Identification of IDO1 degraders

A library of 157,332 small molecules, obtained from commercial and academic sources and developed in house was screened in a cell-based assay (Kyn assay, see Extended Data Fig. 1a) that quantified Kyn, the product of IDO1 reaction. So as to screen under conditions that would be sensitive to IDO1 levels, we considered that IDO1 expression can be induced by interferon gamma (IFN-γ).^39^ We thus performed the screen in IFN-γ-stimulated BxPC3 cells^21^. A hit from the screen, the (-)-myrtanol-derived pseudo-natural product hereafter termed iDeg-1 (see Supporting Scheme 1 for the synthesis), inhibited Kyn formation with an IC50 of 0.83 ± 0.31 µM in the screening assay (Fig. 1a), which we confirmed using orthogonal Kyn assays in IFN-γ-stimulated BxPC3-, SKOV-3- and HeLa cells (IC50 values of 1.1 ± 0.1 µM, 1.6 ± 0.3 µM and 1.7 ± 1.2 µM, respectively, Fig. 1b and Extended Data Fig. 1b-c). Further indication that iDeg-1 targets IDO1 came from a cell viability assay in 2D and 3D cultures^40,41^: iDeg-1 reduced IDO1-dependent SKOV-3 cell death induced by IFN-γ (Extended Data Fig. 1d-e). iDeg-1 only slightly affected IDO1 enzymatic activity (Extended Data Fig. 1f) and did not impair *IDO1* transcription (Extended Data Fig. 1g-h). Instead, iDeg-1 dose-dependently reduced IDO1 protein levels up to 45 ± 15 % at 10 µM after 24 h (Fig. 1c-d) without inhibiting *in vitro* translation of IDO1 or global protein translation (Extended Data Fig. 1i-n).

**Fig. 1:**
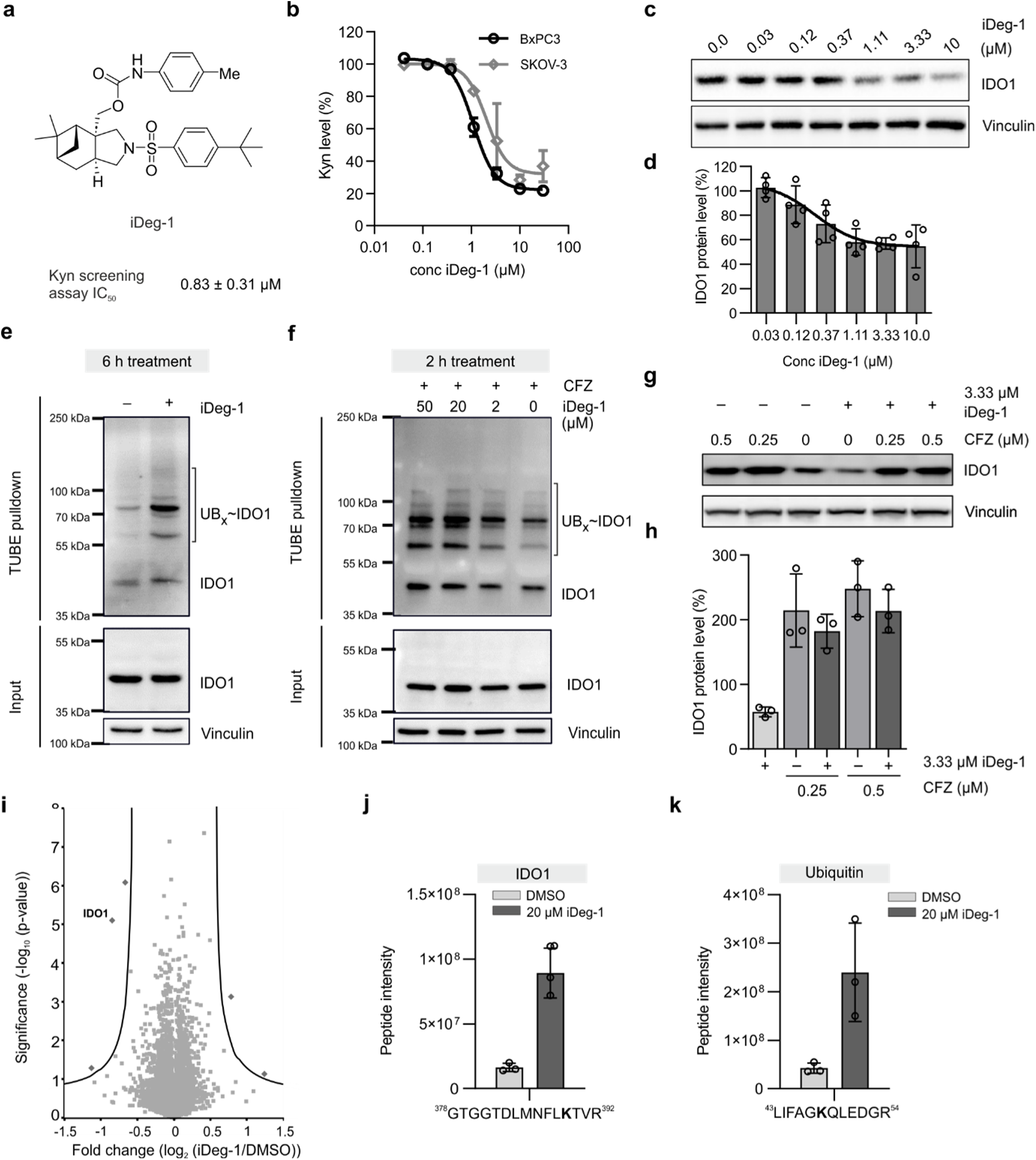
iDeg-1 reduces IDO1 protein levels via the UPS. **a,** Structure of screening hit iDeg-1 and IC_50_ value in the Kyn assay in BxPC3 cells. Mean value ± SD, n = 3. **b,** Kyn assay in BxPC3- and SKOV-3 cells after treatment with iDeg-1 and 50 or 5 ng/mL IFN-γ, respectively, for 48 h prior to detection of Kyn levels utilizing *p*-DMAB. **c,d,** IDO1 protein levels in BxPC3 cells upon treatment with IFN-γ and iDeg-1 for 24 h. Representative immunoblots (c) and quantified band intensities from c (d). Mean values ± SD, n = 4. **e,f,** TUBE pulldown after treatment of IFN-γ-stimulated BxPC3 cells with iDeg-1 or DMSO. **e,** Cells were treated for 6 h with 50 µM iDeg-1 prior to the TUBE pulldown. Representative immunoblots of n = 3. **f,** Cells were treated with 450 nM carfilzomib (CFZ) 60 min prior to the addition of iDeg-1 or DMSO for 2 h followed by TUBE pulldown. Representative immunoblots of n = 3 for IDO1. UB: ubiquitin. **g,h,** HEK239T cells were electroporated with rhIDO1 protein. Cells were treated with CFZ for 30 min prior to the addition of 3.33 µM iDeg-1 and further incubation for 6 h. **g,** Representative immunoblot of n = 3. **h,** Quantified band intensities from g represent samples treated with compound relative to DMSO (set to 100 %). Mean values ± SD, n = 3 or n = 4. **i,** Volcano plot of iDeg-1-induced changes in the global proteome. HEK239T cells were electroporated with rhIDO1 protein followed by treatment with 10 µM iDeg-1 or DMSO for 6 h and MS analysis. In total, 7,541 proteins were detected. Plot generated using Perseus (S0 = 0.5, FDR = 0.01). **j,k,** IDO1 immunoprecipitation (IP) and identified peptide of IDO1 (**j**) or ubiquitin (**k**) with diGly modification. IFN-γ-stimulated BxPC3 cells were treated with 20 µM iDeg-1 or DMSO for 6 h prior to the IP. Uncropped blots are shown in Extended Data Fig. 11.

In order to determine whether iDeg-1 induces degradation via the ubiquitin proteasome system (UPS), IFN-γ-stimulated BxPC3 cells were treated with iDeg-1 for 6 h (Fig. 1e), or for 2 and 4 h after pretreatment with the proteasome inhibitor carfilzomib (CFZ; Fig. 1f and Extended Data Fig. 2a). Increased polyubiquitination was detected with a tandem ubiquitin binding entity (TUBE) pulldown from cell lysates.^42^ We found iDeg-1 induced IDO1 ubiquitination shortly after compound addition. By contrast, IDO1 protein reduction was only detectable after 24 h in IFN-γ-stimulated BxPC3 cells (Fig 1c-d). Since the capacity to induce degradation may be masked by the ongoing IFN-γ-stimulated production of IDO1, we directly introduced enzymatically active recombinant human IDO1 protein (rhIDO1) into HEK293T cells by electroporation (HEK^rhIDO1^ cells). iDeg-1 dose-dependently reduced Kyn amounts with an IC50 value of 0.45 ± 0.1 µM (Extended Data Fig. 2b-c) and lowered IDO1 protein after 6 h by 46 % at a concentration of 10 µM (Extended Data Fig. 2d-e). CFZ inhibited degradation (Fig. 1g-h and Extended Data Fig. 2f), indicating involvement of the UPS. iDeg-1 also inhibited Kyn production in HEK293T cells that transiently express IDO1 with an IC50 of 0.42 ± 0.2 µM (Extended Data Fig. 2g). Therefore, iDeg-1 depletes IDO1 independently of the effects of IFN-γ.

High specificity of iDeg-1 for IDO1 depletion was detected using global proteome analysis of HEK^rhIDO1^ cells after 6 h of treatment with iDeg-1 (Fig. 1i and Extended Data Table 1). Analysis of ubiquitinated proteins for a diglycine (diGly) attached to lysine residues (K-ε-diglycine) which were modified with ubiquitin^43^, after iDeg-1 treatment and IDO1 immunoprecipitation uncovered K389 of IDO1 as a site of ubiquitination (Fig. 1j and (Extended Data Fig. 3a). In addition, we observed a 3.5-fold increase in ubiquitin levels in the IDO1 immunoprecipitate after treatment with iDeg-1 confirming iDeg-1-induced IDO1 ubiquitination (Extended Data Fig. 3b). DiGly analysis further revealed K48 linkages in the ubiquitin chains (Fig. 1k and Extended Data Fig. 3c), which are associated with protein degradation via the UPS ^44^.

To determine if iDeg-1 directly engages IDO1 in cells, we performed a cellular thermal shift assay (CETSA)^45^. In the presence of iDeg-1, IDO1 showed thermal stabilization with a shift in the melting temperature (Δ*T*m) of 3.5 ± 0.4 °C (Fig. 2a-b). Thermal stabilization of IDO1 by iDeg-1 was dose-dependent as demonstrated by an isothermal CETSA experiment at 50°C (Extended Data Fig. 4a-b).

**Fig. 2:**
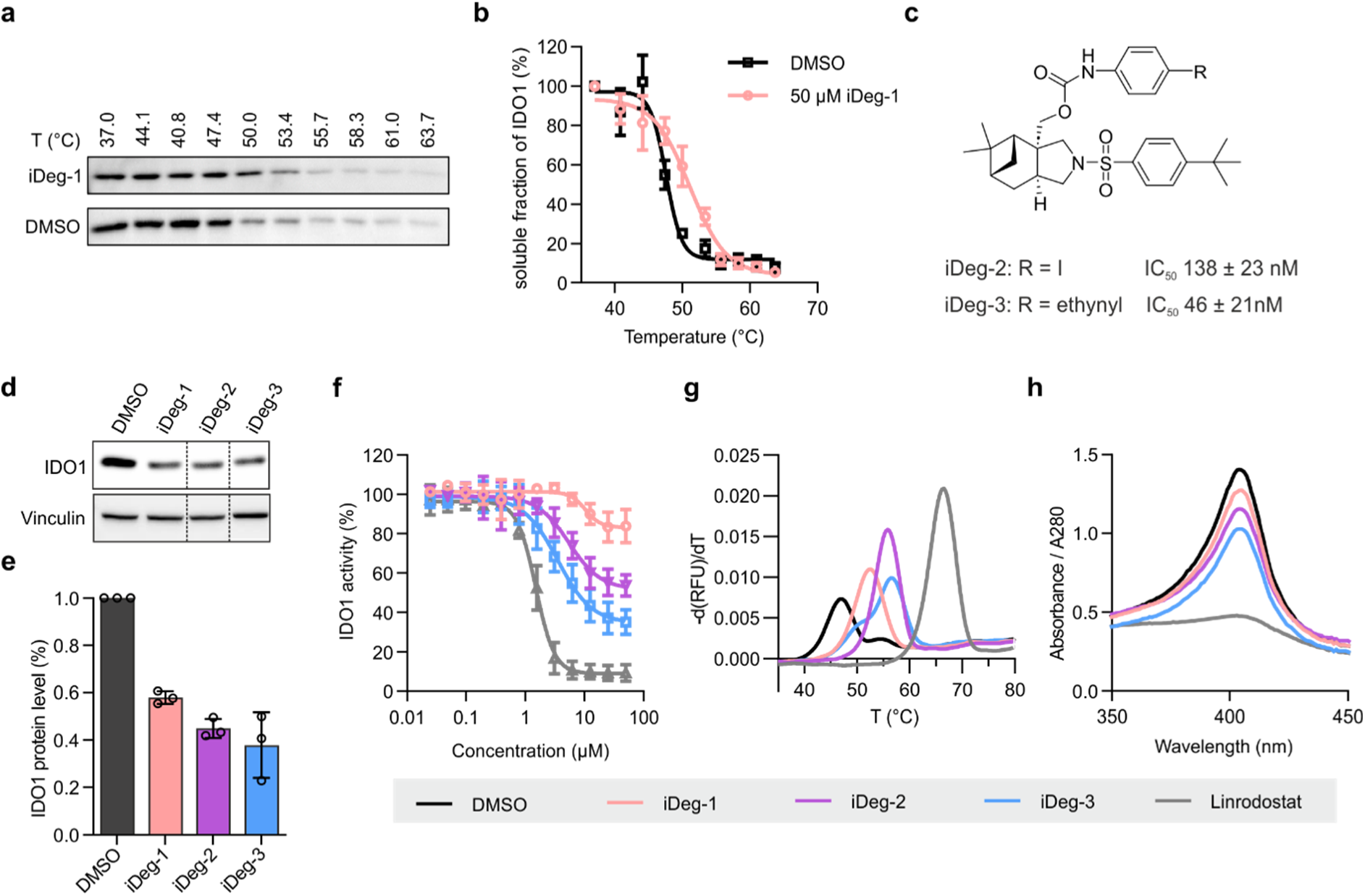
iDegs interact with IDO1. **a,b,** CETSA in intact SKOV-3 cells treated with 50 µM iDeg-1 or DMSO for 1 h followed by heat treatment and immunoblotting. **a,** Representative immunoblot for IDO1. **b,** Quantification of band intensities from a. Mean values ± SD, n = 3. **c,** Structures of iDeg-2 and iDeg-3 and IC_50_ values in the Kyn assay in BxPC3 cells. Mean values ± SD, n = 3. **d,e,** Influence of iDeg-1, 2 and 3 on IDO1 protein levels in BxPC3 cells. Cells were treated with IFN-γ and the compounds (3.33 µM) for 24 h prior to immunoblotting (d). Quantification of band intensities shown in e. Mean values ± SD, n = 3. **f,** Influence on the *in vitro* rhIDO1 activity. rhIDO1 was pre-incubated with the compounds at 37°C for 90 min prior to detection of Kyn levels using *p*-DMAB. Mean values ± SD, n = 3. **g,** rhIDO1 thermal stability in presence of 50 µM iDeg-1, 2 or 3 or DMSO and the apo-IDO1 inhibitor linrodostat (50 µM) using nanoDSF. rhIDO1 and compounds were pre-incubated for 3 h at 37°C prior to the measurement. Representative result (n = 3). **h,** Detection of heme-bound IDO1 by means of UV/Vis spectroscopy in presence of iDeg-1, 2 or 3 (100 µM), DMSO or linrodostat (100 µM). Incubation at 37°C for 3 h. Representative data for n = 3. Uncropped blots are shown in Extended Data Fig. 12.

Initial structure-activity relationship investigations identified two analogs with enhanced activity in the Kyn assay as compared to iDeg-1. These have an iodine (termed iDeg-2, IC50 = 138 ± 23 nM) or an alkyne group (termed iDeg-3, IC50 = 46 ± 21 nM) in *para* position of the phenyl carbamate (Fig. 2c, Extended Data Fig. 4c-d). In BxPC3 cells exposed continuously to IFN-γ, iDeg-2 and iDeg-3 reduced IDO1 protein levels by 55 ± 3 % and 62 ± 11 % respectively as compared to 42 ± 2 % by iDeg-1 at 3.33 µM (Fig. 2d-e). iDeg-2 and 3 also partially inhibited the enzyme *in vitro* (Fig. 2f). Thermal stabilization of rhIDO1 was detected for the three compounds using nanoDSF (Fig. 2g), indicating direct binding to the protein. UV/Vis spectroscopic analysis revealed that in the presence of iDeg-1-3 the specific *Soret* absorbance peak of heme-bound IDO1 (holo-IDO1, Fig. 2h) is reduced, showing that iDegs displace heme and, with different potencies, bind to apo-IDO1. Accordingly, addition of hemin reduced the potency of iDeg-1, -2 and -3 in the Kyn assay (Extended Data Fig. 4e-g) and dose-dependently elevated Kyn levels in the presence of iDeg-2 (Extended Data Fig. 4h). As we detected both enzymatic **i**nhibition and **<u>deg</u>**radation of IDO1 by iDeg-3, the compound class was termed **iDeg**.

### Crystal structure shows iDegs alter conformation of the IDO1 C-terminal region

To gain insights into the mechanism by which iDegs affect IDO1 structure, we performed X-ray crystallography. The co-crystal structure of apo-IDO1 in complex with iDeg-2 at 1.6 Å resolution (PDB ID 9FOH, Extended Data Table 2 and Extended Data Fig. 5a) revealed two remarkable effects of iDeg binding. First, iDeg-2 resides in the heme binding site in apo-IDO1. The phenyl carbamate occupies the previously identified lipophilic pocket A^46^ in the distal heme site, while the pyrrolidine and the sulfonyl group occupy the heme-binding pocket. The monoterpene scaffold only slightly protrudes into the D-pocket, and the *tert*-butyl phenyl group is located in the solvent-exposed B-pocket (Fig. 3a-c). Interestingly, binding to this latter pocket has been observed before mainly for holo-IDO1 inhibitors (Extended Data Fig. 5b-c). iDeg-2 binding occurs through a large number of hydrophobic interactions, a water-bridged hydrogen bond between the carbamate nitrogen and the hydroxyl group of S167 and a hydrogen bond between the sulphonyl oxygen of iDeg-2 and H346 (Fig. 3d-e). As many of these same IDO1 residues are responsible for heme coordination, this binding mode is mutually-exclusive with heme and ensures that iDegs can only bind apo-IDO1. Second, comparison to all IDO1 structures revealed iDeg-2 induces a striking conformational rearrangement (Extended Data Fig. 5d-f). In the presence of iDeg-2, there was no detectable electron density for the C-terminal K-helix. This absence is not attributed to crystal packing forces, as the K-helix is present in the apo-IDO1-apoxidole crystal structure with identical space group, unit cell and similar crystallization conditions (PDB ID: 8ABX, Extended Data Fig. S5d-e).^47^ An overlay of the structures of IDO1 bound to iDeg-2 with the apo-IDO1 inhibitor linrodostat revealed the underlying basis: helices B, C, F, H and J are all reoriented and the J-helix is also remodeled in the iDeg-2 complex (Fig. 3f and Extended Data Fig. 5f). In all previously published IDO1 structures, the K-helix is embraced through F, J, H helices and the EF-loop (Extended Data Fig. 5e). In contrast, in the iDeg-2-bound complex, tethers to the K-helix are weakened due to cumulative movements of several amino acids within the J-helix. Specifically, a substantial rotation and translation of H346 shifts the J-helix C-terminus towards the iDeg-2 binding pocket. This movement generates a steric clash between R343 and F270, forcing R343 to adopt an alternative conformation that clashes with the K-helix (Fig. 3g and Extended Data Fig. 5f). The combination of this steric clash and weakened interactions between the K-helix and surrounding helices F, J and H presumably increases the K-helix flexibility, accounting for its invisibility in the electron density map. As previously reported IDO1 inhibitors do not significantly alter the overall structure of IDO1 compared to holo-IDO1^46^ (Extended Data Fig. 5c-d), the observed conformational rearrangements represent a novel and unique binding mode of the iDegs.

**Fig. 3.**
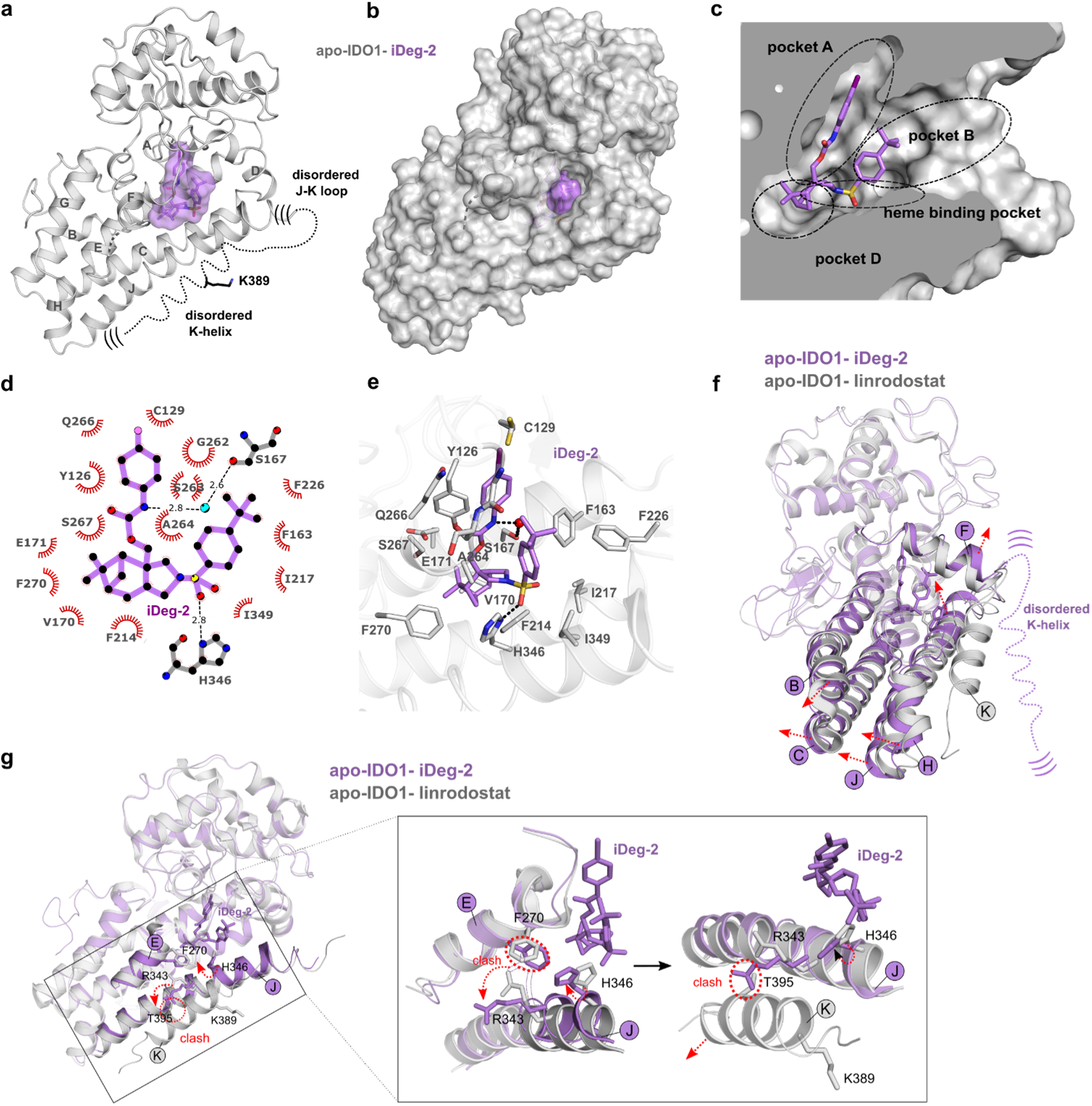
Co-crystal structure reveals that iDeg-2 binds to apo-IDO1 and induces conformational changes in its C-terminal region. **a**, Cartoon diagram of the IDO1-iDeg-2 structure. Dashed lines indicate regions lacking electron density, including the E-F loop, J-K loop, and K-helix, with parentheses denoting their proposed flexibility. The position of ubiquitinated K389 within the K-helix is highlighted. The α-helices of the large domain (residues 121-403) are labeled A to K. **b,** Surface representation of apo-IDO1 with iDeg-2 bound. **c,** Surface model (cut-in side view) showing iDeg-2 in the active site of apo-IDO1. Hydrophobic pockets A, B, D, and the heme binding site are indicated. **d**, LigPlot schematic illustration of interactions between iDeg-2-and apo-IDO1. Hydrogen bonds between iDeg-2 and apo-IDO1 or a water molecule (cyan sphere) are shown as black dashed lines. **e,** Zoom-in view of amino acids involved in iDeg-2 binding. Hydrogen bonds are indicated by black dotted lines. **f**, Overlay of apo-IDO1-iDeg-2 (violet) and apo-IDO1-linrodostat (grey, PDB ID: 6DPR-B) structures. Red arrows highlight helix shifts. **g,** Conceptual model illustrating how iDeg-2 binding triggers local and long-distance structural perturbations (dotted arrows). iDeg-2 binding induces a conformational shift in H343, moving the J-helix toward the iDeg-2 binding pocket, including R343. To avoid steric clash of R343 (dashed lines) with F270, R343 adopts an alternative conformation, which clashes with T395 and ultimately leads to the detachment of the K-helix.

Thus, the structure provided two key mechanistic insights into iDeg function. First, since the compound is deeply buried inside the active site, it seemed unlikely that iDegs directly bind an E3 ligase. Second, iDeg-induced degradation could possibly involve increased accessibility of the C-terminal K-helix bearing the ubiquitinated K389, which could not be achieved by the contrasting conformation with all published IDO1 inhibitors.

### iDegs promote IDO1 ubiquitination by KLHDC3

To identify the E3 responsible for iDeg-mediated IDO1 degradation, we designed a FACS-based CRISPR-Cas9 screen using a custom sgRNA library targeting 1301 ubiquitin-associated human genes (6 sgRNAs per gene)^48^. We generated an IDO1 stability reporter following our previously reported design in KBM7 cells harboring an inducible Cas9 cassette^49^ (Fig 4a, Extended Data Table 3). In brief, IDO1 is expressed as tagBFP fusion followed by the self-cleaving P2A site and mCherry for assay normalization. Supporting a role of the C-terminal region of IDO1, only N-terminally fused BFP enabled iDeg-induced reporter degradation (Fig 4b). Next, cells were transduced with the custom sgRNA library. After selection for sgRNA positive cells, Cas9 expression and subsequent gene editing was induced via doxycycline for 72 h prior to 14 h compound treatment. Finally, cells were enriched for increased or decreased BFP levels using FACS and the corresponding sgRNAs were quantified by deep sequencing (Fig. 4d-e and Extended Data Fig. 6b and 7), revealing genes functionally required for iDeg-induced degradation.

**Fig. 4:**
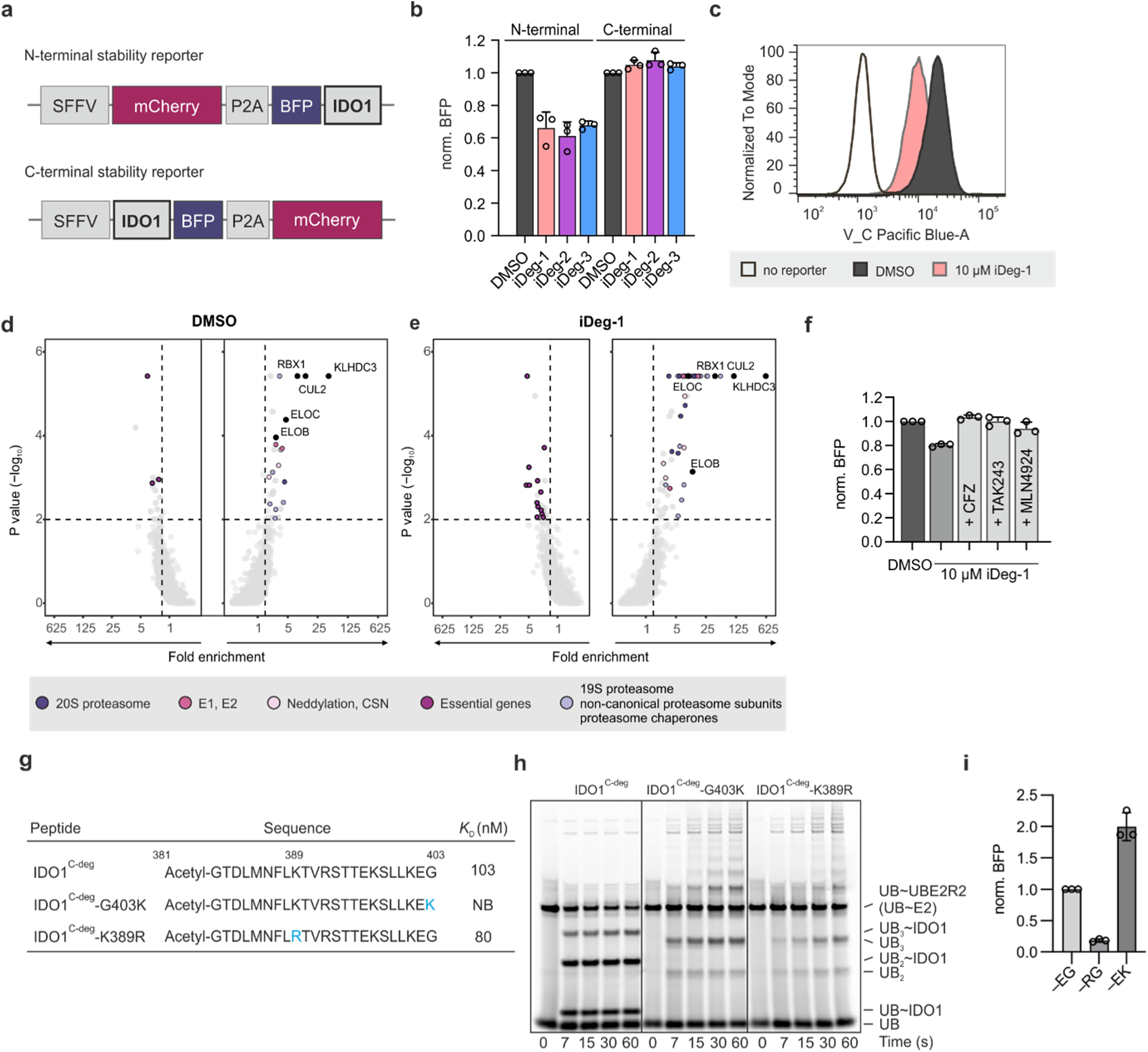
KLHDC3 is involved in iDeg-mediated IDO1 degradation. **a,** IDO1 stability reporter design. **b,** Detection of IDO1 levels using the IDO1 stability reporters. KBM7 IDO1 reporter cells were treated with iDeg-1, 2 or 3 (1 µM) for 24 h prior to detection of IDO1 levels using flow cytometry. Normalized BFP to mCherry ratios (norm. BFP) were calculated per genotype respectively. Mean values ± SD, n = 3. **c,** Representative histogram for iDeg-1 mediated depletion of BFP-IDO1 (24h, 10 µM). **d,** Identification of genes required for native IDO1 degradation. **e,** Identification of genes required for iDeg-1-mediated IDO1 degradation. CSN: COP9 signalosome. **f,** IDO1 depletion is rescued by 10 h co-treatment with either CFZ, TAK243 or MLN4924 (1 µM each). Mean values ± SD, n = 3. **g,** Binding affinities of IDO1 C-terminal peptide IDO1^C-deg^ and peptide mutants (IDO1^C-deg^-G403K and IDO1^C-deg^-K389R) to KLHDC3 determined using ITC. NB: no binding. **h,** Fluorescent scan of pulse-chase assay monitoring ubiquitination of C-terminal IDO1 peptides by UBE2R2. UBE2R2 was “pulse” loaded with fluorescent ubiquitin, and the resulting UBE2R2∼UB thiolester was added to NEDD8∼CRL2^KLHDC3^ incubated with the indicated peptides. Samples were quenched with SDS sample buffer at the indicated timepoints. **i,** IDO1 stability reporter variants in KBM7 cells measured by flow cytometry and depicted normalized to the wt (i.e., –EG) reporter. Mean values ± SD, n = 3. UB: ubiquitin.

As expected, knockout of genes coding for proteins of the 19S and 20S proteasome or involved in neddylation counteracted iDeg activity and increased IDO1 levels (Fig. 4d-e and Extended Data Fig. 6b-c), thus phenocopying the IDO1 stability reporter behavior upon chemical perturbation of the proteasome (CFZ), the ubiquitin-activating enzyme (TAK243) or of the neddylation machinery (MLN4924) (Fig 4f). Importantly, we further identified the cullin-RING ligase (CRL) complex including cullin2 (CUL2), RBX1, elongin B/C (EloB and EloC) and the Kelch domain containing protein 3 (KLHDC3) as required for IDO1 degradation.

Unexpectedly, genetic disruption of the CRL2^KLHDC3^ complex also affected baseline IDO1 turnover under vehicle (DMSO) treatment. In the presence of iDegs, however, knockout of these genes led to an even higher enrichment of the corresponding sgRNAs (Fig. 4e and Extended Data Fig. 6b-c). This indicated that the compounds enhance the efficiency of IDO1 degradation. Contrary to classical degrader modalities such as PROTACs or molecular glue degraders, which typically function by inducing proximity between an E3 and a target that is functionally inconsequential in the absence of the small molecule, iDegs thus appear to promote a native route for IDO1 turnover.

### IDO1 is a natural substrate of CRL2^KLHDC3^

The CRL substrate receptor KLHDC3 has not yet been employed for small molecule-induced protein degradation. KLHDC3 recognizes C-degrons ^50,51^, which explains why the C-terminally fused IDO1-BFP reporter was not degraded while the BPF-IDO1 reporter was (Fig. 4b). The C-terminal EG-sequence of human IDO1 is consistent with distinguishing feature of a KLHDC3 C-degron, which is a C-terminal glycine ^50–52^. In agreement, the peptide IDO1^C-deg^ corresponding to the C-terminal amino acid sequence 381-403 bound to KLHDC3 with a *K*D of 103 nM (Fig. 4g and Extended Data Fig. 8a). Exchanging the C-terminal glycine by a lysine (peptide IDO1^C-deg^-G403K) completely abolished binding to KLHDC3. Replacement of K389, which is ubiquitinated (peptide IDO1^C-deg^-K389R), did not affect formation of the E3-degron complex (Fig. 4g and Extended Data Fig. 8a).

In order to test if IDO1 is indeed a direct substrate of CRL2^KLHDC3^, we reconstituted biochemical ubiquitination assays. Since KLHDC3-EloB/C has been shown to form degron-mimic mediated auto-inhibited tetrameric assemblies, in which the C-degron binding site is occluded by the degron mimic^53^, we simplified the assay by using a C-terminal Gly-to-Lys mutant of KLHDC3 that is strictly monomeric. To specifically monitor ubiquitin transfer, assays were performed in “pulse-chase” format. The pulse reaction generates a thioester-linked E2∼ubiquitin conjugate using fluorescently-labeled ubiquitin that is readily monitored. Next, E3 and substrate are added, and fluorescent ubiquitin transfer from E2 (UBE2R2) to substrate and subsequently to a substrate-linked ubiquitin is observed over time. The wildtype peptide IDO1^C-deg^ was ubiquitinated *in vitro* in a CRL2^KLHDC3^-dependent manner, while the mutant versions were not (Fig. 4h and Extended Data Fig. 8b). In cells, mutation of the C-terminal glycine increased IDO1 amounts, whereas mutation of the non-optimal –EG degron to an optimal –RG C-terminus reduced IDO1 abundance (Fig. 4i). These findings demonstrate that IDO1 is a substrate of KLHDC3 and that the C-terminal glycine-based degron is essential for interaction with the E3 ligase.

Further exploration of the structure-activity relationship (SAR) led to the identification of the inhibitor and degrader iDeg-6 with an IC50 of 16 ± 5 nM in the Kyn assay (Fig. 5a and Extended Data Fig. 9a). In a modified setup including washout of IFN-γ followed by compound addition to avoid continuous IDO1 expression, iDeg-6 most efficiently and potently depleted IDO1 in cells with Dmax of 70 % at 100 nM and DC50 for IDO1 degradation of 6.5 ± 3 nM (Fig. 5a-c and Extended Data Fig. 9b-c). *In vitro* iDeg-6 inhibited IDO1 activity nearly completely and more potently than iDeg-1-3 with an IC50 of 1.6 ± 0.3 µM. The compound induced thermal stabilization of the protein and heme displacement to a higher extent as compared to iDeg-1, -2 and -3 (Extended Data Fig. 9d-f). We therefore used iDeg-6 for further validation.

**Fig. 5:**
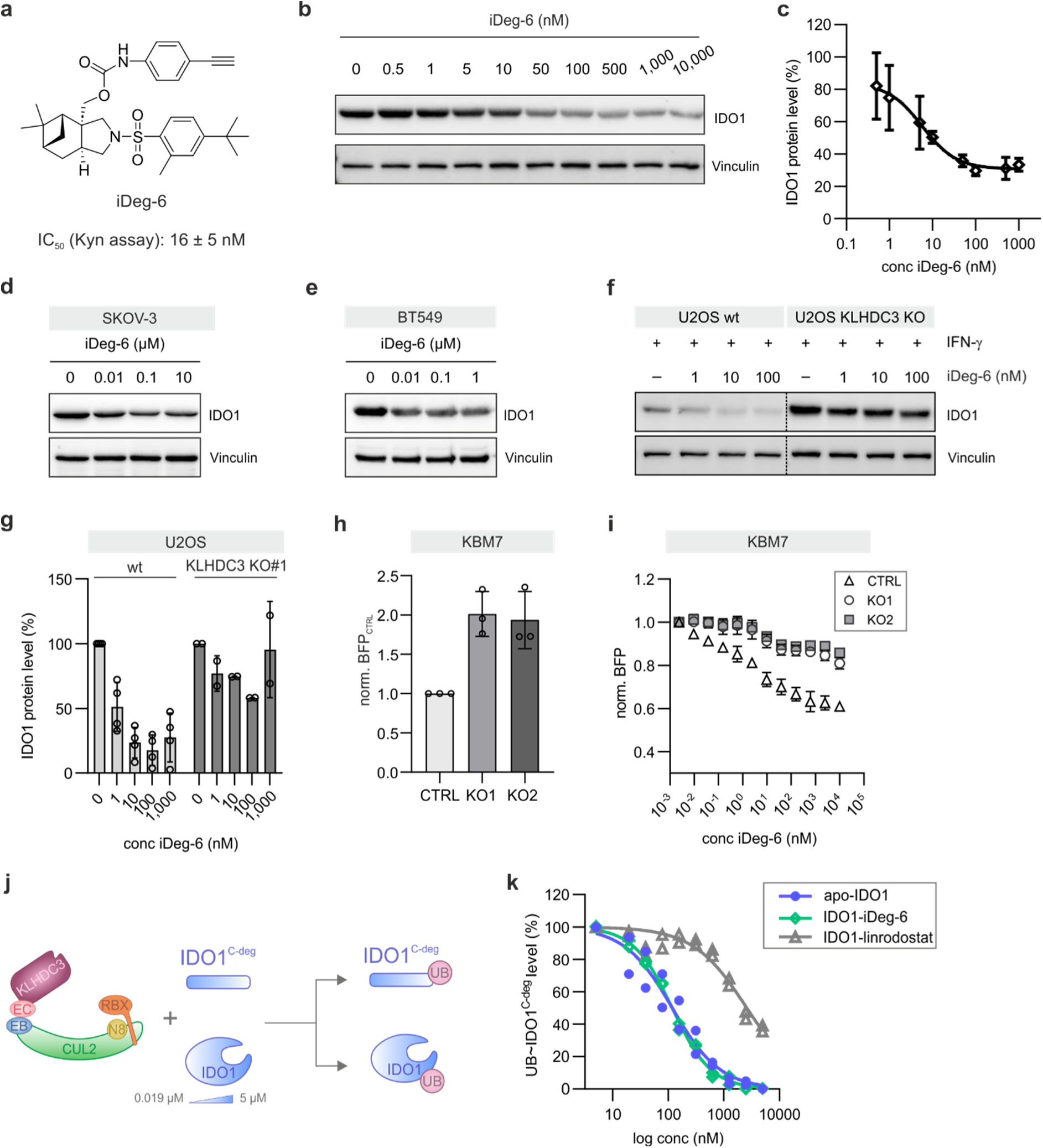
Validation of KLHDC3 as E3 ligase regulating IDO1. **a,** Structure of iDeg-6 and IC_50_ value in the Kyn assay in BxPC3 cells. Mean value ± SD, n = 3. **b,c,** Reduction of IDO1 protein levels by iDeg-6. Degradation efficiency was assessed in a modified setup including IFN-γ washout prior to compound addition. BxPC3 cells were treated with 50 ng/mL IFN-γ for 24 h prior to washout, addition of iDeg-6 for 24 h and immunoblotting (b). **c,** Quantification of IDO1 protein levels from b (mean values ± SD, n = 4 except for 50 and 500 nM (n = 3)). **d,e,** IDO1 levels in SKOV-3 (d) and BT549 (e) cells. Cells were treated with iDeg-6 for 24 h prior to immunoblotting. **f,g,** IDO1 protein levels in wildtype (wt) U2OS or KLHDC3 knockout (KO) U2OS cells. Cells were stimulated with 5 ng/mL IFN-γ for 24 h prior to washout, treatment with iDeg-6 or DMSO for 24 h and immunoblotting (f). Representative data of n = 4 (U2OS wt) or n = 2 (KO cells). **g,** Quantification of band intensities from f and Extended Data Fig. 9n. Mean values ± SD, n = 3 or n = 2. **h,i,** IDO1 protein levels in KBM7-BFP-IDO1 (CTRL) or KBM7-BFP-IDO1 KLHDC3 KO1 or KO2 cells in absence (h, normalized to CTRL) or presence of iDeg-6 (i, normalized to respective genotype). Mean values ± SD, n = 3. **j,** Schematic representation of the IDO1 ubiquitination competition assay. EB: elongin B; EC: elongin C; N8: neddylation; N8: NEDD8; UB: ubiquitin. **k,** Quantification of competition ubiquitination assays monitoring the ability of increasing concentrations of full-length apo-IDO1, iDeg-6-IDO1, or linrodostat-bound IDO1 to inhibit UBE2R2 mediated ubiquitination of the IDO1 C-terminal peptide (IDO1^C-deg^) by CRL2^KLHDC3^ (n = 2).

iDeg-6 depleted IDO1 protein also in SKOV-3 and BT549 cells in the absence of IFN-γ in a dose- and time-dependent manner (Fig. 5d-e and Extended Data Fig. 9g-k). The neddylation inhibitor MLN4924 rescued iDeg-6-induced IDO1 depletion (Extended Data Fig. 9l) and increased IDO1 in the absence of iDegs, demonstrating that neddylation is required for both native and iDeg-induced IDO1 degradation (Extended Data Fig. 9l). Knockout (KO) of KLHDC3 in U2OS or KBM7-BFP-IDO1 cells increased IDO1 amounts and rescued iDeg-6-dependent degradation (Fig. 5f-i and Extended Data Fig. 9m-n, see also Extended Data Fig S6e for iDeg-1, 2 and 3). IDO1 ubiquitination in presence of iDeg-6 was completely abolished for KLHDC3 variants carrying mutations in the degron recognition motif (R240A, S241E, R290A)^52^ (Extended Data Fig. 10a).

Surprisingly, *in vitro*, apo-IDO1 was rapidly ubiquitinated by CRL2^KLHDC3^. IDO1 was also ubiquitinated upon treatment of apo-IDO1 with iDeg-6 (Extended Data Fig. 10b,c). In contrast, both heme and apo-IDO1 inhibitor linrodostat rather suppressed IDO1 ubiquitination under these conditions (Extended Data Fig. 10b,c). Hence, in a biochemical setting, apo-IDO1 is a better substrate for KLHDC3 than holo-IDO1, which is consistent with the structural data showing heme or lindrodostat stabilizing the arrangement of the K-helix. We obtained quantitative insights into KLHDC3 binding to the various versions of IDO1 by performing competitive ubiquitination assays. Briefly, ubiquitination of IDO1 C-terminal peptide (IDO1^C-deg^) was examined while titrating increasing concentrations of IDO1. If a full-length IDO1 binds, then peptide ubiquitination is inhibited, and a shift of substrate is observed. Thus, these assays provide a metric for substrate accessibility, and can be performed at the pure protein concentrations we could be achieve. Apo-IDO1 and iDeg6-bound IDO1 show IC50 values of 119 nM and 118 nM, respectively (Figure 5k,j and Extended Data Fig. 10d). In contrast, the IC50 value in the presence of IDO1-bound to linrodostat was more than 20 times higher (2.6 µM). Thus, apo-IDO1 and iDeg6-bound IDO1 are much better KLHDC3 substrates, whereas linrodostat binding renders IDO1 less suitable for ubiquitination.

These findings suggest a possible model for the regulation of IDO1: Freshly translated apo-IDO1 is rapidly cleared in cells by degradation mediated by KLHDC. In the presence of heme, apo-IDO1 is charged with heme to form holo-IDO1 and the holo-pool may thus escape degradation. Support for such a mechanism in cells (Fig. 6a) was obtained by treatment with heme synthesis inhibitor succinyl acetone to deplete heme and shift the equilibrium to the apo-IDO1 form. In IFN-γ-stimulated BxPC3 cells, succinyl acetone reduced IDO1 protein levels and addition of hemin dose-dependently increased IDO1 (Fig. 6b-c and Extended Data Fig. 10e-f).

**Fig. 6.**
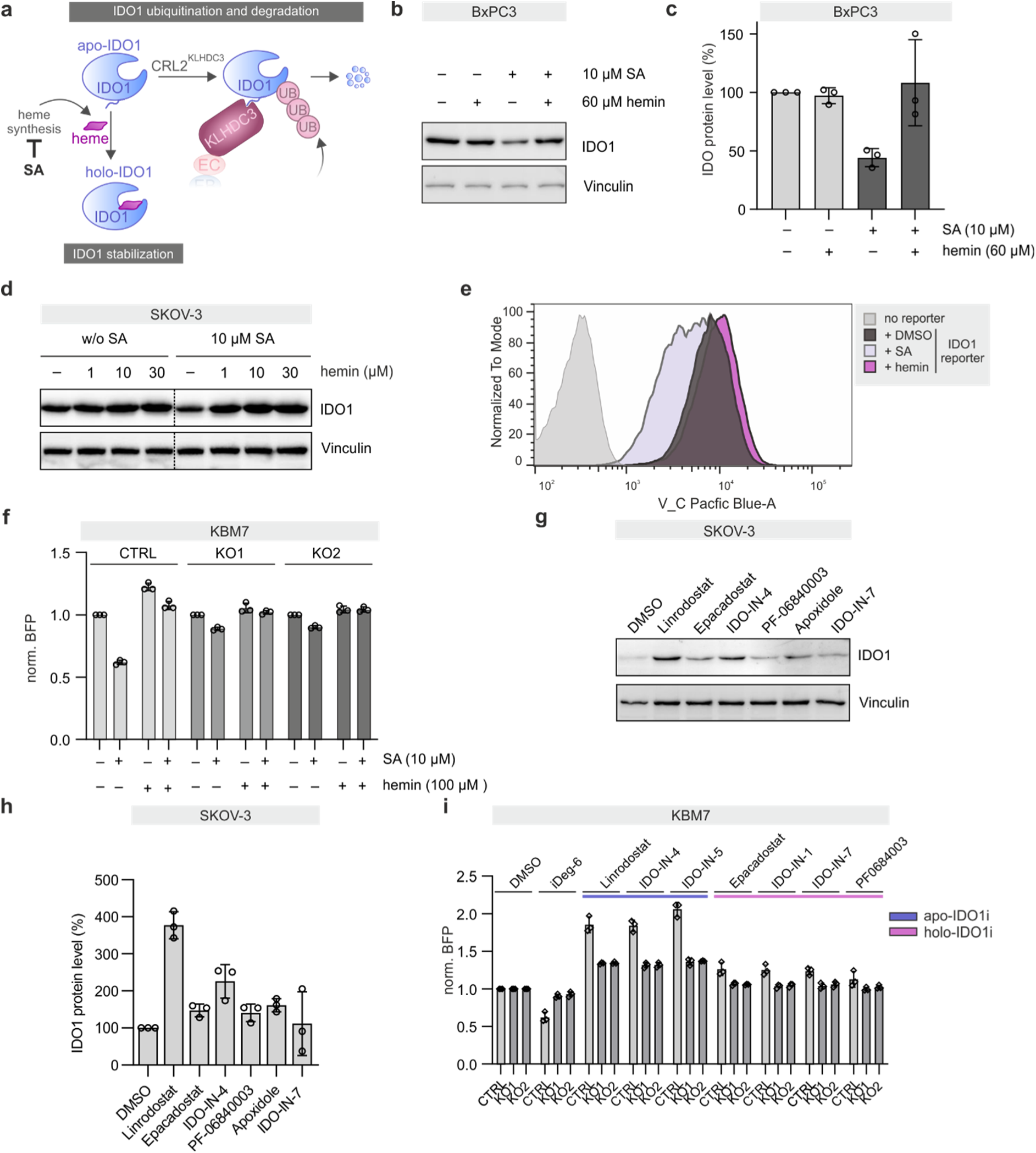
Apo-IDO and not holo-IDO1 is the preferential substrate for degradation. **a,** Regulation of IDO1 by heme synthesis and succinyl acetone (SA) as an inhibitor of heme synthesis. **b,c,** Influence of SA and heme on IDO1 protein levels in BxPC3 cells as detected using immunoblotting. Cells were treated with 50 ng/mL IFN-γ with or without 10 µM SA for 24 h followed by the addition of hemin for further 24 h. **c,** Quantification of band intensities from b. Mean values ± SD, n = 3. **d,** Influence of SA and heme on IDO1 levels in SKOV-3 cells. Cells were treated with SA for 48 h prior to the addition of hemin for another 24 h and immunoblotting. See Extended Data Fig. 10c for quantified band intensities. **e,f,** Influence of SA and hemin on IDO1 abundance in KBM7-BFP-IDO1 (e) or KBM7-BFP-IDO1 KLHDC3 knockout cells (f). Cells were pre-treated with 10 µM SA for 24 h followed by a treatment for 48 h with 100 µM hemin or DMSO as a control. IDO1 levels were quantified using flow cytometry. Mean values ± SD, n = 3. **g,h,** Influence of known IDO1 inhibitors on IDO1 protein levels in SKOV-3 cells. Cells were incubated with the compounds (5 µM) for 48 h prior to immunoblotting (g). **h,** Quantification of band intensities from g. Mean values ± SD (n = 3). **i,** Influence of IDO1 inhibitors (IDO1i, 1 µM) on IDO1 protein levels in KBM7-BFP-IDO1 or KBM7-BFP-IDO1 KLHDC3 knockout cells. Cells were treated with the compounds for 24 h followed by quantification of IDO1 using flow cytometry. Mean values ± SD, n = 3. Uncropped blots are shown in Extended Data Fig. 14.

Similar observations were made in SKOV-3 cells and the KBM7-BFP-IDO1 stability reporter cells (Fig. 6d-e and Extended Data Fig. 10g). Addition of only hemin dose-dependently increased IDO1 protein levels in SKOV-3 cells, which is in agreement with the majority of IDO1 being in the apo-form ^19^. Changes in the IDO1 protein level did not result from altered *IDO1* mRNA levels (Extended Data Fig. 10h). Knockout of KLHDC3 counteracted succinyl acetone-mediated IDO1 depletion, indicating that apo-IDO1 degradation is dependent on KLHDC3 (Fig. 6f). These results support involvement of KLHDC3 in the regulation of apo-IDO1 turnover, both under native conditions and in the presence of iDegs. The mechanism of action relies on further pushing IDO1 towards the degradation-sensitive state.

Finally we asked whether other IDO1 inhibitors might control IDO1 abundance. Apo-IDO1 inhibitors like linrodostat, IDO-IN-4 and IDO-IN-5 even increased IDO1 protein levels in cells, which was partially reverted in KLHDC3 knockout cells (Fig. 6g-i). These findings are in agreement with the biochemical data showing lower ubiquitination of IDO1 in the presence of linrodostat (Fig. 5l), and our structural data showing iDegs bind a distinct conformation of IDO1. Hence, the IDO1-KLHDC3 degradation circuit can be modified in both directions by liganding common residues albeit with distinct overall structural consequences.

## Discussion

Clinical investigation of IDO1 inhibition has had limited success thus far ^46,54^, possibly due to its non-enzymatic signaling function^29,32^. This limitation could potentially be addressed by depleting IDO1 by targeted protein degradation, which requires an IDO1 degrader strategy. We identified a pseudo-natural product class termed iDegs that potently induce IDO1 ubiquitination and degradation mediated by the E3 ligase CRL2^KLHDC3^. Notably, KLHDC3 mediates native IDO1 ubiquitination, thus, iDegs exploit the native pathway to IDO1 degradation. This unique mechanism is favored by the inherent conformational plasticity of the enzyme that exists in an apo and heme-bound state, and the activity of the enzyme depends on heme availability. iDegs bind in the heme pocket, and, therefore, can be considered apo-IDO1 inhibitors. Only the most potent compounds iDeg-2, -3 and -6 inhibited *in vitro* the enzymatic formation of Kyn, which reflects the efficiency for heme displacement *in vitro*. However, in cells IDO1 is expressed as apo-IDO1 and heme binding to IDO1 is reversible. iDegs bind to apo-IDO1 that is either present in the cells or generated by intrinsic heme loss, leading to loss of enzymatic activity. More importantly, iDegs induce a conformational shift of the C-terminal helix and lock IDO1 in a state favoring ubiquitination by KLHDC3. As degraders, iDegs are exceptional apo-IDO1 inhibitors since other apo-IDO1 modulators rather increase IDO1 protein abundance. Our data demonstrate that apo-IDO1 but not heme-bound IDO1 is preferentially ubiquitinated and degraded, suggesting that apo-IDO1 as well can readily adopt a ubiquitination-sensitive conformation. In contrast, heme binding prevents IDO1 turnover. Therefore, targeting the same binding pocket by the heme cofactor or small-molecule ligands has completely opposite impact on the IDO1-KLHDC3 degradation circuit and IDO1 fate.

Thus far, degraders employing the native degradation mechanisms have largely been overlooked and only few examples, i.e., targeting BCL6 or EZH2^55,56^, support the notion that monovalent degraders in a more general sense may regulate the native mechanisms of protein homeostasis for the respective targets^2^. Moreover, Scholes et al. identified supercharging of endogenous degradation pathways as a frequently observed mechanism of action of kinase degradation that is induced by kinase inhibitors.^57^ Mechanistically, kinase inhibitors can prompt degradation by changing activity, localization or aggregation state of kinases. Functionally differentiated, iDegs represent a first account of small molecules that act as a switch to induce a conformational state that shifts the equilibrium to the degradation-sensitive state, thereby channeling the IDO1-iDeg complex to the native degradation mechanism. This new route to pharmacologic protein degradation complements current degradation strategies that are based on PROTACs or MGDs and may also apply to other proteins.

The dual mechanism of action distinguishes iDegs since protein inhibition and depletion will impair all IDO1 functions, i.e., including the Kyn-independent signaling activity of IDO1. In addition, the increase of IDO1 protein by apo-IDO1 inhibitors is expected to diminish their efficacy. The failure of IDO1 inhibitors in the clinic may be due to its non-enzymatic signaling function that cannot be antagonized by inhibitors only, and to the fact that IDO1 inhibitors may upregulate IDO1 protein^32,38^. In contrast, IDO1 degradation would eliminate both, enzymatic activity and signaling function, and may open up new opportunities for the treatment of cancer or, as recent studies imply, the treatment of diseases related to Epstein Barr virus infections^13^.

## Acknowledgements

Research at the Max Planck Institute of Molecular Physiology and Max Planck Institute of Biochemistry was supported by the Max Planck Society. We acknowledge the European Synchrotron Radiation Facility (ESRF) for provision of synchrotron radiation facilities under proposal number MX-2391 and we would like to thank the local beamline support scientists for assistance and support in using beamline ID30B. We acknowledge Elizabeth D. Arnold, Shondra M. Pruett-Miller and the St. Jude Center for Advanced Genome Engineering (CAGE) for generating the U2OS-KLHDC3 knockout cells and St. Jude Children’s Research Hospital, ALSCAC, and NIH P30 CA021765 as providing funding for the CAGE and work performed by D.C.S. together with B.A.S.. E.H acknowledges the International Max-Planck Research School for a doctoral scholarship. The compound management and screening center (COMAS) in Dortmund is acknowledged for performing the high-throughput screening. We thank Malte Metz, Andreas Brockmeyer und Walburga Hecker for the mass spectrometry measurements and Christine Nowak, Christiane Pfaff, Jens Warmers and Philipp Lampe for experimental support. We thank the Core Facility Flow Cytometry of the Medical University of Vienna for access to flow cytometry instruments and assistance with flow cytometric cell sorting as well as the CeMM Biomedical Sequencing Facility for NGS sample processing, sequencing, and data curation. The research leading to these results also received support from the Innovative Medicines Initiative Joint Undertaking under Grant No. 115489, resources of which are composed of financial contribution from the European Union’s Seventh Framework Programme (FP7/2007–2013) and the EFPIA (The European Federation of Pharmaceutical Industries and Associations) companies’ in-kind contribution. The project is also receiving funding from the programme “Netzwerke 2021”, an initiative of the Ministry of Culture and Science of the State of Northrhine Westphalia. CeMM and the Winter lab are supported by the Austrian Academy of Sciences. The Winter lab is further supported by funding from the European Research Council (ERC) under the European Union’s Horizon 2020 research and innovation program (grant agreement 851478), as well as by funding from the Austrian Science Fund (FWF, projects P7909, P36746 and P5918723). M.L acknowledges research support by grant PID2021-122611NB-100 funded by the Agencia Estatal de Investigación of Ministerio de Ciencia e Innovación (MCIN/AEI/ 10.13039/501100011033). N.S.S. is further supported by the FWF postdoctoral Esprit fellowship ESP 426 and Marie Skłodowska-Curie postdoctoral fellowship (grant agreement number: 101029199).

## Author contribution

H.W. and S.Z. designed the project. E.H., B.L., N.S.S., X.-F.C, A.B., D.C.S., A.B., K.R., L-M.P., L.D., and T.T.X. and P.J.M. performed the biological research. X.-F.C., M.B. and K.R.N. synthesized the compounds. E.H., B.L., X.-F.C., N.S.S., analyzed the data. H.I. quantified the CRISPR/Cas9 screen sequencing data. B.L., R.G. and M.L. solved the crystal structure. S.S. adapted and performed the screening assays. K.K., M.G., P.J.M., B.A.S. and G.E.W supervised project parts. E.H., B.L., S.Z., and H.W. wrote the text. All authors discussed the results and commented on the manuscript.

