## Supplemental data file for "Identification of a Monovalent Pseudo-Natural Product Degrader Class Supercharging Degradation of IDO1 by its native E3 KLHDC3" (from another version of this preprint: version 2, 2024-12-19)

### Supplementary Information

### Extended Data Figures

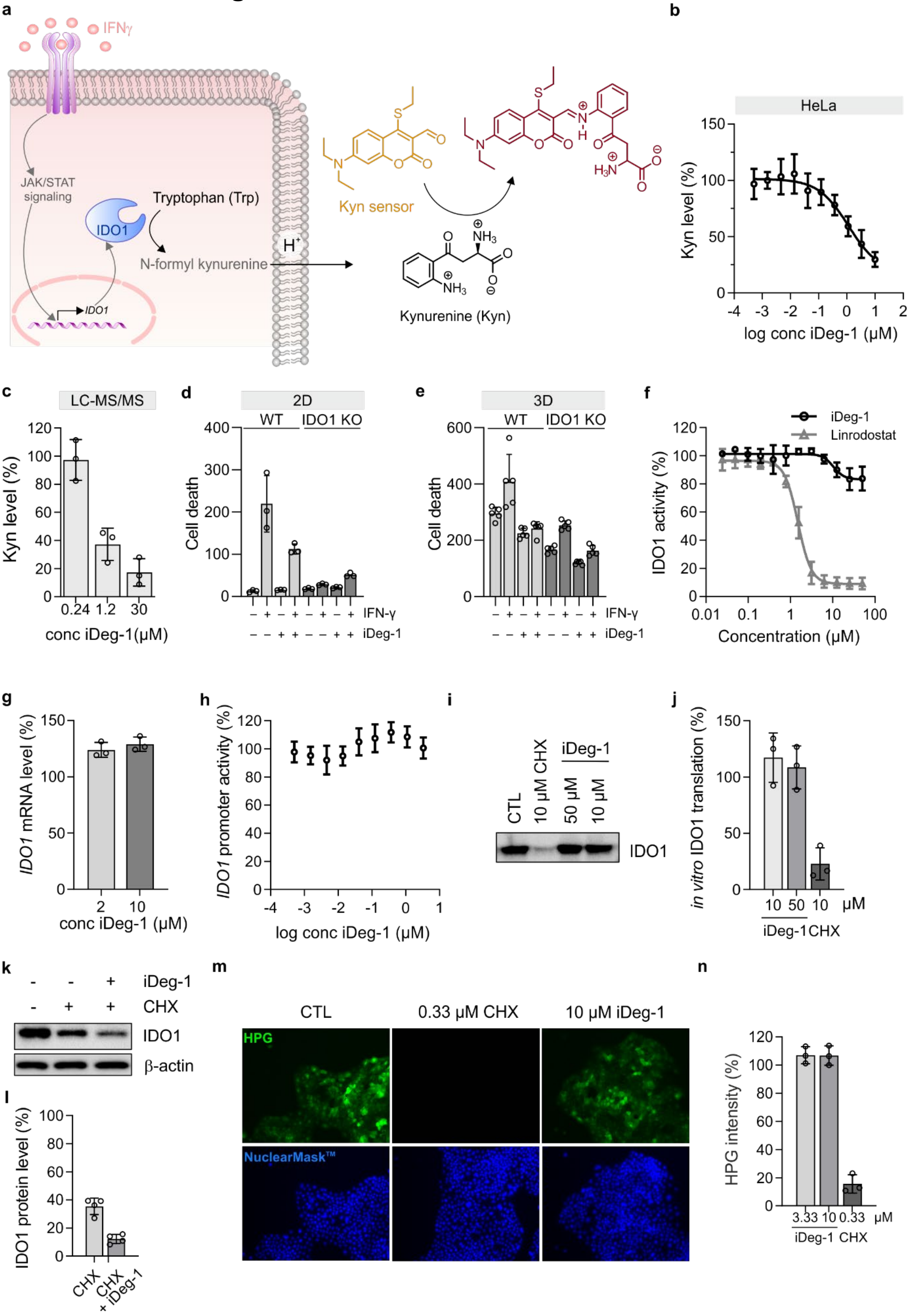

**Extended Data Fig. 1: Validation of iDeg-1 as a small molecule reducing IDO1 protein levels. a,** Schematic overview of the Kyn screening assay. BxPC3 cells are stimulated with IFN-γ to induce IDO1 expression via the JAK/STAT signaling pathway. After 48IFN-γ, L-Trp and the compounds, cells are hydrolysed and Kyn levels are quantified in the cell supernatant by means of the Kyn sensor. Fluorescence intensity (ex/em 535/595 nm) is proportional to the Kyn amount generated by the cells. **b,** Kyn assay in HeLa cells treated with IFN-γ, Trp and iDeg-1 for 48 h prior Kyn level detection utilizing *p*-DMAB. Mean values ± SD, n = 3. **c,** Kyn levels in BxPC3 cells treated with IFN-γ, Trp and iDeg-1 for 48 h. Kyn levels were detected using LC-MS/MS. Mean values ± SD, n = 3. **d,e,** Influence of iDeg-1 (5 µM) on cell death in SKOV-3 cells grown in 2D (d) or 3D (e). Cell death was detected by means of CellTox Green in presence or absence of IFN-γ (1 ng/mL). Mean values ± SD for N = 3 after 96 h (d) or N = 5 after 6 days (e). Representative data of n = 3. **f,** Influence of iDeg-1 on the *in vitro* IDO1 enzymatic activity. The apo-IDO1 inhibitior linrodostat was used as a control. Mean values ± SD, n = 3. **g,** *IDO1* mRNA expression levels. BxPC3 cells were treated with IFN-γ and iDeg-1 for 24 h prior to isolation of RNA, RT-qPCR. Mean values ± SD, n = 3. **h,** Influence on the *IDO1* promoter activity. HEK293T cells were transfected with a construct encoding firefly luciferase (Fluc) under the control of the *IDO1* promoter and a plasmid for constitutive expression of *Renilla* luciferase (Rluc). Cells were treated with IFN-γ and the compounds for 24 h. Fluc/Rluc ratios were normalized to samples that were treated with IFN-γ only (set to 100%). Mean values ± SD, n = 3. **i**,**j,** *In vitro* IDO1 protein translation. IDO1 translation was analyzed in cell lysates in the presence of iDeg-1, cycloheximide (CHX) or DMSO (CTL). Representative immunoblot (n = 3 or n = 4) (i) and quantified band intensities (j). **k, l,** CHX chase assay. BxPC3 cells were treated with IFN-γ for 24 h prior to addition of 5 µM CHX or 5 µM CHX and 10 µM iDeg-1. Representative immunoblots (n = 4) (k). (l) Normalized band intensities from k. **m, n,** Analysis of global protein translation in cells 24 h after compound treatment by means of HPG detection. Representative images of n = 3 of HPG intensities displaying protein translation (green) and nuclear stain (blue). **n,** Normalized HPG signal intensities of the images in m. Mean values ± SD, n = 3. Uncropped blots are shown in Extended Data Fig. 11.

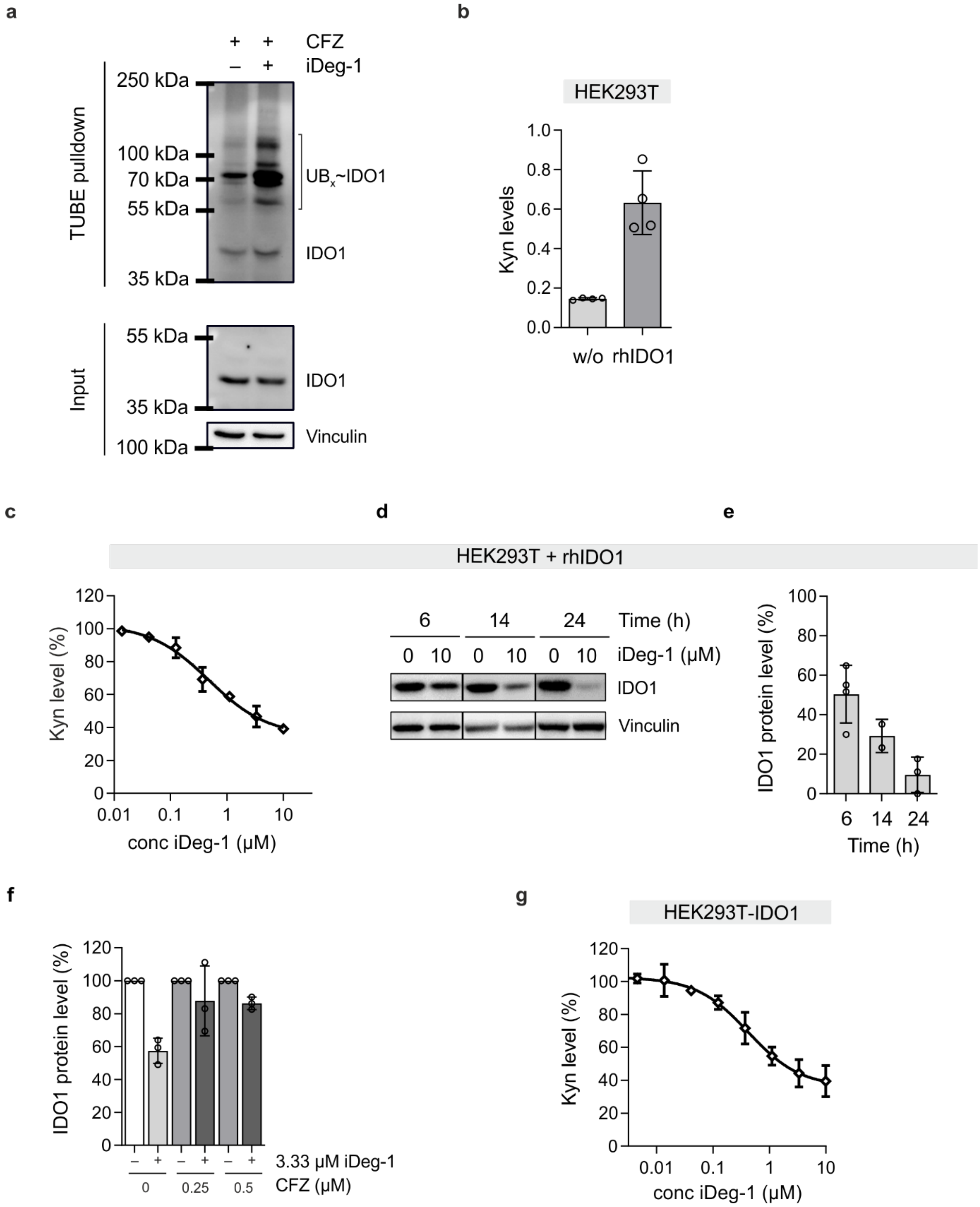

**Extended Data Fig. 2: iDeg-1 inhibits Kyn production in cells by IDO1 degradation. a,** TUBE pulldown after treatment of IFN-γ-stimulated BxPC3 cells with iDeg-1 or DMSO. Cells were pre-treated for 1 h with the proteasome inhibitor carfilzomib (CFZ, 450 nM) followed by addition of 50 µM iDeg-1 for 4 h prior to lysate preparation and TUBE pulldown. Representative immunoblot (n = 3). UB: ubiquitin. **b,** HEK293T cells were electroporated with and without IDO1 and treated with Trp. 24 h later, Kyn levels were detected using *p*-DMAB. Mean values ± SD, n = 4. **c,** HEK$^{rhIDO1}$ cells treated with Trp and iDeg-1 for 24 h prior Kyn level detection using *p*-DMAB. Mean values ± SD, n = 3. **d,e,** HEK239T cells were electroporated with rhIDO1 protein followed by treatment with iDeg-1 (10 µM) for 6, 14 or 24 h. IDO1 protein levels were determined via immunoblotting (d). **e,** Normalized band intensities from d. Mean values ± SD, n = 2 (14 h), n = 3 (24 h), n = 4 (6 h). **f,** Normalized data from quantified band intensities from Fig. 1g. The effect of iDeg-1 in co-treatment conditions is represented relative to the treatment of carfilzomib (CFZ) alone (set to 100 %). Mean values ± SD, n = 3. **g,** HEK293T cells were transfected with an IDO1 expression plasmid 24 h prior the addition of Trp and iDeg-1. Kyn levels were determined 24 h after compound addition using *p*-DMAB. Mean values ± SD, n = 3. Uncropped blots are shown in Extended Data Fig. 11.

a

1 MAHAMENSWT ISKEYHIDEE VGFALPNPQE NLPDFYNDWM FIAKHLPDLI ESGQLRERVE KLNMLSIDHL

71 TDHKSQRLAR LVLGCITMAY VWGKGHGDVR KVLPRNIAVP YCQLSKKLEL PPILVYADCV LANWKKKDPN

141 KPLTYENMDV LFSFRDGDCS KGFFLVSLLV EIAAASAIKV IPTVFKAMQM QERDTLLKAL LEIASCLEKA

211 LQVFHQIHDH VNPKAFFSVL RIYLSGWKGN PQLSDGLVYE GFWEDPKEFA GGSAGQSSVF QCFDVLLGIQ

281 QTAGGGHAAQ FLQDMRRYMP PAHRNFLCSL ESNPSVREFV LSKGDAGLRE AYDACVKALV SLRSYHLQIV

Ub(K389)

361 TKYILIPASQ QPKENKTSED PSKLEAK[GTG GTDLMNFLKT VR]STTEKSLL KEG

b

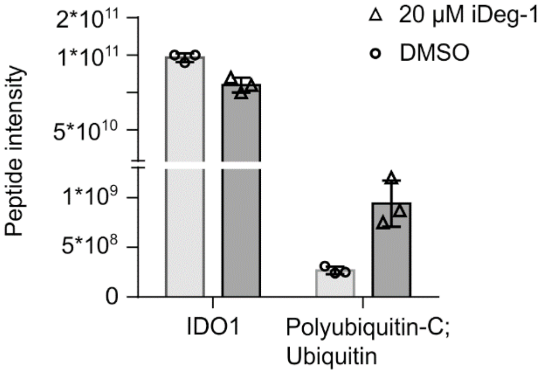

c

1 MQIFVKTLTG KTITLEVEPS DTIENVKAKI

Ub(K48)

31 QDKEGIPPDQ QR[LIFAGKQL EDGR]TLSDYN

61 IQKESTLHLV LRLRGG

**Extended Data Fig. 3: iDeg-1 induces polyubiquitination of lysine 389 of IDO1**. IFN-γ-stimulated BxPC3 cells were treated with iDeg-1 or DMSO as a control for 6 h prior to lysate preparation. IDO1 was immunoprecipitated and the enriched fraction was digested and analyzed by MS. **a,** IDO1 protein sequence. Amino acids in bold illustrate regions identified in the MS experiment. K389: the identified ubiquitination site of IDO1. Due to incomplete sequence coverage in the MS experiment, ubiquitination of additional lysines cannot be excluded **b,** Measured relative peptide intensity for IDO1 and polyubiquitin-C. Mean values ± SD, n = 3. **c,** Ubiquitin protein sequence. Identified peptides are displayed in bold. The identified ubiquitination site K48 is indicated. K6, K11, K29 and K33 in ubiquitin could not be identified due to incomplete sequence coverage in the MS experiment.

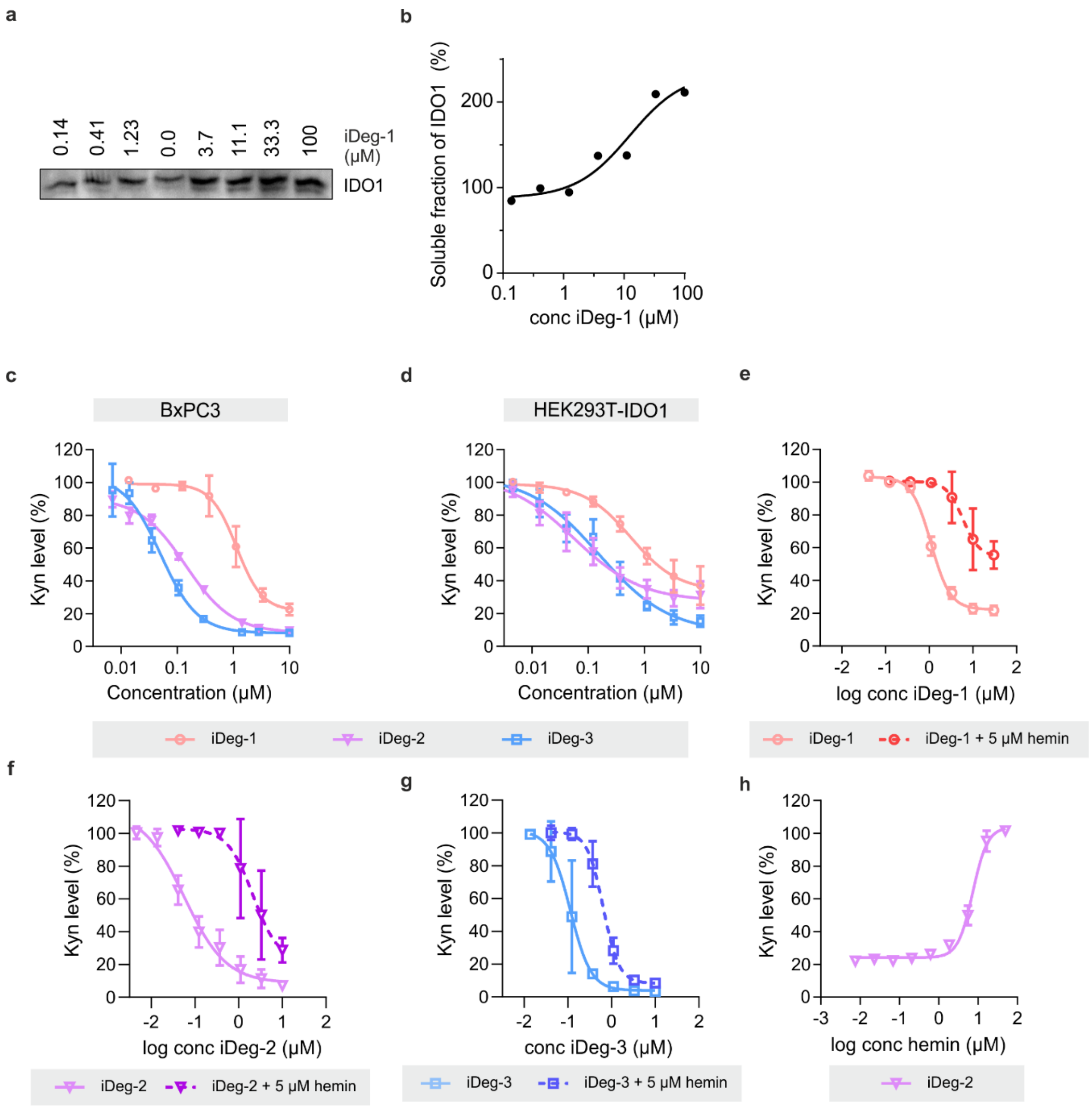

**Extended Data Fig. 4: iDegs target IDO1. Validation of iDeg-2 and iDeg-3 as IDO1 degraders. a, b,** Isothermal CETSA using iDeg-1. SKOV-3 cells were treated with iDeg-1 or DMSO as a control for 1 h at 37°C prior to heat treatment at 50°C, lysis and analysis of the soluble fraction via immunoblotting. Representative immunoblot for IDO1 (a) and respective melting curve (b). **c,** Kyn assay in BxPC3 cells treated with iDegs, Trp and IFN-γ for 48 h prior to detection of Kyn levels utilizing *p*-DMAB. Mean values ± SD, n = 3. **d,** HEK293T cells were transiently transfected with pCMV3-IDO1 24 h prior to compound addition. Another 24 h later, Kyn levels were detected utilizing *p*-DMAB. Mean values ± SD, n = 3. **e-g,** Influence of hemin on iDeg activity. Kyn assay in IFN-γ-stimulated BxPC3 cells for iDeg-1 (e), iDeg-2 (f) or iDeg-3 (g) in the presence or absence of 5 µM hemin. Kyn levels were detected after 48 h using p-DMAB. Mean values ± SD, n = 3 for iDeg-1, 3 and n ≥ 2 for iDeg-2. **h,** Dose-dependent influence of hemin on iDeg-2-mediated decrease of Kyn levels. IFN-γ–stimulated BxPC3 cells were incubated with 0.5 µM iDeg-2 and different concentrations of heme for 48 h prior to detection of Kyn levels with p-DMAB. Mean values ± SD, n = 3. Uncropped blots are shown in Extended Data Fig. 12.

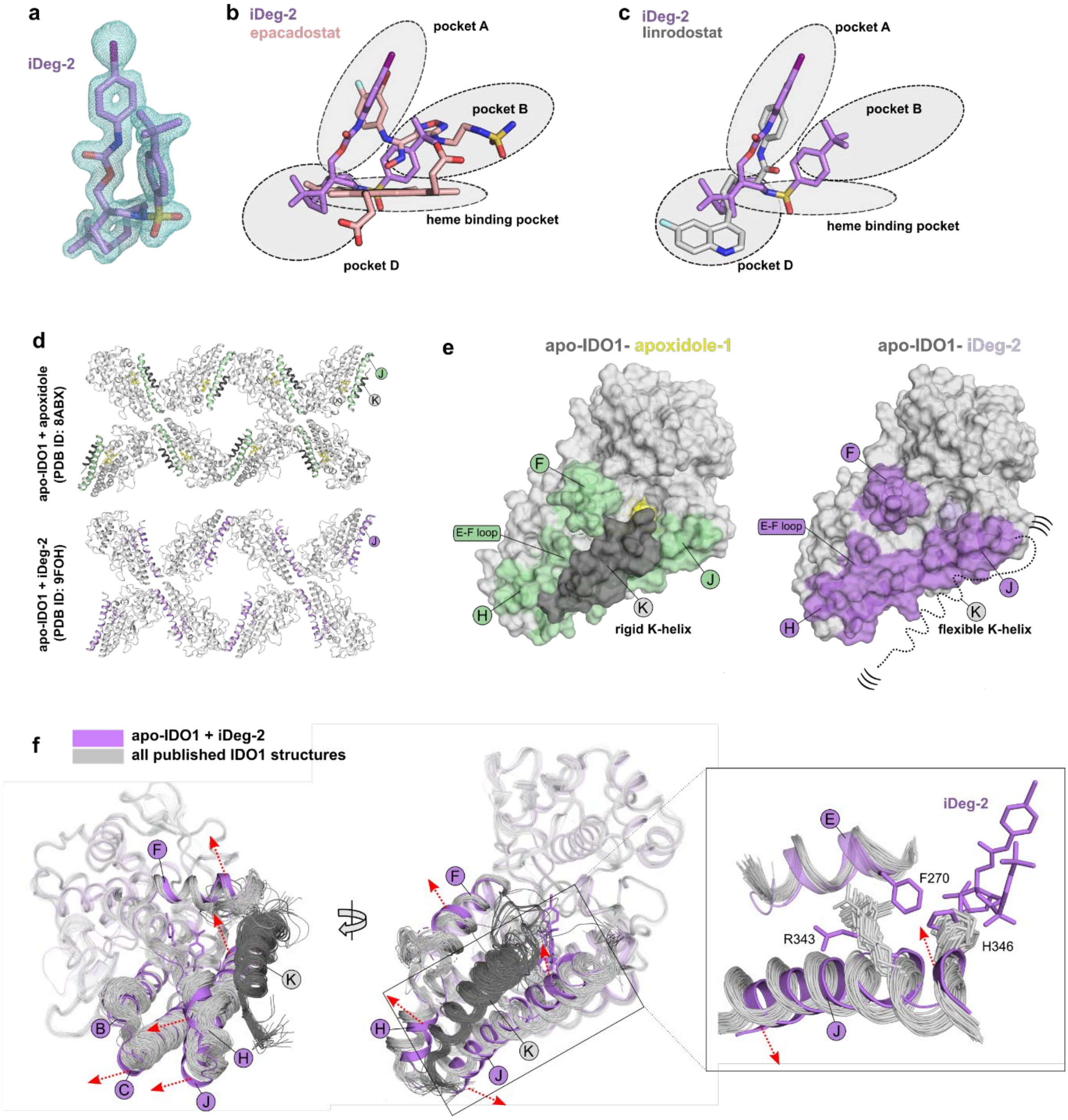

**Extended Data Fig. 5: Co-crystal structure of IDO1 with iDeg-2. a,** Electron density map (2Fo-Fc) calculated with phases from the final refined model, contoured at 1.6σ, showing the refined iDeg-2 structure in stick representation. **b,** Overlay of iDeg-2 (violet) with the holo-IDO1 inhibitor epacadostat in complex with heme (PDB ID: 5wn8, pink). **c,** Overlay of iDeg-2 with the apo-IDO1 inhibitor linrodostat (PDB ID: 6DPRB, gray). **d,** The highly similar crystal lattice packing of apo-IDO1 + iDeg-2 and apo-IDO1 + apoxidole, indicates that the absence of electron density for the K-helix (dark gray) in the iDeg-2 structure is not an artifact of crystal packing. The J-helix is highlighted in green (8ABX) or purple (9FOH). Apoxidole and iDeg-2 are shown in yellow and purple, respectively. **e,** Comparison of the apo-IDO1 structures with apoxidole (PDB ID: 8ABX) and iDeg-2, illustrating the transition of the K-helix from a rigid to a dynamic conformation upon iDeg-2 binding. Interacting surface regions with the K-helix are highlighted. **f,** Overlay of the apo-IDO1-iDeg-2 structure with all published IDO1 structures, excluding seven with high coordinate uncertainty. Shown are the front view, a 90º rotated side view, and a zoom-in view of the E- and J-helices, with the positions of H346 and R343 indicated. PDB IDs included: 7B1O, 7AH5, 7AH6, 7AH4, 6KOF, 8I7L, 6O3I, 8FUR, 7A62, 6MQ6A, 6MQ6B, 7P0N, 7P0R, 7M7D, 7NGE, 6F0A, 6R63, 7E0T, 5WN8, 5WHR, 6PU7, 5WMV, 5WMU, 5WMX, 5WMW, 6KPS, 6X5Y, 6WJY, 6UBP, 7E0P, 6PZ1, 7E0U, 7E0O, 7E0S, 6V52, 6WPE, 7M63, 6KW7, 2D0U, 2D0T, 6E41, 7E0Q, 4PK6, 5EK3, 6DPQA, 6DPQB, 6E43, 6E46, 4U74, 6CXV, 6AZW, 5EK2, 5EK4, 5ETW, 6E45, 6E44, 6CXU, 5XE1, 6E40, 6E35, 6DPRA, 6DPRB, 4PK5, 4U72, 6AZV, 6AZU, 6E42, 8ABX, 7ZV3, 7RRD, 7Z2L, 7RRB, 7RRC, 7YXT AND 8U5I.

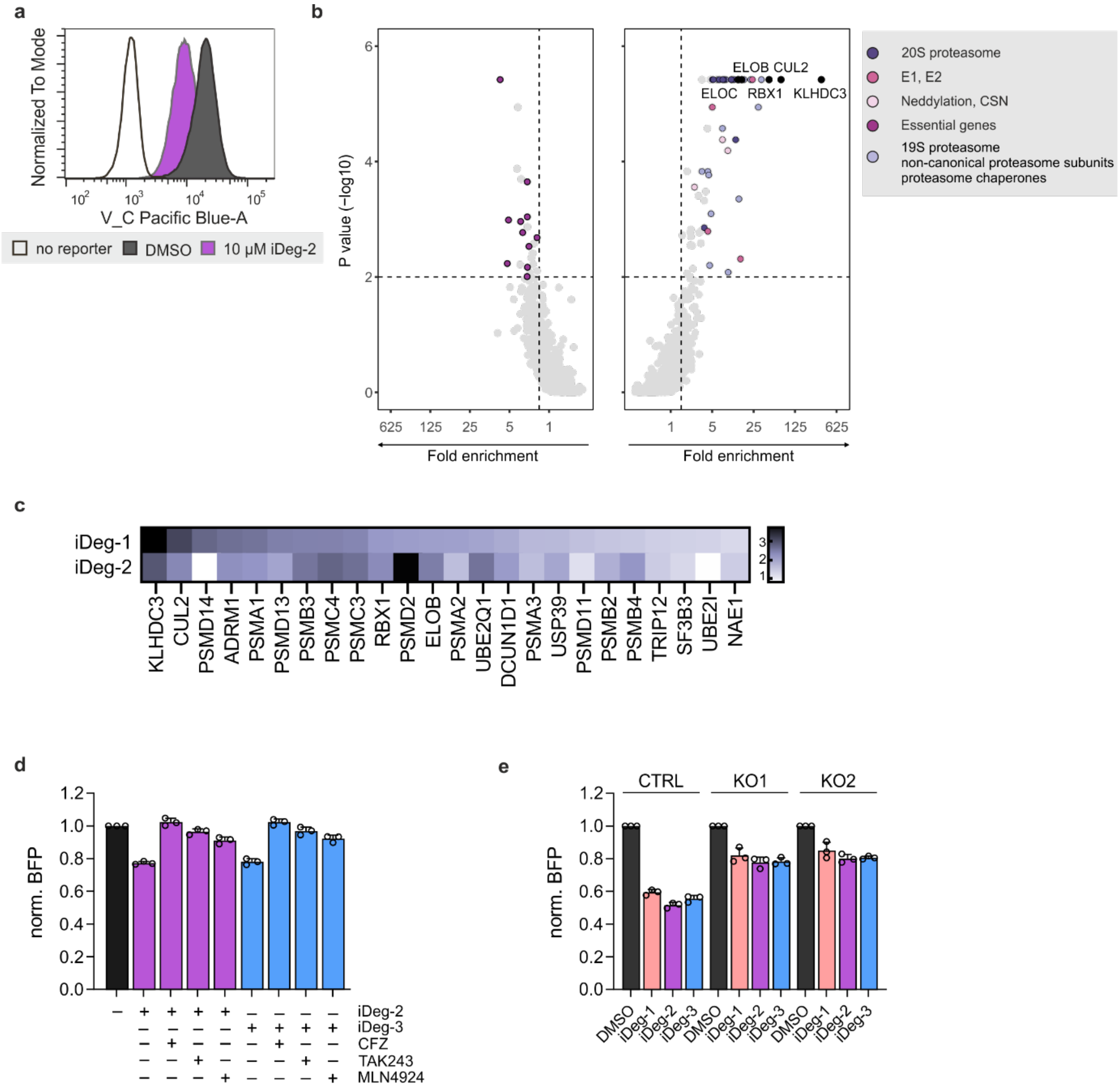

**Extended Data Fig. 6: iDeg-mediated IDO1 depletion is dependent on KLHDC3. a,** Representative flow cytometric histogram for iDeg-2 mediated depletion of BFP-IDO1 (24 h, 10 µM) in KBM7 iCas9 cells. **b,** Identification of genes required for iDeg-2-mediated IDO1 degradation. CSN: COP9 signalosome. **c,** Heatmap of the gene enrichment in the CRISPR screen calculated as fold-change between the DMSO control and iDeg-1 and iDeg-2 treatment. **d,** Detection of IDO1 levels using the IDO1 stability reporter. KBM7 IDO1 reporter cells were treated with iDeg-2 or 3 (10 µM) +/- indicated co-treatments for 10 h prior to detection of IDO1 levels using flow cytometry. CFZ: 1 µM; TAK243: 1 µM, MLN4324: 1 µM. Mean values ± SD, n = 3. **e,** IDO1 protein levels in KBM7-BFP-IDO1 CTRL or KBM7-BFP-IDO1 KLHDC3 KO1 or KO2 cells upon treatment with iDeg-1, 2 or 3 (1 µM) for 24 h. Values were normalized to the respective DMSO control, which was set to 1. Mean values ± SD, n = 3.

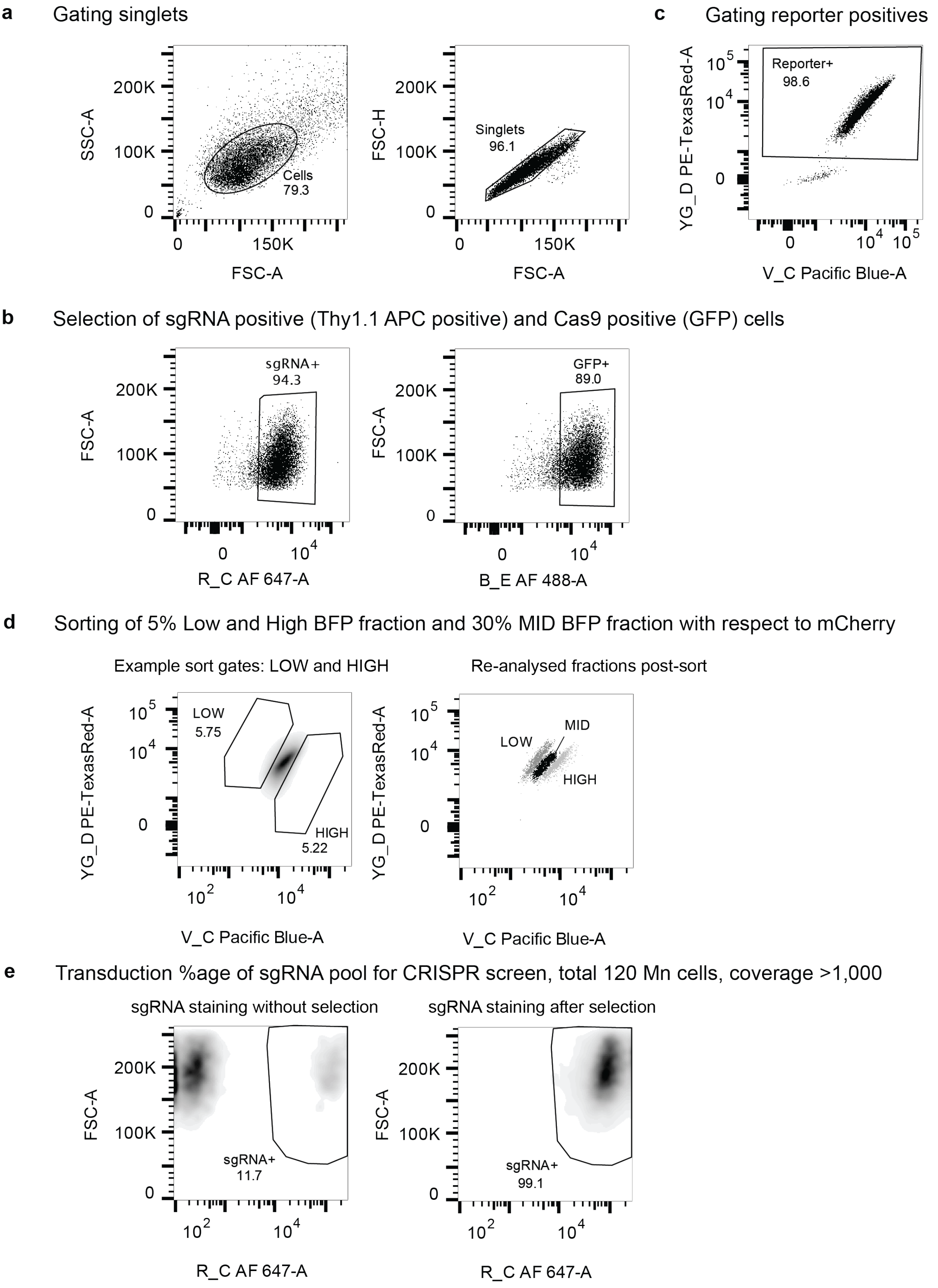
a
Gating singlets
SSC-A
200K
100K
0
Cells
79.3
150K
FSC-A
FSC-H
Singlets
96.1
c
Gating reporter positives
YG_D PE-TexasRed-A
$10^5$
$10^4$
Reporter+
98.6
V_C Pacific Blue-A
b
Selection of sgRNA positive (Thy1.1 APC positive) and Cas9 positive (GFP) cells
FSC-A
sgRNA+
94.3
R_C AF 647-A
GFP+
89.0
B_E AF 488-A
d
Sorting of 5% Low and High BFP fraction and 30% MID BFP fraction with respect to mCherry
Example sort gates: LOW and HIGH
Re-analysed fractions post-sort
LOW
5.75
HIGH
5.22
$10^2$
MID
e
Transduction %age of sgRNA pool for CRISPR screen, total 120 Mn cells, coverage >1,000
sgRNA staining without selection
sgRNA staining after selection
sgRNA+
11.7
sgRNA+
99.1

**Extended Data Fig. 7**. **Gating strategy for flow cytometric assays and FACS. a,** The identification of viable cells was performed using the forward and side scatter area (FSC-A and SSC-A, respectively). Singlets were selected using the FSC-A and FSC-H (height) as indicated. **b,** sgRNA transduction was assessed by staining for the Thy1.1 surface antigen for both single and pooled sgRNA experiments. GFP expression monitored Cas9 induction. **c,** Reporter-positive cells were selected in the BFP and mCherry channel. **d,** For the CRISPR-Cas9 screen the highest (HIGH) and lowest 5% (LOW) of BFP expressing cells were sorted and compared against the 30 % mid-fraction (MID). Re-analysis of sorted fractions was routinely performed to confirm sorting accuracy. **e,** The sgRNA pool for the CRISPR-Cas9 screen was transduced at 11.7 % for a total of 120 million cells and selected with G-418 (see Materials and Methods) prior to screening.

a

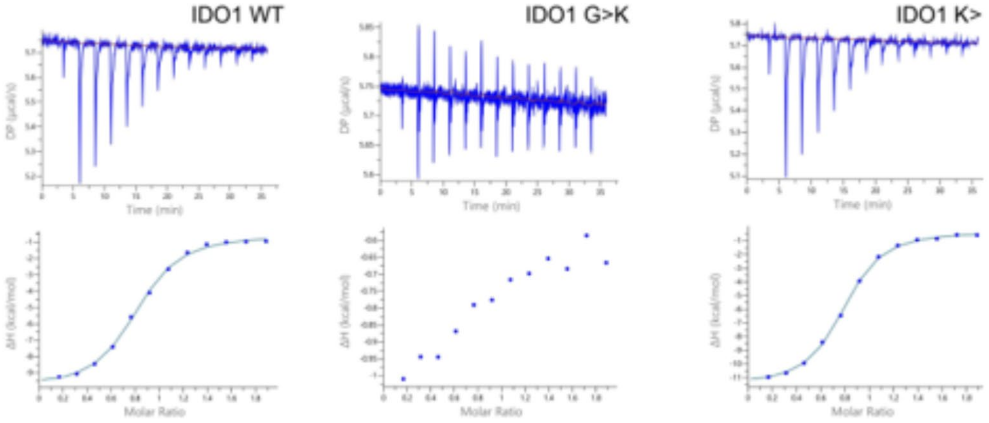

b

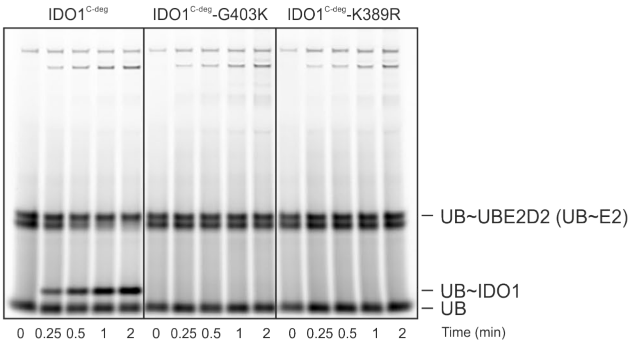

**Extended Data Fig. 8: Binding and ubiquitination of C-terminal IDO1 peptides. a,** Measurement of KLHDC3 binding to C-terminal IDO1 peptides using ITC. NB: no binding. **b,** Fluorescent scan of pulse-chase assay monitoring ubiquitination of C-terminal IDO1 peptides by UBE2D2. UBE2D2 was "pulse" loaded with fluorescent ubiquitin, and the resulting UBE2D2~UB thiolester was added to NEDD8~CRL2$^{KLHDC3}$ incubated with the indicated peptides. Samples were quenched with SDS sample buffer at the indicated timepoints. UB: ubiquitin.

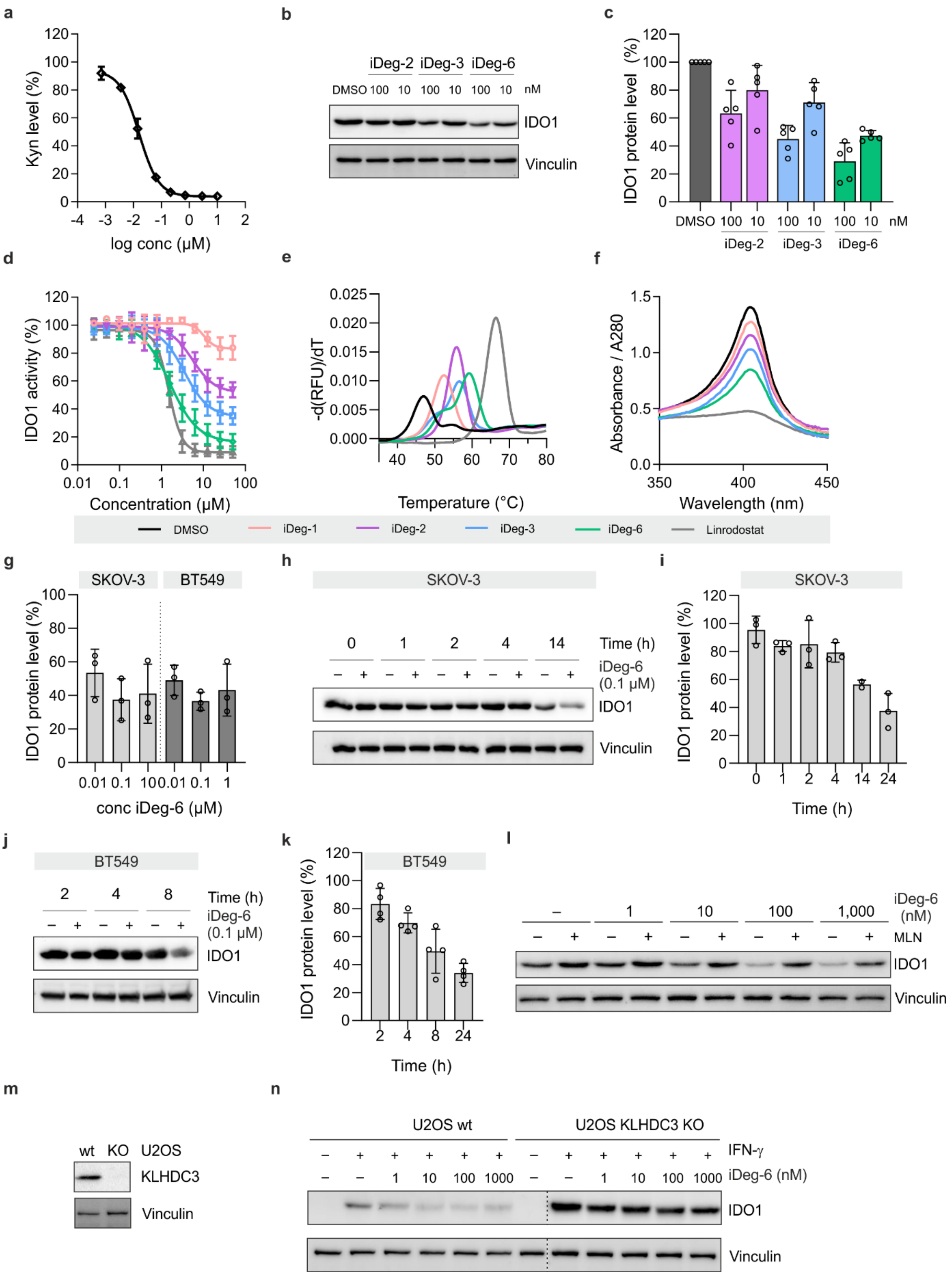

**Extended Data Fig. 9: Characterization of iDeg-6. a,** Kyn assay in BxPC3 cells after treatment with iDeg-6 and IFN-γ, respectively, for 48 h prior to detection of Kyn levels utilizing *p*-DMAB. Mean values ± SD, n = 3. **b,c,** IDO1 protein levels in BxPC3 cells upon treatment with IFN-γ for 24 h followed by washout and addition of iDeg-2, 3 or iDeg-6 for 24 h. Representative immunoblot (n = 5) for IDO1 and vinculin as a loading control. **c,** Quantified band intensities from b. Mean values ± SD, n = 5. **d,** *In vitro*

IDO1 enzymatic activity assay in presence of iDeg-1-6 or linrodostat. rhIDO1 was pre-incubated with the compounds at 37°C for 90 min prior to detection of Kyn levels using *p*-DMAB. Mean values ± SD, n = 3. **e,** rhIDO1 thermal stability in presence of 30 µM iDeg-1, 2, 3 or iDeg-6 (50 µM), or DMSO or linrodostat (50 µM) determined by means of nanoDSF. rhIDO1 and the compounds were pre-incubated for 3 h at 37°C prior to the measurement. Representative data of n = 3. **f,** Detection of heme-bound IDO1 by means of UV/Vis spectroscopy in presence of iDeg-6 (100 µM) as compared to iDeg-1, 2 and 3, DMSO or linrodostat (100 µM). Incubation at 37°C for 3 h. Representative data (n = 3). **g,** IDO1 protein levels in SKOV-3 and BT549 cells. Quantification of band intensities from Fig. 5d-e. Mean values ± SD, n = 3. **h-k,** IDO protein levels in SKOV-3 (h and i) or BT549 cells (j and k) after treatment with iDeg-6 for different time. IDO1 and vinculin were detected by means of immunoblotting. Quantification of band intensities from h and j is shown in i and k, respectively. Mean values ± SD, n = 3 except for 14 h (n = 2 for SKOV-3 and n = 4 for BT549). **l,** Influence of neddylation inhibition on IDO1 protein levels in BxPC3 cells. BxPC3 cells were stimulated with IFN-γ for 24 h prior to washout, treatment with 500 nM MLN4924 and iDeg-6 for 16 h and immunoblotting. Representative blot (n = 3). **m,** Detection of CRISPR-Cas9-mediated KLHDC3 knockout in U2OS cells. Levels of KLHDC3 and vinculin were detected by means of immunoblotting. Representative blot (n = 3). **n,** IDO1 protein levels in U2OS and U2OS cells carrying a KLHDC3 knockout. Representative blot (n = 3). Uncropped blots are shown in Extended Data Fig. 13.

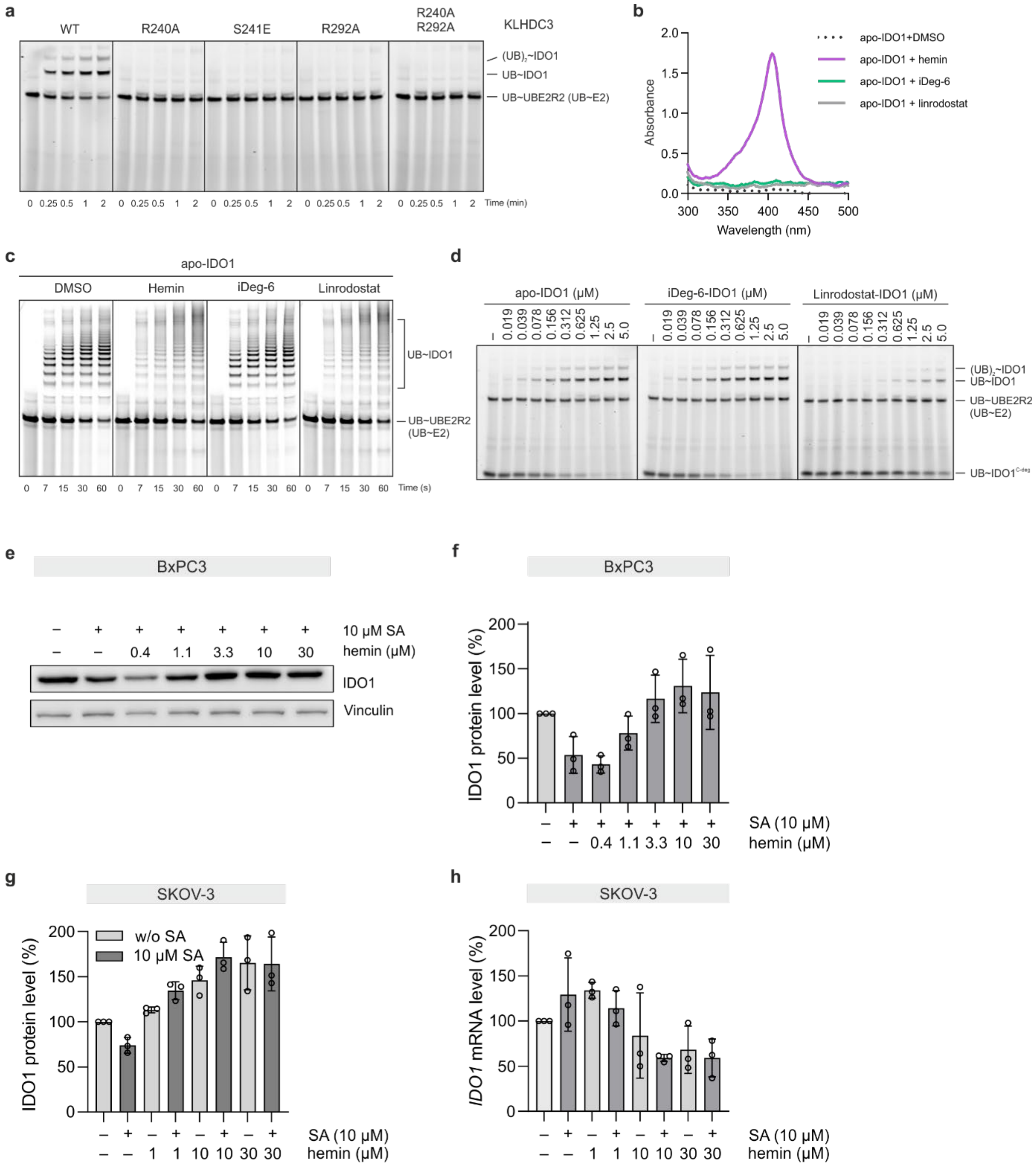

**Extended Data Fig. 10: Ubiquitination of IDO1 by CRL2$^{KLHDC3}$ and influence of succinyl acetone (SA) and hemin on IDO1 levels. a,** Fluorescent scan of pulse-chase assay monitoring ubiquitination of iDeg6-IDO1 by KLHDC3 mutants. iDeg6-IDO1 was incubated with CRL2$^{KLHDC3}$ E3 complex and ubiquitination reactions were initiated by the addition of the pre-formed thioester-linked E2~ubiquitin conjugate (UBE2R2~UB) with the ubiquitin fluorescently-labeled. **b,** Detection of heme load for the conditions in c detected by means of UV/Vis spectroscopy. **c,** Fluorescent scan of pulse-chase assay monitoring ubiquitination of apo-IDO1 with or without supplementation with small molecules by CRL2$^{KLHDC3}$ (b). Apo-IDO1 was incubated with the indicated compounds (30 µM) for 90 min at 37°C prior to incubation with CRL2$^{KLHDC3}$ E3 complex. Ubiquitination reactions were initiated by the addition of the pre-formed thioester-linked E2~ubiquitin conjugate (UBE2R2~UB) with the ubiquitin fluorescently-labeled. Representative data for n = 2. **e,f,** BxPC3 cells were treated with 50 ng/mL IFN-γ with or without 10 µM SA for 24 h followed by the addition of hemin for further 24 h. **f,** Quantification of band intensities from e. Mean values ± SD, n = 3. **g,** Quantification of band intensities from Figure 6d. SKOV-3 cells were treated with SA for 48 h prior to the addition of hemin for another 24 h and immunoblotting (see Fig. 6d). Mean values ± SD, n = 3. Uncropped blots are shown in Extended Data Fig. 14. **h,** Detection of *IDO1* mRNA levels in SKOV-3 cells upon treatment with succinyl acetone (SA) for 24 h followed by addition of hemin for another 24 h. Mean values ± SD (n = 3). Uncropped blots are shown in Extended Data Fig. 13-14. UB: ubiquitin.

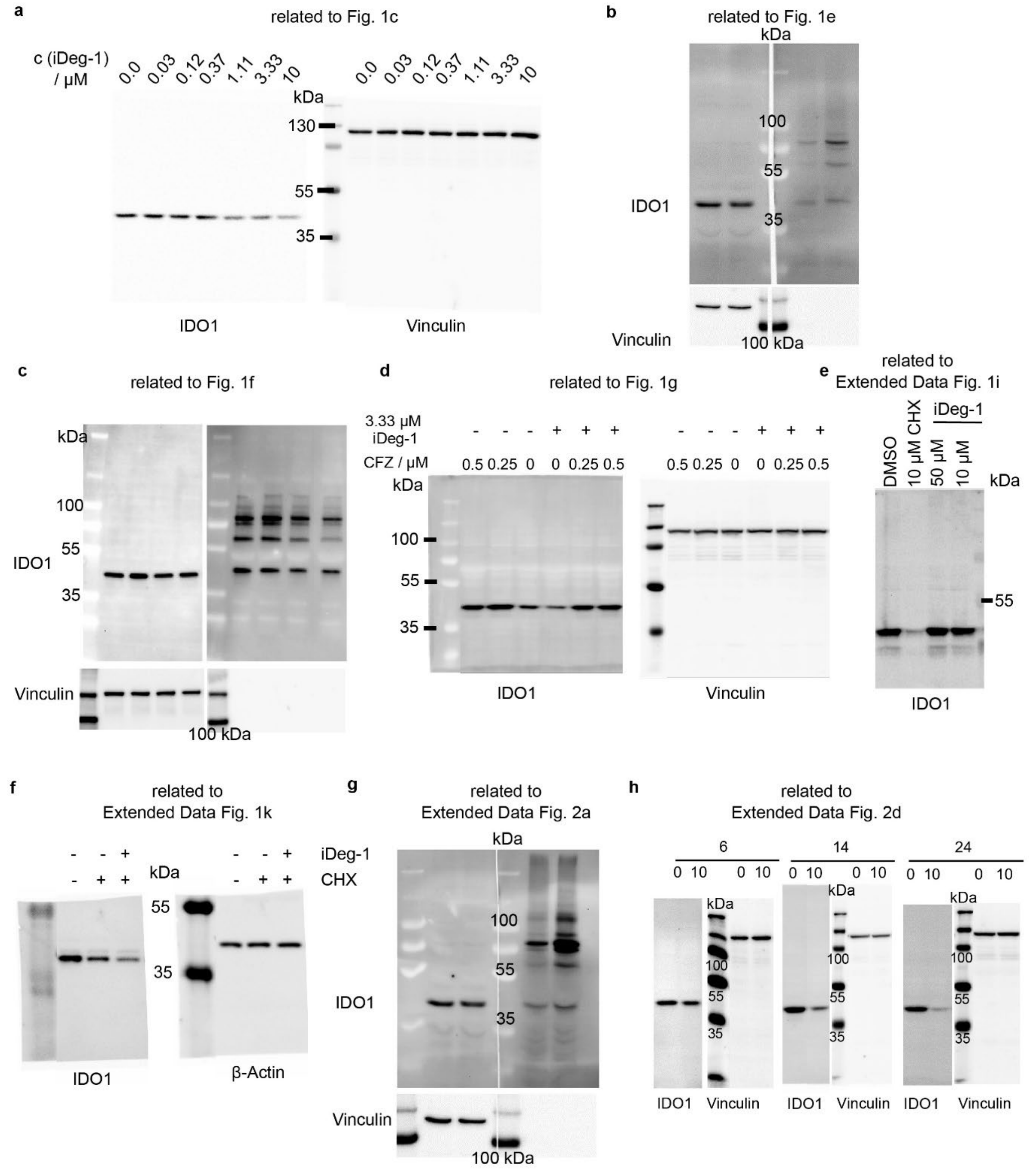

**Extended Data Fig. 11: Uncropped immunoblots.** Related to Figure 1 and Extended Fig. 1-2.

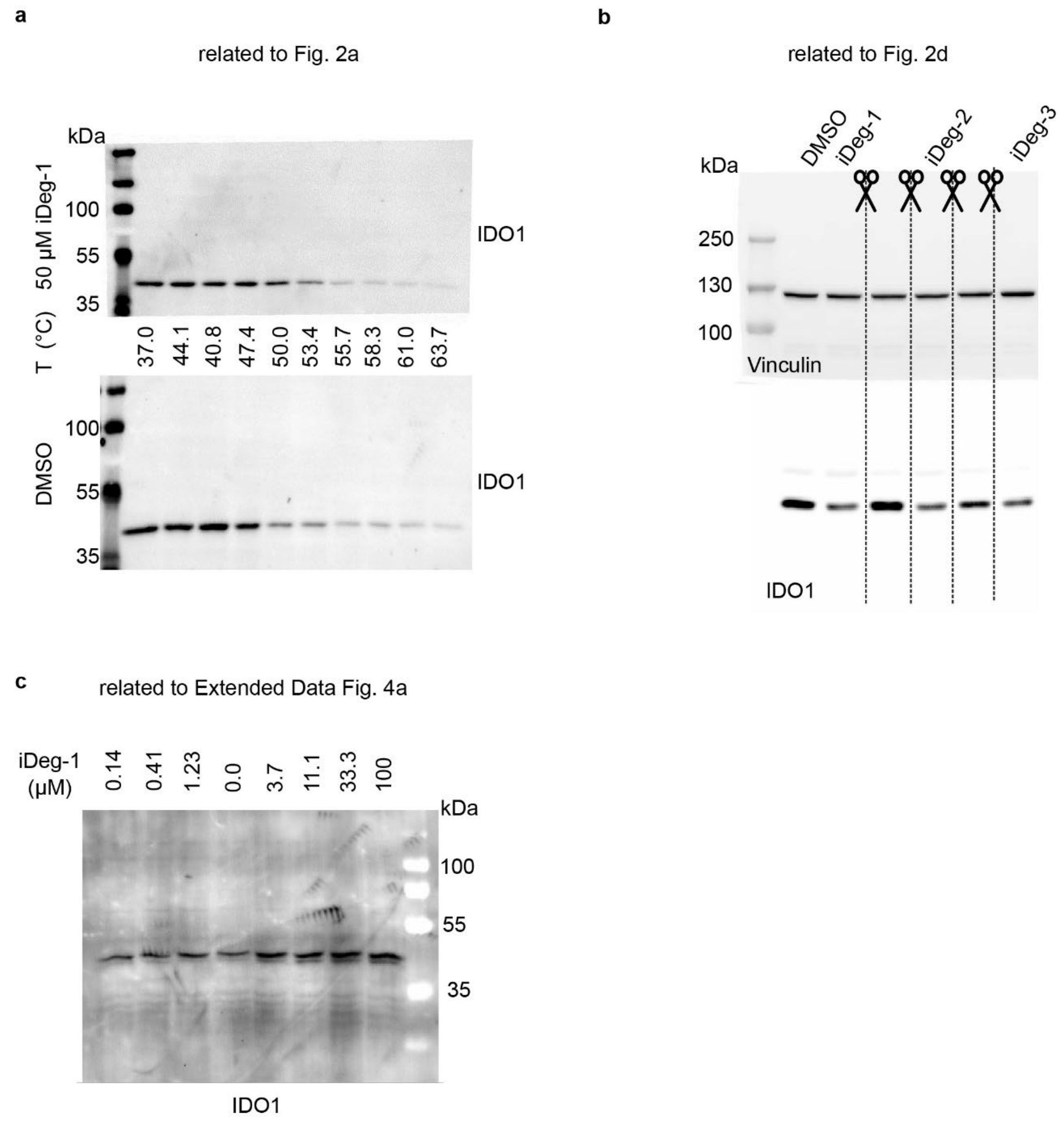

**Extended Data Fig. 12: Uncropped immunoblots.** Related to Fig. 2 and Extended Data Fig. 4.

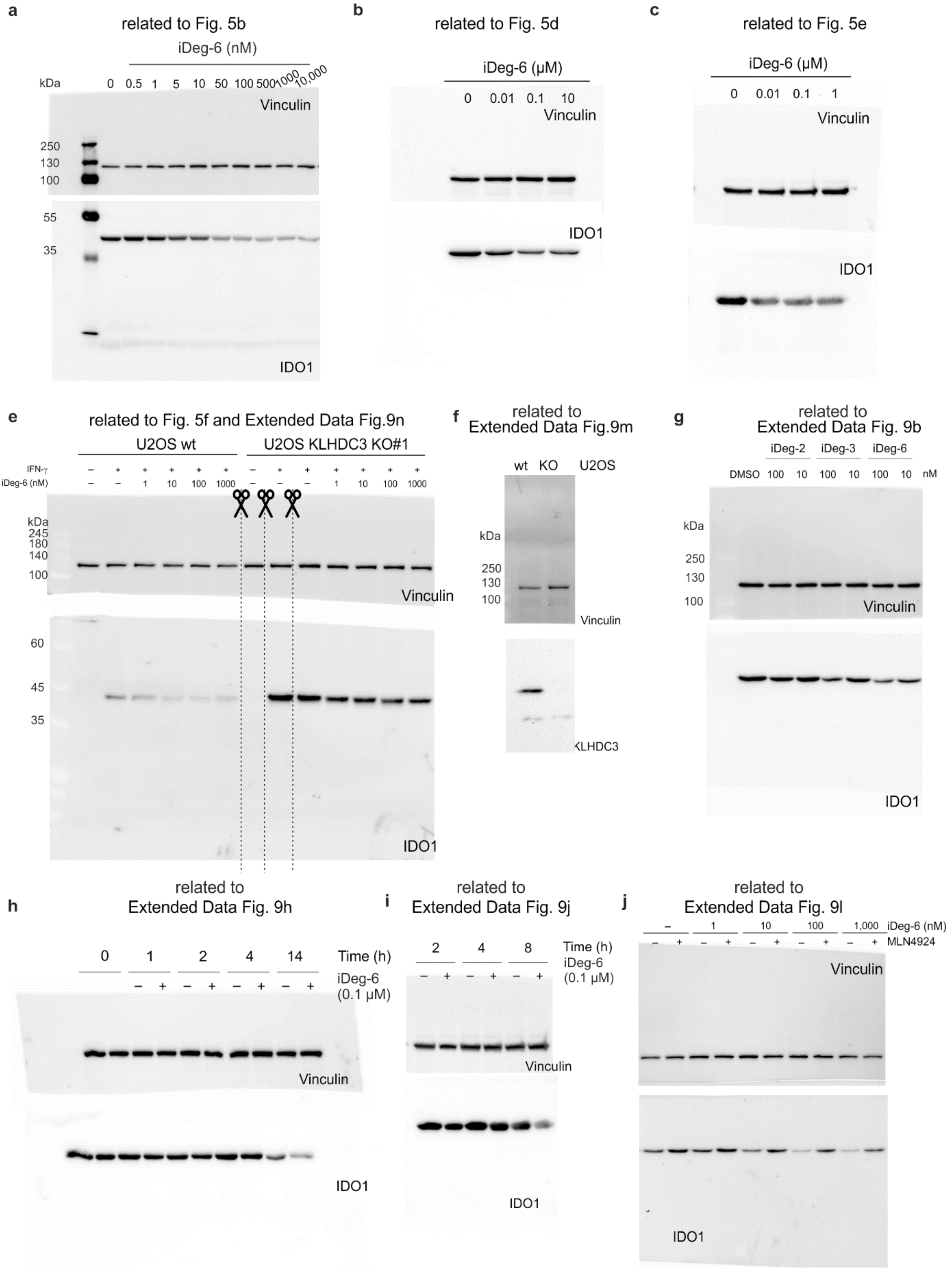

**Extended Data Fig. 13: Uncropped immunoblots.** Related to Fig. 5 and Extended Data Fig. 9.

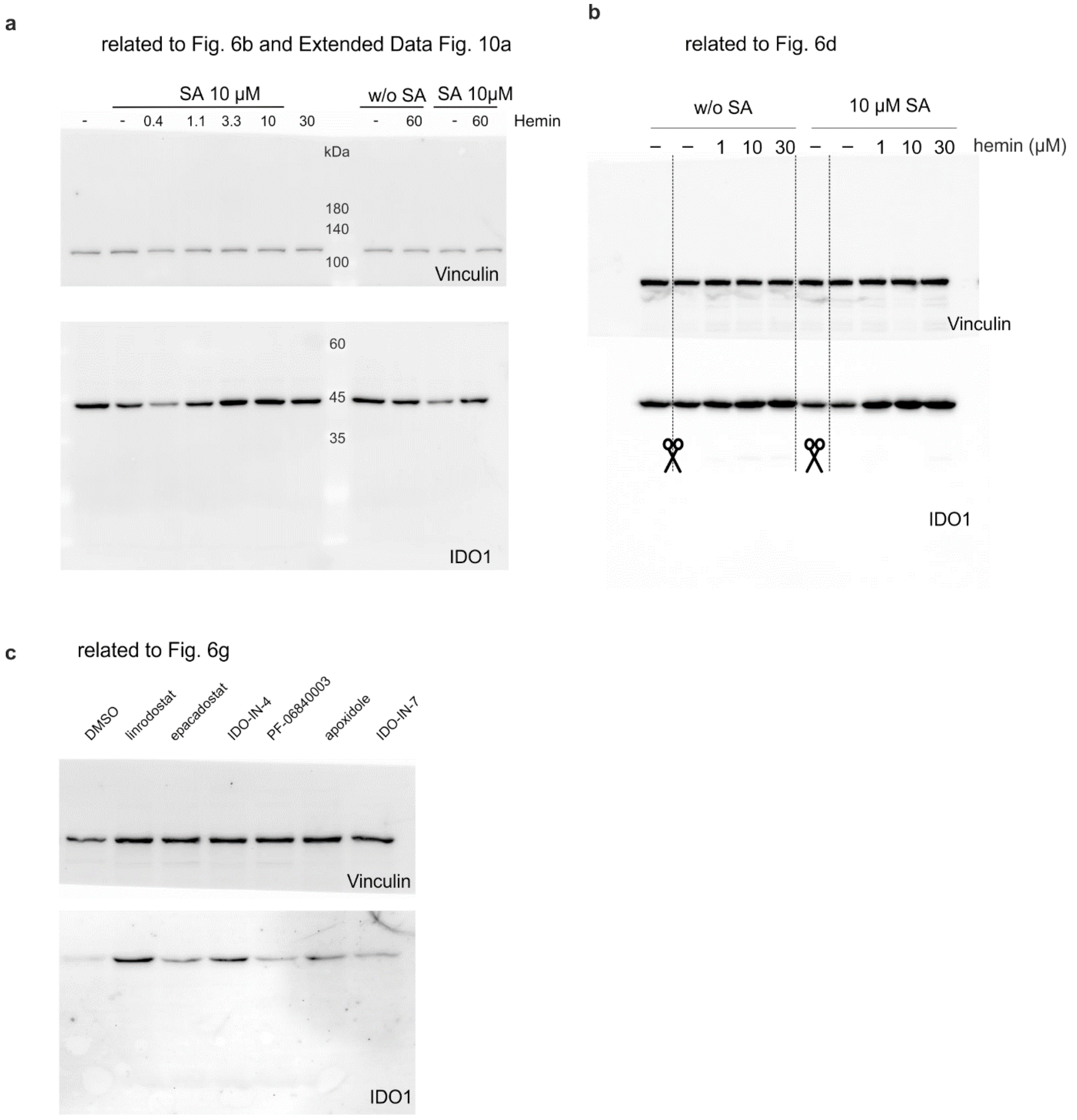

**Extended Data Fig. 14. Uncropped immunoblots.** Related to Fig. 6 and Extended Data Fig. 10.

### Extended Data Tables

**Extended Data Table 1. Changes in protein levels after treatment with iDeg-1.** HEK239T cells were electroporated with rhIDO1 protein prior to treatment with 10 µM iDeg-1 or DMSO for 6 h and MS analysis.

| **-log(P-value)** | **Difference** | **Protein names** | **Gene names** |
|---|---|---|---|
| 6.08 | -0.66 | Dedicator of cytokinesis protein 8 | DOCK8 |
| **5.10** | **-0.85** | **Indoleamine 2,3-dioxygenase 1** | **IDO1** |
| 3.13 | 0.78 | Ras-related protein Rab-9B | RAB9B |
| 1.29 | -1.13 | Rho-related BTB domain-containing protein 3 | RHOBTB3 |
| 1.13 | 1.24 | Zinc finger protein 64 homolog, isoforms 3 and 4 | ZFP64 |

**Extended Data Table 2: Data collection and refinement statistics of apo-IDO1 with iDeg-2 structure.** Values in parentheses correspond to the highest-resolution shell.

| | **apo-IDO1 + iDeg-2** |
|---|---|
| **PDB code** | **9FOH** |
| **Data collection** | |
| Space group | P $2_1$ $2_1$ 2 |
| Cell dimensions | |
| *a*, *b*, *c* (Å) | 104.84 110.42 36.14 |
| α, β, γ (°) | 90 90 90 |
| Resolution range (Å) | 34.73-1.60 (1.66-1.60) |
| Total reflections | 715765 (62798) |
| Unique reflections | 56429 (5574) |
| $R_{meas}$ | 0139 (2.43) |
| $R_{\mathrm{pim}}$ | 0.0388 (0.718) |
| CC1/2 | 1.00 (0.69) |
| $I / \sigma I$ | 13.57 (1.42) |
| Completeness (%) | 99.96 (99.95) |
| Redundancy | 12.7 (11.3) |
| **Refinement** | |
| Resolution (Å) | 1.6 |
| No. reflections | 56419 (5572) |
| $R_{\mathrm{work}}$ / $R_{\mathrm{free}}$ | 0.171 / 0.189 |
| No. of non -H atoms | 3272 |
| Protein | 2927 |
| Ligand/ion | 231 |
| Water | 243 |
| *B*-factors | 29.13 |
| Protein | 27.87 |
| Ligand | 44.16 |
| Water | 38.04 |
| R.m.s. deviations | |
| Bond lengths (Å) | 0.12 |
| Bond angles (°) | 1.67 |

**Extended Data Table 3: Oligo sequences for stability reporter generation and single sgRNA cloning.**

| Oligo | Sequence (5' → 3') | Reporter |
|---|---|---|
| mCherry_fw | ctcggcgcgccagtcctccgagtcgacccacc atggtgagcaagggcgaggag | N-terminal stability reporter design |
| mCherry_rev | cacatctccggcctgcttcagcaggctgaagtt ggtggcgccgctgcctttatacaattcatccatg ccgc | N-terminal stability reporter design |
| BFP_fw | ctgctgaagcaggccggagatgtggaggaga accctggccccagcgagctgattaaggagaa c | N-terminal stability reporter design |
| BFP_rev | cagaggttgattgttccagacgcgtattaagctt gtgccccagtttg | N-terminal stability reporter design |
| IDO1_cterm_fw | cggcgcgccagtcctccgagtcgacgccacc atggcccacgccatgg | C-terminal IDO1 stability reporter |
| IDO1_cterm_rev | cacctcccgagccacctccggatcctccaccg ccctccttcagcaggctc | C-terminal IDO1 stability reporter |
| IDO1_nterm_fw | tactgcgacctccctagcaaactggggcacaa gcttaatacgcgtatggcccacgccatgg | N-terminal IDO1 stability reporter |
| IDO1_nterm_WT_rev | ctttcacaaattttgtaatccagaggttgattgttc cagatcagccctccttcagcaggctcttctcggt ggtg | N-terminal IDO1 stability reporter |
| IDO1_nterm_RG_rev | ctttcacaaattttgtaatccagaggttgattgttc cagatcagccTCTcttcagcaggctcttctcg gtggtg | Degron mutant "-RG" N-terminal IDO1 stability reporter |
| IDO1_nterm_EK_rev | ctttcacaaattttgtaatccagaggttgattgttc cagatcaCTTctccttcagcaggctcttctcgg tggtg | Degron mutant "-EK" N-terminal IDO1 stability reporter |
| sgRNA_KLHDC3_1_fw | caccGCATCGGGTATACTCCTTC G | Exon 2 targeting sgRNA for KLHDC3 (corresponds to KO1) |
| sgRNA_KLHDC3_1_rev | aaacCGAAGGAGTATACCCGATG C | Exon 2 targeting sgRNA for KLHDC3 (corresponds to KO1) |
| sgRNA_KLHDC3_2_fw | caccGGTGTCAAAGACTCGAATG | Exon 6 targeting sgRNA for KLHDC3 (corresponds to KO2) |
| sgRNA_KLHDC3_2_rev | aaacCATTCGAGTCTTTGACACC | Exon 6 targeting sgRNA for KLHDC3 (corresponds to KO2) |
| sgRNA_AAVS1_fw | caccGCTGTGCCCCGATGCACAC | CTRL sgRNA |
| sgRNA_AAVS1_rev | aaacGTGTGCATCGGGGCACAGC | CTRL sgRNA |

#### Material and Methods

##### Data availability

Data that support the findings of this study have been deposited in MassIVE with the accession code MSV000094270, PXD050474 (global proteome profiling) and MSV000094271, PXD050475 (IDO1 immunoprecipitation).

The crystal structure of IDO1 with iDeg-2 was deposited in PBD with the accession number 9FOH.

##### Material

BxPC3 cells, HEK293T and HeLa cells were purchased from DSMZ GmbH (Germany), BT549, SKOV-3 and U2OS cells obtained from ATCC (USA). U2OS KO1 cells were obtained from St. Jude Center for Advanced Genome Engineering (CAGE) (Memphis, USA). KBM7 iCas9 cells were a gift from J. Zuber (IMP, Vienna). Lenti-X 293T lentiviral packaging cells were obtained from Clontech. pCMV3-IDO1 was obtained from Sino Biological US Inc. pSpCas9(BB)-2A-GFP (PX458) plasmid was a gift from Feng Zhang (Addgene plasmid # 48138; http://n2t.net/addgene:48138; RRID:Addgene_48138[1])). pCMVR8.74 was a gift from Didier Trono (Addgene plasmid # 22036; http://n2t.net/addgene:22036 ; RRID:Addgene_22036). pMD2.G was a gift from Didier Trono (Addgene plasmid # 12259; http://n2t.net/addgene:12259; RRID:Addgene_12259). Fetal bovine serum (FBS), penicillin/streptomycin, anti-IDO1 (14-9750-80) and anti-vinculin (V9131) antibodies were obtained from ThermoFisher, USA. Anti-KLHDC3 (HPA030131) was obtained from Sigma-Aldrich. Anti-Rabbit HRP (7074) was obtained from Cell Signaling Technology. anti-IDO1 (ab211017) and anti-β-actin (ab8227) antibodies were obtained from abcam. Mouse Thy1.1 antibody was purchased from BioLegend. IRDye-conjugated secondary antibodies were obtained from LI-COR Biosciences. DMEM, McCoy's medium, sodium pyruvate, and non-essential amino acids were obtained from PAN biotech, Germany. Where indicated, DMEM was obtained from Gibco, USA. IMDM was obtained from Gibco, USA. 1 mM L-tryptophan was obtained from Sigma. IFN-γ was purchased from 50 ng/mL, PeproTech.Succinyl acetone was obtained from TCI. Doxycycline was obtained from PanReac AppliChem. IDO-IN-1 (19402) and IDO-IN-5 (33178) were obtained from Cayman Chemical. IDO-IN-4 (HY-18769), IDO-IN-7 (HY-13983), PF-06840003 (HY-101111), linrodostat (BMS 986205, HY-101560) and epacadostat (HY- 15689) were obtained from MedChem Express.

*Cell Culture*

BxPC3 cells were cultured in RPMI-1640 (PAN biotech, Germany) containing 10% heat-inactivated fetal bovine serum (FBS). BT549, HeLa and HEK293T cells were cultured in DMEM (PAN biotech, Germany) supplemented with 10 % FBS, sodium pyruvate and non-essential amino acids. SKOV-3 and U2OS wt and knockout cells were cultured in McCoy's 5A (PAN biotech, Germany) medium containing 10% heat-inactivated FBS. Lenti-X 293T lentiviral packaging cells were cultured in DMEM (Gibco) supplemented with 100 U/mL penicillin/streptomycin and 10% heat-inactivated FBS. IDO1-deficient SKOV-3 cells were produced using the CRISPR-Cas9 system. For this, 100,000 cell/well in 6-well plates were transfected using Lipofectamine 3000 with the PX458 plasmid containing three separated sgRNA located in *IDO1*. sgRNA sequences: 5'-CACACGCTATGGAAAACTCC-3'; 5'-AGCCCACTTCTTCATCAATA-3'; 5'-TACCATATTGATGAAGAAGT-3'. 48 hours later, the cells were sorted for single cells based on GFP positivity. Single clones were expanded and successful knockout of *IDO1* was verified using immunoblotting. KBM7 iCas9 cells were cultured in IMDM supplemented with 100 U/mL penicillin/streptomycin, 10% heat-inactivated FBS and 1 mM L-tryptophan. Cell lines were maintained in a humidified atmosphere at 37°C and 5 % $CO_2$. Regular mycoplasma testing confirmed cell lines as mycoplasma-negative.

*Automated Kyn Screening Assay*

The Kyn screening assay was performed as previously described[2]. BxPC3 cells were seeded in phenol red-free RPMI 1640 medium in 1536-well or 384-well plates (Greiner #782086, Corning #3770) 24 h prior to the addition of compounds, IFN-γ (50 ng/mL, PeproTech) and L-Trp (380 µM, Sigmal-Aldrich). After 48 h of incubation, first cell viability was accessed by the addition of Hoechst-33342. Afterwards, trichloroacetic acid (TCA, Sigma-Aldrich) was added to a final concentration of 7 % and incubated for 10 min prior to a centrifugation step of 10 min at 1620 x g. Kyn levels were determined by means of the Kyn sensor[2] at a final concentration of 17.5 µM in sensor buffer (50 mM $H_3PO_4$ and 120 mM NaCl (pH=1)). Fluorescence intensity (excitation: 535 nm, emission: 595 nm) was measured using a Spectramax Paradigm reader (Molecular Devices). Data were normalized to the values for cells that were treated with DMSO. All HTS data was analyzed using the Quattro Software Suite (Quattro Research GmbH).

*Manual Kyn Assay*

For the manual assay, BxPC3 or SKOV-3 cells were seeded in clear 96-well or 384-well plates, respectively, 24 h prior to compound treatment and IFN-γ (50 or 5 ng/mL, respectively) and L-Trp (380 µM) addition as described before.[2] 48 h after treatment, TCA was added and Kyn levels were detected by means of 2 % w/v of p-DMAB (Ehrlich reagent) in acetic acid as described by Braun et al[29]. The absorbance of the Ehrlich reagent was detected at 492 nm and at 650 nm as background control on Spark® multimode microplate plate reader (Tecan, Austria). Kyn levels were determined by subtracting the absorbance value at 650 nm and presented relative to the DMSO control. Dose-response curves and $IC_{50}$ values were generated and fitted with GraphPad Prism 9.2 (GraphPad software, USA) using four-parameter variable slope non-linear regression curve fit.

For label-free Kyn detection, the cell supernatant was analyzed via LC-MS/MS upon the addition of 30 % TCA. Kyn levels were measured by HPLC-MS/MS using the LTQ Velos Pro and Dionex HPLC (Thermo Fisher Scientific). Data were analyzed using Thermo Xcalibur™ (Thermo Fisher Scientific) and presented with GraphPad Prism 9.2 (GraphPad software, USA). Values are shown relative to the DMSO control.

*Live-cell Analysis*

SKOV-3 WT and IDO1-deficient cells were seeded in 48-well-plates (8000 cells/well, Corning, 3548) for 2D experiments or in a 96 ultra-low attachment plates (5000 cells/well, S-bio, MS-9096UZ) for 3D experiments. After seeding, cells were treated with 10 ng/mL (2D) or 1 ng/mL (3D) of IFN-γ and CellTox Green (Promega, G8731) was added in the culture media (1:6000). The following day, cells were treated with either DMSO or 5 µM of iDeg-1. iDeg-1 was freshly added every three days. Cells were monitored by live phase-contrast microscopy using the IncuCyte S3 system (Sartorius) with the 10X objective, capturing images every day up to 4 days (2D) and 6 days (3D). Cell death was quantified by counting the numbers of green fluorescence positive cells (CellTox Green positive) normalized to the confluence of each respective image (2D) or by measuring the green signal mean intensity of each spheroid (3D).

*Immunoblotting*

To analyze IDO1 levels, IFN-γ-stimulated BxPC3 cells (50 ng/mL IFN-γ), U2OS cells (5 ng/mL IFN-γ), $HEK^{rhIDO1}$ (HEK293T cells electroporated with rhIDO1), SKOV-3 and BT549 cells were treated with compounds for the indicated times (1 h to 24 h) followed by cell lysis (150 mM sodium chloride in 50 mM Tris (pH 8.0), 1% NP-40 Alternative, protease and phosphatase inhibitors). Protein concentrations were determined (DC protein assay, Bio-Rad) followed by addition of 1 x SDS loading buffer and protein separation by a 10 % SDS-PAGE. Separated protein samples were transferred to a polyvinylidene difluoride (PVDF) membrane using a wet-tank blotting system (Bio-Rad). IDO1 in IFN-γ-stimulated cells as well as vinculin or β-actin as loading controls were detected by blocking the membranes with Intercept® Blocking Buffer (LI-COR Biosciences, 927-70001) and using the primary antibodies anti-IDO1 (1:2500, 4-9750-82, Thermo Fisher), anti-vinculin (1:10,000, V9131, Thermo Fisher) and anti-β-actin (ab8227, abcam) and IRDye-conjugated secondary antibodies (1:5000, LI-COR Biosciences). For detection of IDO1 protein levels in BT549 and SKOV-3 cells, membranes were blocked with 5% milk in PBS + 0.1% Tween-20 (PBS-T), followed by incubation with anti-IDO1 (1:5000, ab211017, abcam) and HRP conjugated secondary antibody (1:5,000, 31460, Pierce Biotechnology). KLHDC3 protein levels in U2OS and U2OS KO cells were detected by blocking the membranes with 5% milk in PBS-T and using anti-KLHDC3 antibody (1:1000, HPA030131, Atlas) and anti-rabbit HRP-conjugated secondary antibody (1:3000, 7074, Cell Signaling Technology). Primary antibodies were incubated at 4°C overnight. Washing steps were performed with PBS-T. Chemiluminescence was detected using SuperSignal™ West Dura Extended Duration Substrat (Thermo Fisher Scientific). Protein bands were visualized using the ChemiDoc™ MP system (Bio-Rad). Band intensities were quantified using Image Lab software 6.0 (Bio-Rad).

*Reverse Transcription-Quantitative PCR*

BxPC3 cells (100,000 cells/well) or SKOV-3 cells (150,000 cell/well) were seeded in a 24-well plate or 12-well plate, respectively. BxPC3 cells were incubated for 24 h prior to treatment with the indicated compound concentrations and 50 ng/mL IFN-γ. SKOV-3 cells were directly treated with or without 10 µM succinyl acetone for 24 h prior to the addition of hemin for further 24 h. Total RNA was isolated by means of RNeasy Mini Kit as described by the manufacturer (Qiagen). Obtained RNA was reverse transcribed into cDNA by means of QuantiTect Reverse transcription kit according to the manufacturer’s instructions (Qiagen) prior to the qPCR using QuantiFast SYBR green PCR kit (Bio-Rad). Gene-specific primers for *IDO1* (fw: 5’-GCCTGATCTCATAGAGTCTGGC-3’ and rv: 5’-TGCATCCCAGAACTAGACGTGC-3’)[30] and GAPDH (fw: 5’- GTCTCCTCTGACTTCAACAGCG-3’ and rv: 5’-ACCACCCTGTTGCTGTAGCCAA-3’) as a reference gene were used. qPCR was performed using CFX96 Real-Time PCR Detection System (Bio-Rad). Data were analyzed using the CFX Manager (Bio-Rad). *IDO1* expression levels in each sample were normalized to the levels of the reference gene *GAPDH*. Relative quantification was performed using the $2^{-\Delta\Delta Ct}$ method[3].

*IDO1 Promoter-Dependent Reporter Gene Assay*

For the assay, the pXPG-IDO1 plasmid348 containing a firefly luciferase (Fluc) gene under the control of the full-length *IDO1* promoter (kindly provided by Gina M. Doody, Leeds, UK) was used as a reporter. The *Renilla* (Rluc) luciferase expression plasmid pRL-TK vector (Promega Corporation, US) was used as a control. HEK293T cells were transiently reverse-transfected with the plasmids using Lipofectamine™ 2000 (Invitrogen, US) following a modified protocol of the manufacturer's guidelines. 4 µg of the Fluc plasmid and 300 ng of the Rluc plasmid were mixed in serum-free Opti-MEM in a 1:3 ratio with Lipofectamine™ 2000. The mixture was added to 9 mL cell suspension at a cell density of 2.78 x $10^5$ cells/mL. 25,000 cells were seeded per well in a clear 96-well plate and incubated for 24 h. On the next day, *IDO1* promoter-mediated Fluc expression was induced by addition of 50 ng/mL IFN-γ with simultaneous compound treatment. Cells were incubated for 48 h prior to determining the activity of Fluc and Rlus using the Dual-Glo® Luciferase Assay System with the Spark® Multimode Microplate Reader (Tecan, AT). Fluc / Rluc ratios were normalized to the values of the DMSO control (set to 100 %).

*CHX Pulse Assay*

BxPC3 cells (200,000 cells/well) were seeded in a 12-well plate and 50 ng/mL IFN-γ were added. After 24 h, 5 µM CHX was added 30 min prior to the addition of 5 µM iDeg-1 followed by cell lysis preparation and immunoblotting.

*TUBE Pulldown*

Tandem ubiquitin binding entity protein (TUBE) was expressed from pGex-6P-1-GST-4xUBA-His (UBA: Ubiquitin-associated domains) in *E. coli* and purified by means of affinity purification using a Ni-NTA column. TUBE protein expression was induced by IPTG for 4 h at 30 °C. Afterwards cells were lysed in buffer containing 20 mM sodium phosphate (pH 7.4), 500 mM NaCl, 20 mM imidazole, 1 mM PMSF (Phenylmethylsulfonylfluorid, 39107.01, Serva) and 1 mM 2-mercaptoethanol and TUBE protein was separated using a Ni-NTA affinity chromatography. The TUBE protein was eluted by 20 mM sodium phosphate (pH 7.4) buffer containing 500 mM imidazole and 1 mM 2-mercaptoethanol and subsequently subjected to a size exclusion column (HiLoad 16/600 Superdex 200 pg) in 20 mM potassium phosphate buffer (pH 7.4, supplemented with 300 mM NaCl and 1mM DTE and 2 % Glycerol) to yield pure TUBE3.

For the TUBE pulldown, BxPC3 cells were seeded in either 10 or 15 $cm^2$ cell culture dishes and after cell attachment treated with 40 ng/mL IFN-γ to induce IDO1 expression overnight.

Next, the medium was aspired and medium containing the respective concentration of iDeg-1 was added and incubated for 6 h at 37 °C and 5 % $CO_2$. In case of proteasome inhibition, 450 nM Carfilzomib (CFZ, ab216469, abcam) was added 1 h prior to addition of iDeg-1. After compound incubation, cell lysates were prepared in TUBE lysis buffer (20 mM $Na_2HPO_4$, 20 mM $NaH_2PO_4$, 1 % NP-40 Alternative, 2 mM EDTA and 10 mM N-ethylmaleimide (NEM, freshly prepared)) supplemented with 100 µg/mL TUBE protein. Samples were kept on ice for 30 min followed by a centrifugation step for 30 min at 16,000 x g at 4°C. Protein concentrations were determined to adjust protein concentration prior to the addition of 40 µL GST magnetic beads (ThermoFisher). Samples were rotated for 2 h at 4 °C followed by four washing steps using ice-cold PBS-T for 5 min. To elute ubiquitinated protein species, 20 µL 1x SDS sample buffer were added to the beads and incubated for 15 min at 55 °C. Afterwards, all samples were boiled for 5 min at 98 °C prior the separation by an SDS-PAGE and immunoblotting for IDO1 and vinculin as a control.

*Electroporation of rhIDO1 into HEK293T Cells*

HEK239T (3,000,000 cell/per electroporation) cells were electroporated with 80 µg rhIDO1 utilizing the NeonTM transfection kit according to the manufactures description (ThermoFisher). For electroporation, 1000 V and 2 pulses for 35 ms were applied using the Neon™ transfection system pipette station. Afterwards, cells were washed in pre-warmed PBS followed by resuspension in 3 mL trypsin/EDTA solution and incubated for 3 min at 37 °C. Electroporated cells were washed again and resuspended in phenol red-free DMEM growth medium (900,000 cells/mL). For the Kyn assay in $HEK^{rhIDO1}$ cells, cells were plated in 96-well plates and supplemented with 150 µM L-Trp and the respective compound concentrations for 24 h prior read out as described above for the manual Kyn assay. For immunoblotting, electroporated cells were seeded in a 12-well plate and treated with the indicated compounds for 6, 14 or 24 h prior to immunoblotting as described above. For co-treatment, cells were pre-treated with the proteasome inhibitor CFZ or the neddylation inhibitor MLN4924 for 30 min prior to iDeg-1 addition.

*HEK293T-based Kyn Assay*

HEK293T (270,000 cell/mL) cells were transfected with 0.5 µg pCMV3-IDO1 (Sino Biological US Inc.) using Lipofectamine 2000 (Thermo Fisher) while seeding in a 96-well plate followed by incubation overnight. Next, 500 µM L-Trp and compounds at the indicated concentrations were added and cells were incubated for 24 h prior to the detection of Kyn levels using p-DMAB as described above for the manual Kyn assay.

*Proteome Profiling*

HEK239T cells were electroporated with rhIDO1 as described above and treated with 10 µM iDeg-1 or DMSO for 6 h prior to lysate preparation by means of four freeze-thaw cycles in PBS containing 0.4 % NP-40 Alternative. Protein concentration was determined and 200 µg of protein lysates were subjected to the sample preparation and MS analysis. Protein lysates were dissolved in equal volume of 100 mM triethylammonium bicarbonate (TEAB) buffer supplemented with 7.5 µL of a 200 mM Tris(2-carboxyethyl)phosphine (TCEP) and incubated at 55°C for 1 h. Afterwards, 7.5 µL of 375 mM iodoacetamide were added and samples were incubated for another 30 min at room temperature in the dark prior to acetone-based protein precipitation (incubation at -20 °C overnight). On the next day, samples were centrifuged at 8,000 x g and 4 °C, the supernatant was aspired and protein pellets were dried at room temperature. Pellets were dissolved in 50 mM TEAB containing Trypsin/Lys-C (25:1 protein:protease ratio (w/w), pre-dissolved in 1 mM HCl (Trypsin/Lys-C Mix, Mass spec. grade, 5x from Promega Corporation)) and incubation at 37 °C overnight, while shaking. On the next day, samples were dried in a vacuum concentrator at 30 °C and subjected to MS.

*Mass Spectrometry (MS)*

All solvents used during MS measurements were LC-MS grade. Digested samples were first dissolved in 120 µL of 20 mM ammonium formate ($NH_4COOH$) at pH 11 in an ultra-sonicator for 15 min followed a 15 min incubation step, while rotating and a subsequent centrifugation step (8,000 x g for 3 min at room temperature). 50 µL of each sample was separated into 10 fractions by means of a XBridge C18 column (130 Å, 3.5 µm, 1mm x 150 mm) using a U3000 capHPLC system (Thermo Fisher Scientific, Germany) under high pH conditions to reduce sample complexity an increase the number of quantified proteins. Separation was performed at a flow rate of 50 µl/min starting with 95 % solvent A (20 mM $NH_4COO$ pH 11 in $H_2O$) / 5% solvent B (40% 20 mM $NH_4COO$ pH 11, 60 % acetonitrile in $H_2O$) isocratic for the first 10 min, followed by a linear gradient increasing solvent B up to 25 % over 5 min. Afterwards, a second linear gradient was applied, increasing solvent B up to 65 % in 60 min, followed by a third gradient increasing solvent B up to 100% over 10 min. Fractions were detected at a wavelength of 214 nm.

The 10 fractions were collected 15 and 100 min (30 s per fraction, circular fractionation using 10 vials) and dried in a SpeedVac at 30°C until complete dryness for the final analysis by nanoHPLC-MS/MS. The dried samples were then dissolved in 10 µL of 0.1% TFA in water prior to the injection of 9 µL onto an UltiMateTM 3000 RSLCnano system (Thermo Fisher Scientific, Germany) online coupled to a Q Exactive™ HF Hybrid Quadrupole-Orbitrap Mass

Spectrometer equipped with a nanospray source (Nanospray Flex Ion Source, Thermo Fisher Scientific).

To finally detect the peptides within the samples a mass range of m/z 300 to 1650 was attained with a resolution of 60000 for full scan prior to 15 high energy collision dissociation (HCD) MS/MS scans of the most intense at least doubly charged ions using a resolution of 15 000 and an NCE energy of 27%. The obtained data were analyzed using MaxQuant software (v.1.6.17.0)[4] including the Andromeda search algorithm and using the human reference proteome of the Uniprot database. A full enzymatic trypsin cleavage search was conducted allowing for two miscleavages. Carbamidomethylation was chosen as fixed modification and oxidation of methionine and acetylation of the N-terminus as variable modifications.

For the first analysis, the mass accuracy for full mass spectra was set to 20 ppm while in the second search it was set to 4.5 ppm and for the MS/MS spectra to 20 ppm. A false discovery rate of 1 % was selected for peptide and protein identification and only proteins for which at least two peptides were quantified were chosen for further validation. Relative protein quantification was achieved by means of the label-free quantification algorithm implemented in MaxQuant.

To consider the differences of IDO1 concentration after electroporation of IDO1 into the HEK293T cells, all LFQ-intensities were normalized to those of the DMSO control of the same biological replicate by dividing the LFQ-intensity of the sample through that of the corresponding DMSO control (resulting in a value of 1 for the DMSO-controls themselves).

Statistical data analysis was completed using Perseus[5] v.1.6.14.0. The normalized label-free quantification (LFQ) intensities were log-transformed (log2) and grouped. Only proteins were retained for further analysis when quantified at least two times in at least one of the groups. Missing values were ascribed to '0' and a t-test (FDR=0.01, s0=0.5) was conducted. Results were exported into an Excel file (xlsx) and visualized using Perseus. Detailed analysis procedure is described by Cox *et al.* and Tyanova *et al.*[4,5]

*IDO1 Immunoprecipitation Followed by MS Analysis*

BxPC3 cells were seeded in 10 cm$^2$ dishes and stimulated with 40 ng/mL IFN-γ overnight at 37 °C and 5 % $CO_2$. On the next day, fresh medium was added containing 20 µM of iDeg-1 or DMSO a as control and incubated for 8 h prior cell detachment and cell lysis using immunoprecipitation lysis buffer (20 mM Tris-HCl pH 7.5, 100 mM NaCl, 0.1 mM EDTA and 0.5 % NP-40 Alternative) supplemented with fresh N-ethylmaleimide as well as phosphatase (PhosSTOP phosphatase inhibitors, Sigma-Aldrich) and protease inhibitors (cOmplete™ protease inhibitor cocktail, Sigma-Aldrich). Three freeze-and-thaw cycles were performed for cell lysis followed by centrifugation at 14,000 x g for 20 min at 4 °C and determination of the protein concentration (DC protein assay, Bio-Rad). 800 µg lysate was subjected to the immunoprecipitation.

For the following steps, a magnetic rack was used to separate magnetic beads from its supernatant. First, cell lysates were pre-cleared using 25 µL of Pierce Protein A/G Magnetic beads per sample for 1 h at 4°C while rotating. Afterwards, pre-cleared lysates were added to pre-coated Pierce Protein A/G Magnetic beads with IDO1 antibody (ab211017, abcam) followed by continuous rotation for 2 h at 4 °C. Beads were washed three times with immunoprecipitation wash buffer (50 mM PIPES pH 7.4, 150 mM NaCl, 5 mM $MgCl_2$, 0.5 mM EGTA, 0.1 % NP-40 alternative, 0.1 % Triton X-100, 0.1 % Tween® 20 and 1 mM DTT) for 5 min while rotating. Subsequently, beads were transferred to a fresh sample tube after being washed using three times 500 µL PBS prior to further washing steps with 500 µL PBS (thrice) to remove detergents. Afterwards, beads were resuspended in 50 µL denaturation buffer (50 mM Tris pH 7.5, 8 M Urea and 1 mM DTT) and incubated for 30 min while shaking prior to addition of 5.55 µL 50 mM chloroacetamide solution for another 30 min while shaking. For on bead digestion, 1 µL LysC (stock solution of 0.5 µg/µL in water) was added for 1 h at 37 °C while shaking. The LysC digestion supernatant was transferred into a fresh sample tube and to the remaining beads 165 µL of a 50 mM Tris (pH 7.5) solution containing 0.25 µg trypsin was added and incubated for another 1 h at 37 °C while shaking. The two digestion solutions were combined and supplemented with additional 0.5 µg trypsin and the digestion was continued overnight at 37 °C while shaking. The remaining beads were discarded. On the next day, the addition of 20 µL 10 % TFA stopped the digestion and samples were desalted by means of stage tip purification. A double layer of C18 chromatography matrix was placed into 200 µL pipette tip and the matrix was activated by adding 100 µL of 100 % methanol. The matrix was washed once with buffer A (0.1 % formic acid, 80 % acetonitrile in $H_2O$) and twice with buffer B (0.1 % formic acid in $H_2O$). For each washing step, a centrifugation in a tip centrifuge was performed. Afterwards, samples were subjected to the tip washed once with buffer B prior sample elution using twice 20 µL buffer A. The samples were dried in a vacuum concentrator at 30 °C and analyzed by MS.

All solvents used during MS measurements were LC-MS grade. The samples were dissolved in 20 µl of 0.1% TFA in water and 3 µl were subjected to nanoHPLC-MS/MS analysis. Therefore, the samples were injected onto UltiMateTM 3000 RSLCnano system (ThermoFisher scientific, Germany) online coupled to a Q Exactive™ Plus Hybrid Quadrupole-Orbitrap Mass Spectrometer equipped with a nanospray source (Nanospray Flex Ion Source, Thermo Scientific). Samples were desalted by means of a pre-column cartridge (5 µm, 100 Å, 300 µm ID * 5 mm, Dionex, Germany) using 0.1% TFA in water and a flow rate of 30 µL/min for 5 min with eluent flow to waste followed by back-flushing of the sample during the whole analysis from the pre-column to the PepMap100 RSLC C18 nano-HPLC column (2 µm, 100

Å, 75 µm ID × 50 cm, nanoViper, Dionex, Germany). Therefore, a linear gradient starting with 95% solvent A (0.1% formic acid in $H_2O$) and 5% solvent B (0.1% formic acid in acetonitrile) was used which was increased to 30% solvent B over 90 min (flow rate 300 nL/min). Afterwards the column was washed (95% solvent B as highest acetonitrile concentration) and re-equilibrated to starting conditions. To finally detect the peptides within the samples the nano-HPLC was online coupled to the Quadrupole-Orbitrap Mass Spectrometer using a standard uncoated SilicaTip (ID 20 µm, Tip-ID 10 µM, New Objective, Woburn, MA, USA). A mass range of m/z 300 to 1650 was attained with a resolution of 70000 for full scan prior to 10 high energy collision dissociation (HCD) MS/MS scans of the most intense at least doubly charged ions using a resolution of 17,500 and an NCE energy of 25 %.
The obtained data were analyzed using MaxQuant software (v.1.6.17.0)[4] including the Andromeda search algorithm and using a database containing the IDO1 (Uniprot accession number P14902) or the ubiquitin (Uniprot accession number P0CG48) sequence together with typical contaminants included in the MaxQuant software. The search was performed for full enzymatic trypsin cleavages allowing two miscleavages. For protein modifications carbamidomethylation was chosen as fixed and oxidation of methionine, acetylation of the N-terminus, and GlyGly of lysine as variable modifications. The mass accuracy for full mass spectra was set to 20 ppm (first search) and 4.5 ppm (second search), respectively and for MS/MS spectra to 20 ppm. The false discovery rates for peptide and protein identification were set to 1%. Results were exported into a txt-file and visualized using Prism 6.0 (GraphPad, USA). Detailed analysis procedure is described by Cox *et al.*[4]

*Cellular Thermal Shift Assay in Intact Cells (InCell CETSA)*[6]

SKOV-3 cells were seeded into T-75 flask (or a 6-well plate for the isothermal dose-response experiment) 48 h prior to compound addition. On day 3, the indicated iDeg-1 concentrations or DMSO as a control were added for 1 h and incubated at 37 °C and 5 % $CO_2$ prior to cell detachment by means of trypsin. Cells were washed twice with PBS (4 °C, 300 x g). Afterwards, the cell pellet was resuspended in PBS and equally divided into ten different PCR samples tubes that were subjected to the gradient heat treatment (3 min) by means of the Mastercycler ep gradient (37, 40.8, 44.1, 47.4, 50.0, 53.4, 55.7, 56.3, 61.0 and 63.7°C).

For the isothermal dose-response of iDeg-1, cells were heat-treated at a constant temperature of 50 °C. Afterwards, SKOV-3 cells were placed on ice and cell lysis was performed by adding 0.4 % NP-40 prior to freeze-thaw lysis. After cell lysis, samples were centrifuged at 100,000 x g at 4 °C for 20 min to separate denaturated protein. Supernatants were collected and subjected to an SDS-PAGE and immunoblotting followed by immunostaining for IDO1. Obtained band intensities were quantified using Image Lab software and normalized to the intensities of the bands at 37°C. Melting temperatures in presence or absence of the test compound were determined via non-linear regression using Prism 6.0 (GraphPad, USA).

*Protein Translation Assay in Cells*

BxPC3 cells (100,000 cells/well) were seeded in a 96-well plate and incubated at 37 °C and 5 % $CO_2$ overnight prior to addition of 25 ng/mL IFN-γ and the respective compounds for 24 h. Afterwards, cells were washed three times with pre-warmed PBS followed by Protein translation analysis using the Click-iT HPG Alexa Fluor 488 Protein Synthesis Assay kit (Thermo Fisher). Therefore, fresh medium containing the respective compounds as well as 50 µM L-homopropargylglycine (L-HPG) to tag newly synthesized proteins were added to the cells followed by incubation for 45 min at 37 °C and 5 % $CO_2$. Next, cells were fixed using 4 % *para*-formaldehyde and L-HPG was fluorescently label with Alexa 488 reagent provided by the assay kit using a click reaction as described by the manufacturer (Click-iT HPG Alexa Fluor 488 Protein Synthesis assay, Thermo Fisher). Cell nuclei were stained using the nuclei stain provided in the assay kit. Images were acquired at 10 x magnification using the microscope Axiovert 200M (Zeiss). Images were analyzed by means of MetaXpress software. The L-HPG-mediated fluorescence intensity as well as a cell count analysis were performed and the obtained data were normalized to the cell count and visualized by Fiji ImageJ. Data were plotted using Prism 6.0 (GraphPad, USA).

*In vitro IDO1 Translation Assay*

Full length IDO1 cDNA was cloned into the pT7CFE vector (pT7CFE1-IDO1) and utilized in the *in vitro* translation assay by means of 1-Step Human Coupled DNA IVT kit (Thermo Fisher). To obtain proteins required for protein translation, HeLa cell lysate was prepared according to the manual of the manufacturer. Obtained lysates were combined with pT7CFE1-IDO1 plasmid and supplemented with the provided reagents of the 1-Step Human Coupled DNA IVT kit (Thermo Fisher). The reaction mixture was incubated for 3 h at 30 °C followed by immunoblotting for IDO1.

*Cloning and Purification of Recombinant IDO1 Protein and KLHDC3-EloB/C Complex*

Full length human IDO1 cDNA was cloned into the pGEX6p-2rbs vector using EcoRI and SalI restriction sites. For protein translation assays, full length IDO1 cDNA was cloned into the pT7CFE vector (pT7CFE1-IDO1). Human recombinant IDO1 protein was expressed in *E.coli* according to Littlejohn *et al.*[7] yielding pure and active IDO1 protein. IDO1 protein expression in *E.coli* was induced by the addition of 200 μM IPTG and cells were incubated at 18°C overnight. Harvested cells were resuspended in lysis buffer, disrupted by sonication and the lysate was centrifuged at 13,000 x g for 35 min at 10 °C. The supernatant was applied to a GST Trap HP column to enrich GST-tagged IDO1 equilibrated in 50 mM Tris-HCl, 150 mM NaCl and 1 mM DTE, pH 7.0. To obtain untagged full length IDO1, GST was cleaved from IDO1 overnight using PreScission protease. Eluted IDO1 protein was further purified using a size exclusion column (26/60 G75 HiLoad) to yield pure rhIDO1.

For hIDO1crystallization, a truncated version (5-400) of IDO1 was cloned into the pGEX6p-2rbs vector using the EcoRI and SalI restriction sites. Heme was removed by overnight incubation with 166 mM 2-mercaptoethanesulfonate (MESNA) to obtain apo-IDO1. Heme occupancy was determined by UV/VIS spectrophotometry. For the ubiquitination assays, IDO1 (residues 5-C) was cloned into a pRSF Duet based vector with a N-terminal His-TEV tag and purified as described above.

Expression constructs for KLHDC3-EloB/C (both monomeric and tetrameric versions) were previously described[8]. Briefly, KLHDC3, EloB, and His-TEV-EloC were first cloned into pLib via Gibson assembly. Cassettes were then generated via PCR as described[9] and Gibson assembled into pBig1a to generate a single vector for co-expression of all components. Proper assembly into pBig1a was confirmed by PmeI and SwaI restriction digestion.

KLHDC3-EloB/C was expressed in Hi5 insect cells with an N-terminal 6XHis tag on EloC as previously described[8]. Tagged proteins were purified from cell lysates by Ni affinity chromatography. Nickel elutions were directly loaded onto a HiTrap Q-HP column and eluted with a gradient of 2 %-50 % NaCl. Protein containing fractions were pooled, concentrated, and further purified by size exclusion chromatography in 25 mM HEPES, 200 mM NaCl, 1 mM DTT pH 7.5.

*<u>Protein Modifications</u>*

To introduce a cysteine for fluorescent labeling of UB and K0UB we mutated the protein kinase A site in the pGEX2TK backbone converting the PKA site from RRASV to RRACV[10]. UB or UBK0 purified from this expression construct was labeled with AlexaFluor-647-Maleimide or Fluorescein-5-Maleimide respectively as previously described[10]. Briefly, DTT was added to UB or UBK0 at a final concentration of 10 mM and incubated on ice for 20 minutes to completely reduce cysteines for labeling. DTT was removed by buffer exchange over a NAP-5 column (GE Healthcare) in labeling buffer (25 mM HEPES, 200 mM NaCl). Labeling reactions consisted of UB or UBK0 at 150 μM final concentration and were initiated by the addition of 600 μM AlexaFluor-647-Maleimide or Fluorescein-5-Maleimide (4X excess over labeling target and <5% final DMSO concentration). Reactions were incubated at room temperature for 2 h and quenched by the addition of DTT to 10 mM. Quenched reactions were desalted over a PD-10 column in labeling buffer containing 1mM DTT to remove unreacted probe. Desalted protein was concentrated and further purified over a Sephadex SD75 column.

*IDO1 Enzymatic Assay*

Full-length rhIDO protein (1 µM) in 50 mM potassium phosphate buffer (50 mM $KH_2PO_4$ and 50 mM $K_2HPO_4$, pH 6.5) was incubated with the respective compounds for 40 min at 37 °C prior to the addition of 10 mM ascorbic acid, 10 µM methylene blue, 2 mM Trp and 100 µg/mL catalase (Sigma-Aldrich, C100). Samples were incubated for 60 min followed by the addition of trichloroacetic acid to a final concentration of 7 % and further incubation for 30 min at 37 °C. Kyn levels were detected by the addition of an equal volume of 2 % (w/v) *p*-DMAB in acetic acid and absorbance was measured at 492 nm and 650 nm as a background control. Values are presented relative to the DMSO control. Dose-response curves and $IC_{50}$ values were generated and fitted with GraphPad Prism 9.2.0 (GraphPad software, USA) using four-parameter variable slope non-linear regression curve fit.

*Differential Scanning Fluorimetry (nanoDSF)*

Full-length rhIDO1 (10 µM) in 100 mM potassium phosphate buffer (pH 6.5) was incubated with the respective compound or DMSO as a control at 37 °C for 3 h. Afterwards, the protein solution was loaded onto a capillary followed by the measurement of fluorescence intensities at 350 nm and 330 nm from 20 °C up to 90 °C using the Prometheus Panta device (NanoTemper® Technologies, DE). Melting curves were visualized by GraphPad Prism 9.2.0 (GraphPad software, USA).

*UV/VIS analysis*

Full-length rhIDO1 (10 µM) in 100 mM potassium phosphate buffer (pH 6.5) was incubated with the respective compounds or DMSO as a control for 3 h at 37°C followed by a centrifugation step for 10 min at 1000 x g. Subsequently, the UV/VIS spectra of the samples were monitored between 250 to 650 nm in 1 nm increments using UV transparent microplates (UV-STAR®, Greiner AG, AT) by the Spark® multimode microplate reader (Tecan, Austria).

*Isothermal Titration Calorimetry*

All experiments were performed on a MicroCal AutoITC200. KLHDC3-EloB/C was buffer exchanged by desalting over a NAP5 column in ITC buffer (25 mM HEPES, 150 mM NaCl, 1 mM 2-mercaptoethanol, pH 7.5). Lyophilized peptides were dissolved in ITC buffer to a final concentration of 1 mM. Titrations contained KLHDC3-EloB/C in the sample cell at 24 µM and 240 µM test peptide in the syringe. Titrations were performed at 25°C with one injection of 0.4 µL, followed by 12 injections of 3 µL. Experiments were performed n = 2 times, and data evaluated with MicroCal PEAQ software.

*Co-crystallization of Apo-IDO1 Protein with iDeg-2*

10 mg/mL (225 µM) of truncated IDO1 protein was incubated with 450 µM iDeg-2 (1.8% (v/v) DMSO) in binding buffer (25 mM Tris pH 8.0, 100 µM tris(2-carboxyethyl)phosphine (TCEP) at 37°C for 2 h. 0.4 µL of the rhIDO1:iDeg-2 solution was mixed with 0.2 µL of precipitant solution in a sitting drop setup (iQ plates, SPT Labtech, UK) at 20 °C. Crystals were obtained in wells with precipitant solution containing 39 % (v/v) PEG200 in 100 mM MES pH 6.95. Crystals appeared in one day as thin bars. After 5 days, crystals were harvested and frozen in liquid nitrogen using the precipitant solution as cryoprotectant. X-ray diffraction data collection was carried out at ESRF (Grenoble, FR) on beamline ID30B. Data were processed with XDS and scaled with XSCALE. The structure was solved by molecular replacement with Phaser[11] from the Phenix suite[12] using as initial model the coordinates of PDB 8ABX. Iterative refinement cycles were carried out using phenix.refine and COOT[13]. The figures were generated using the PyMOL Molecular Graphics System (Version 2.5.4, Schrödinger, LLC). LigPlot+[14] was used to analyze protein ligand interactions. The PDB ID code, data collection and refinement statistics are presented in Extended Data Table 2. The crystal structure of IDO1 with iDeg-2 was deposited in the Protein Data Bank (PDB) with the accession number 9FOH.

*Ubiquitination Assays*

The use of pulse-chase assays allowed comparing the paths of UB transfer starting from UBE2R2 or UBE2D2. First, UBE2R2 or UBE2D2 was pulse-labeled by incubating a mixture of UBA1 (0.3 µM), E2 (10 µM), and fluorescently labeled UB or KOUB (15 µM) in 25 mM HEPES, 100 mM NaCl, 100 mM $MgCl_2$, 2 mM ATP, pH 7.5 at room temperature for 13 min. Pulse-loading reactions were quenched by the addition of EDTA to 50 mM and incubated on ice for 5 min. Ubiquitination chase reactions consisted of mixing the E2~*UB thioester conjugate (0.4 µM final concentration) with the indicated pre-equilibrated NEDD8~CUL2-

RBX1-KLHDC3-EloB/C (0.3 µM final concentration) with a threefold excess of IDO1 variants in 25 mM HEPES, 100 mM NaCl, 50 mM EDTA, 0.5 mg/mL BSA, pH 7.5 at room temperature. Experiments following loading of apo-IDO1 with small molecules consisted of incubating IDO1 (15 µM final concentration) with a twofold excess of the indicated test molecules for 1.5 h at 37°C. Mixtures were pelleted for 10 min at 14,000 rpm to remove insoluble material. A UV-Vis spectrum of the resulting supernatant confirmed heme incorporation and the samples were directly diluted into ubiquitination reactions. Reactions were quenched at the indicated times with 2X SDS-PAGE sample buffer and separated on 4-12 % Bis-Tris gradient gels and scanned for fluorescence on a Typhoon imager. All experiments, except where indicated, utilized a monomeric version of KLHDC3-EloB/C.

To assess the relative binding strength of apo-IDO1, or small molecule bound variants of IDO1, we employed a competition pulse-chase ubiquitination assay. First, KLHDC3-EloB/C (50 nM) was equilibrated at room temperature 15 min with 200 nM IDO1$^{C\text{-}deg}$ peptide and variable concentrations of full-length apo-IDO1, iDeg-6-bound IDO1, or linrodostat-bound IDO1. Chase reactions were then initiated by the addition of the pre-formed fluorescent UBE2R2~UB thiolester conjugate (100 nM final concentration). Reaction aliquots were removed at 30 s and quenched with 2X SDS-PAGE sample buffer, separated on 4-12% Bis-Tris gradient gels, and scanned for fluorescence on a Typhoon imager. Data were quantified and visualized as the loss of UB~IDO1$^{C\text{-}deg}$ ubiquitination upon increasing concentrations of full-length IDO1 variants, yielding apparent $IC_{50}$ values for KLHDC3 binding by the differing IDO1 full-length variants

*Reporter and sgRNA Generation*

The N-terminal stability reporter was generated starting from the previously reported C-terminal stability vector.[15,16] The backbone was digested using SalI and MluI (NEB), while the mCherry and TagBFP inserts were amplified using Phusion® High-Fidelity DNA Polymerase following the manufacturer's instructions and using the primers indicated in Extended Data Table 3. Fragments were subsequently assembled using NEBuilder ® HiFi DNA Assembly Cloning Kit. The MluI restriction site was maintained and utilized for the insertion of the various IDO1 constructs amplified with the primers indicated in Extended Data Table 3 and again assembled using the NEBuilder ® HiFi Kit. As starting material for the genetic versions of IDO1, the Addgene plasmid #187026 (pcDNA3.1-IDO1-p2a-eGFP gifted by Geert van den Bogaart[17]) was used. The custom-made sgRNA library targeting elements of the ubiquitin proteasome system[18] generated as previously described in Hsia et al[15]. Single sgRNAs were designed using the VBC score[19] and ordered as custom oligos (Sigma). After phosphorylation (T4 PNK, NEB) and annealing of the oligos, the NEBridge® Golden Gate Assembly Kit

(BsmBI-v2) was used for insertion of the respective sgRNAs (sequences in Extended Data Table 3). sgRNAs were cloned into pLenti-U6-sgRNA- IT-EF1αs-THY1.1-P2A-NeoR.[15] Insertion of the correct DNA sequence was verified by Sanger Sequencing (Microsynth, Austria).

*Virus Production, Transductions and Cell Line Generation for CRISPR-Cas9-based Screen*

Lenti-X cells were transfected at 70 % confluence with the reporter or sgRNA plasmids as well as the two packaging plasmids (pCMVR8.74 helper, and pMD2.G envelope) using polyethylenimine (PEI MAX® MW 40,000, Polysciences). pCMVR8.74 was a gift from Didier Trono (Addgene plasmid # 22036; http://n2t.net/addgene:22036 ; RRID:Addgene_22036). pMD2.G was a gift from Didier Trono (Addgene plasmid # 12259; http://n2t.net/addgene:12259 ; RRID:Addgene_12259). The virus was collected and filtered using a 0.45 µm poly-ethersulfone filter. One million KBM7 iCas9 cells or KBM7 iCas9 IDO1 N-terminal stability reporter cells per 2 mL were transduced with varying volumes of virus solution and incubated with 8 µg/mL polybrene for 24 h prior to cell expansion. Cells were selected either via FACS using a CytoFLEX SRT Benchtop Cell Sorter (for reporter cell lines: BFP and mCherry positive cells, see Extended Data Fig. 7c) or using G418 (1 mg/mL, Gibco) for sgRNA harboring cell lines, which was added 72 h after the transduction. Knockout (KO) cell lines were generated by selection of the respective sgRNA (sgAAVS1 = CTRL or sgRNA1/2 KLHDC3 for KO1/KO2) for at least 14 days with G418 followed by subsequent induction with doxycycline (0.4 µg/mL, PanReac AppliChem) and cell recovery.

*Flow-cytometric IDO1 Reporter Assay*

IDO1 KBM7 reporter cells were treated for the times, compounds and concentrations indicated in the respective figure legends or as specified in the text. All compounds were diluted starting from DMSO stock solutions. BFP and mCherry levels were measured using an LSR Fortessa (BD Biosciences) with the BD FACSDiva software (v9.0). To quantify the changes across conditions the mean BFP and mCherry values were exported after gating for singlets and reporter positive cells (see gating strategy in Fig S7a,c) utilizing FlowJo 10.6.2.. Normalization was performed either against the respective *wt IDO1* reporter cells treated with DMSO (Fig. 5h), their respective co-treatment (Fig. 4f, Extended Data Fig. 6d) or for each respective genotype (Fig. 4b,i, Fig. 5i, Fig. 6f,i, Extended Data Fig. 6e).

*FACS-based CRISPR Cas9 Screen*

The screen, library preparation and sequencing analysis was performed as previously described.[15] In brief, 120 million KBM7 iCas9 IDO1 N-terminal stability reporter cells were transduced at an MOI of ~ 0.12 sgRNA positive cells (> 1,000-fold library representation, Fig. 7e) with the customized ubiquitin focused library. Transduced cells were selected with G418 (1 mg/mL, Gibco) for 14 days, expanded and Cas9 expression was induced with doxycycline (0.4 µg/mL, PanReac AppliChem). Three days after Cas9 induction, 50 million cells per condition and replicate were treated with DMSO, iDeg-1 or iDeg-2. The experiment was performed in two biological replicates. Cells were washed with PBS, stained with APC anti-mouse Thy1.1 antibody (1:400, 202526, BioLegend) and Human TruStain FcX Fc Receptor Blocking Solution (1:1000, 422302, BioLegend) for 5 min at 4°C. Subsequently cells were fixed with BD Fixation buffer 4 % (Thermo Scientific™ Pierce™) for 45 min at 4°C. All steps were performed protected from light. Cells were washed twice with PBS and stored in FACS buffer (PBS, 5 % FBS and 1 mM EDTA).

For sorting, cells were strained using a 35-µm nylon mesh and sorted on a BD FACSAria Fusion (70 µm nozzle, BD Biosciences) with the BD FACSDiva software (v8.0.2). Singlets, Cas9 and sgRNA positive cells were selected following the scheme shown in Extended Data Fig. 7a-d. Next, fractions with different levels of BFP were enriched: the 5 % highest or lowest BFP expressing cells were used for the respective HIGH and LOW fractions, and 30 % of the population's center was selected for the MID fractions. For each replicate, cells corresponding to at least a 1,000-fold library representation were sorted.

To quantify sgRNA levels per each fraction and sample next generation sequencing (NGS) libraries were prepared. Genomic DNA was isolated by cell lysis (10 mM Tris-HCl, 150 mM NaCl, 10 mM EDTA, 0.1 % SDS) and proteinase K treatment (New England Biolabs) overnight at 55°C shaking at 1200 rpm, followed by 2 h of DNAse-free RNAse (Thermo Fisher Scientific) incubation at 37°C. Next, DNA was extracted by two rounds of phenol extraction and subsequent isopropanol precipitation. Barcoded NGS libraries for each sorted population were generated using a two-step PCR protocol using AmpliTaq Gold polymerase (Invitrogen). The resulting PCR products were purified after each step using Mag-Bind TotalPure NGS beads (Omega Biotek). The final NGS libraries were pooled and sequenced on a NovaSeq 6000 platform (Illumina).

Sequencing reads were trimmed using fastx-toolkit (v0.0.14), aligned using Bowtie2 (v2.4.5) and quantified using featureCounts (v2.0.1). The utilized workflows are available at https://github.com/ZuberLab/crispr-process-nf/tree/566f6d46bbcc2a3f49f51bbc96b9820f408ec4a3 and https://github.com/ZuberLab/crispr-mageck-nf/tree/c75a90f670698bfa78bfd8be-

786d6e5d6d4fc455. To calculate gene-level enrichment, the sorted populations (HIGH or LOW) were compared to the MID populations using the median-normalized read counts.

#### Chemistry

##### *General Experimental Procedures*

Reactions were performed in heat gun dried flasks under an argon atmosphere. All reagents purchased from commercial sources were used directly without further purification.
Thin-layer chromatography was performed on pre-coated silica gel F-254 plates from Merck. The spots were visualized by UV light and/or by staining of the TLC plate with potassium permanganate stain followed, if necessary, by heating with a heat gun. For column chromatography, silica gel 60 from Merck with a particle size of 0.040-0.063 mm was used.
$^{1}$H- and $^{13}$C-NMR spectra were recorded on Bruker AV 400 Avance III HD (400 MHz), Bruker AV 500 Avance III HD (500 MHz), Bruker AV 600 Avance III HD (600 MHz) and Bruker AV 700 Avance III HD (700 MHz) spectrometers at room temperature.
HPLC-MS analyses were performed with an Agilent 1100 Series connected to a Thermo LCQ Advantage mass spectrometer using a C18 HPLC column 3 µm from Macherey Nagel.
High resolution mass spectra were recorded on a QLTOrbitrap mass spectrometer coupled to an Acceka HPLC-System (HPLC column: Hypersyl GOLD, 50 mm × 1 mm particle size, 1.9 µm).

For the synthesis of iDegs, (1*R*)-(−)-myrtenal (**1**) was subjected to a 1,3-dipolar cycloaddition with the azomethine ylide generated *in situ* from *N*-(methoxymethyl)-*N*-(trimethylsilylmethyl) benzylamine by treatment with trifluoroacetic acid to yield the corresponding pyrrolidine cycloadduct (**2**) (Supporting Scheme 1a) analogous to previously reported comparable cycloadditions (Mutti *et al.* and Sheffler. *et al.*)[13-14]. Subsequently, the aldehyde (**2**) was reduced to the corresponding alcohol **3** (Supporting Scheme 1b). For synthesis of iDeg-1 the alcohol was further functionalized as urethane, followed by *N*-debenzylation and sulfonamide formation (Supporting Scheme 2). For synthesis of iDeg-2, -3, and -4, the *N*-benzyl protecting group was removed from **3** to yield amine **7** followed by sulfonamide formation (⇨ **8, 9**) and functionalization of the alcohol as urethanes **10** (iDeg-2) and **11** (Supporting Scheme 3). Subsequently, the aryl iodide in iDeg-2 (**10)** and in urethane **11** were subjected to a Pd(0) mediated Sonogashira coupling reaction to yield **12** (iDeg-3) and **13** (iDeg-6) respectively (Supporting Scheme 3).

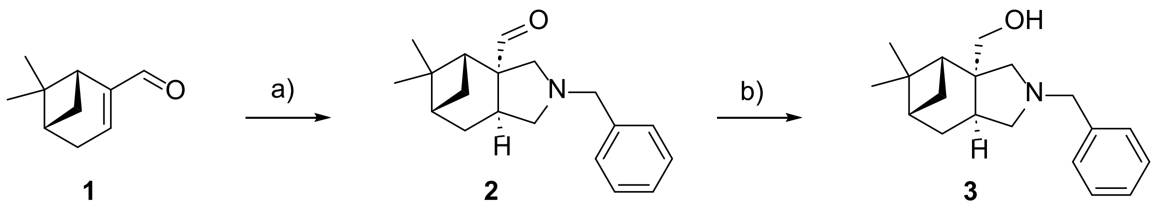

**Supporting Scheme 1**: **a**) *N*-(Methoxymethyl)-*N*-(trimethylsilylmethyl) benzylamine, TFA, DCM, 0 °C to rt, 16 h; **b**) $NaBH_4$, MeOH, 0 °C to rt, 1 h

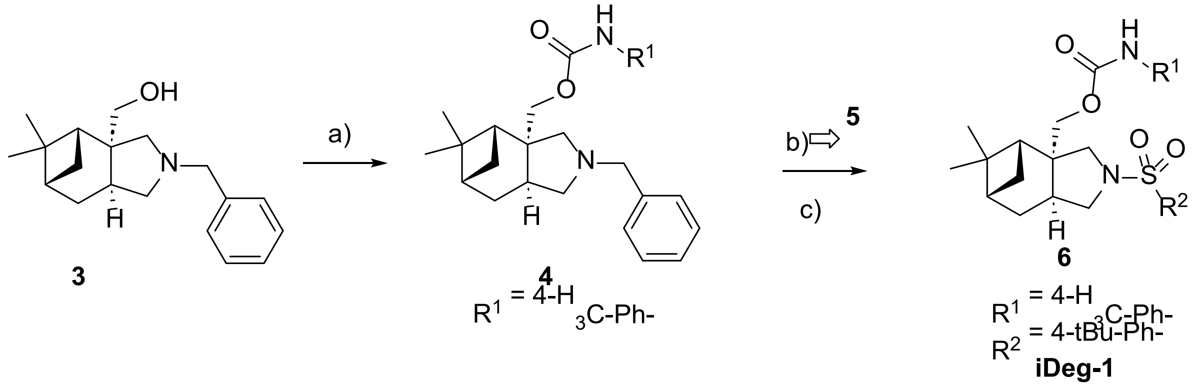

**Supporting Scheme 2**. a) $R^1$NCO, $NEt_3$, THF, rt, 2 h; b) Pd/C, $NH_4HCO_2$, MeOH, 60 °C, 1 h or Pd/C, $H_2$, EtOH/THF, rt, 18 h; c) $R^2SO_2Cl$, $NEt_3$, DCM, 0 °C to rt, 2 h.

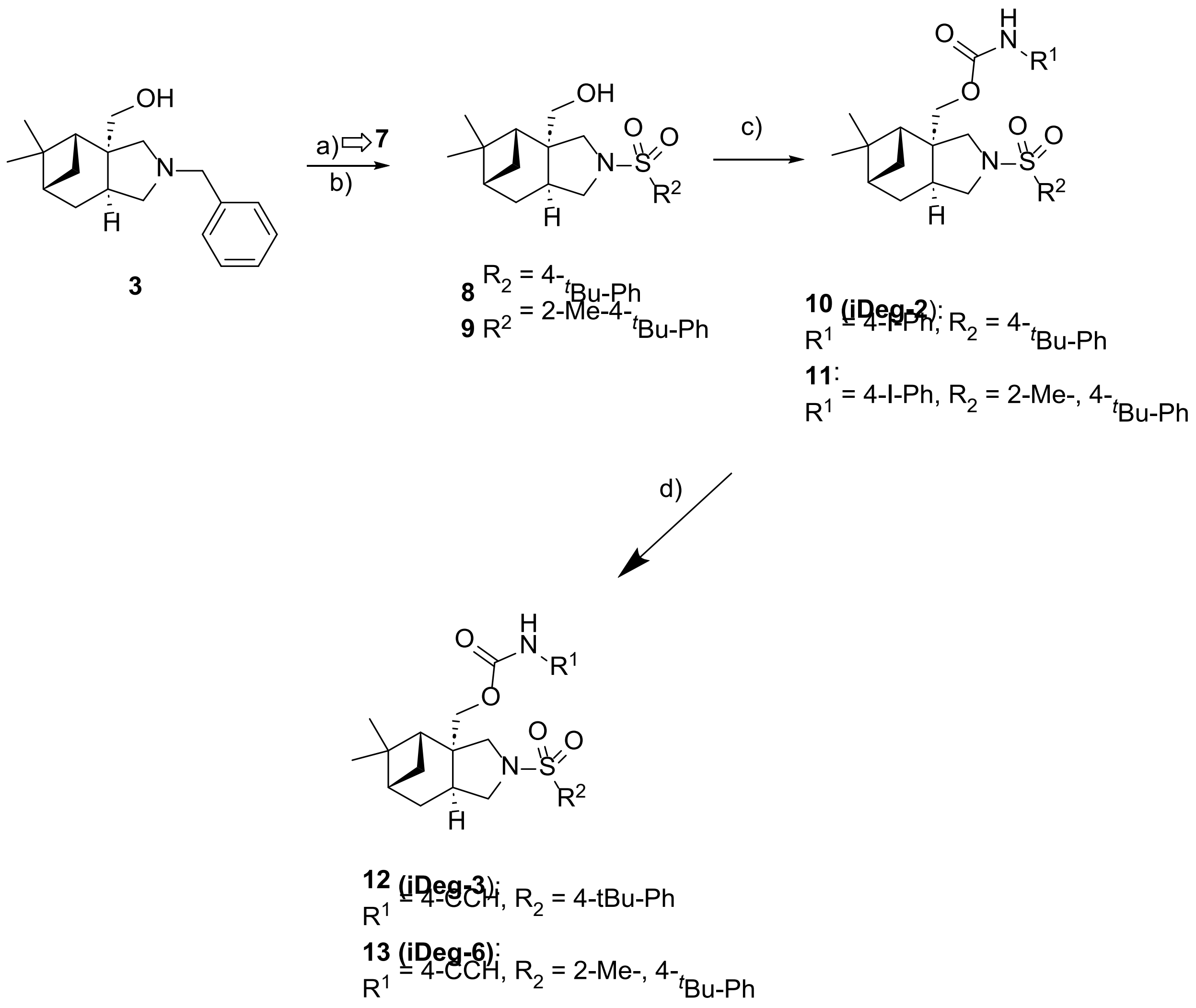

**Supporting Scheme 3.** a) Pd/C, $H_2$, MeOH, rt, 16 h; b) $R^2SO_2Cl$, $NEt_3$, DCM, 0 °C to rt, 2 h; c) $R^1NCO$, DBU, THF, rt, 18 h; d) $Me_3SiCCH$, $NEt_3$, THF, $Pd(PPh_3)_4$, then TBAF.

*General procedure A: carbamate formation*

To a solution of the alcohol (1.0 equiv.) in THF (0.2 M) was added $NEt_3$ (2.0 equiv.) and the respective isocyanate (1.3 equiv.) and the solution was stirred at room temperature for 2 h. The solvent was removed under reduced pressure. Column chromatography afforded the carbamate.

*General procedure B: benzyl group cleavage*

**B1:**

To a degassed solution of the *N*-benzyl protected amine (1.0 equiv.) in MeOH (0.1−0.2 M) was added ammonium formate (5.0 equiv.) and Pd/C (10 wt.%) and the suspension was stirred at 60 °C for 1 h. After cooling to room temperature, the reaction mixture was filtered over Celite, washed with EtOAc and the solvent was removed under reduced pressure to give the amine.

**B2:**

To a degassed solution of the *N*-benzyl-protected amine (1.0 equiv.) in MeOH (0.1 M) was added Pd/C (10 wt.%) and the suspension was stirred under a $H_2$ atmosphere for 16−22 h. The reaction mixture was filtered over Celite, washed with EtOAc and the solvent was removed under reduced pressure to give the amine.

*General procedure C: sulfonamide formation*

To a solution of the amine (1.0 equiv.) in dichloromethane (0.2 M) at 0 °C was added $NEt_3$ (2.0 equiv.) and the respective sulfonyl chloride (1.2 equiv.) and the solution was stirred at room temperature for 2 h. The solvent was removed under reduced pressure. Column chromatography afforded the sulfonamide.

*General procedure D: carbamate formation*

To a solution of the alcohol (1.0 equiv.) in THF (0.1−0.2 M) was added DBU (1.5 equiv.) and the respective isocyanate (1.2 equiv.) and the solution was stirred at room temperature for 18-22 h. The solvent was removed under reduced pressure. Column chromatography afforded the carbamate.

**(3a*S*,4*R*,6*R*,7a*S*)-2-benzyl-5,5-dimethyloctahydro-3a*H*-4,6-methanoisoindole-3a-carbaldehyde (2)**

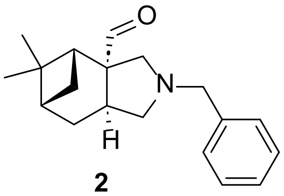

**2**

To a solution of myrtenal (**1**) (2.0 g, 13.3 mmol, 1 eq.) and *N*-(methoxymethyl)-*N*-trimethylsilylmethyl) benzylamine 1 (4.5 mL, 15.9 mmol, 1.2 eq., 90% pure) in 65 mL dry $CH_2Cl_2$ under argon atmosphere in an ice bath at 0°C, a solution of TFA in dry DCM (0.2 mL, 2.6 mmol, 0.2 eq., 0.1 M) was added dropwise over 1 min. The mixture was stirred for 12 h at 5-10°C. Progress of the reaction was monitored by TLC. After complete consumption of the starting material the reaction was quenched by the addition of aqueous saturated sodium bicarbonate and the mixture was stirred for 5 min. The layers were separated and the aqueous layer was extracted with dichloromethane. The combined organic layers were dried over $Na_2SO_4$, filtered, and concentrated under reduced pressure to give a crude residue, which was purified by chromatography (PE/EtOAc 4:1). The product was obtained as colourless oil (2.79 g, 9.84 mmol, 74 % yield).

**$^1$H NMR** (400 MHz, $CDCl_3$): δ 9.69 (s, 1H), 7.38 – 7.13 (m, 5H), 3.65 (d, *J* = 13.1 Hz, 1H), 3.50 (d, *J* = 13.1 Hz, 1H), 3.18 (t, *J* = 8.4 Hz, 1H), 2.92 (dtd, *J* = 10.7, 7.8, 3.0 Hz, 1H), 2.69 (d, *J* = 10.2 Hz, 1H), 2.43 (d, *J* = 10.2 Hz, 1H), 2.35 – 2.08 (m, 4H), 1.88 (tt, *J* = 6.0, 3.1 Hz, 1H), 1.69 – 1.57 (m, 2H), 1.24 (s, 3H), 0.69 (s, 3H).

**$^{13}$C NMR** (101 MHz, $CDCl_3$): δ 206.1, 138.9, 128.6, 128.4, 127.1, 66.2, 61.2, 60.9, 59.7, 45.8, 40.5, 38.7, 32.2, 31.0, 26.6, 26.5, 23.7.

**HRMS** (ESI) for $C_{19}H_{26}ON$: calculated 284.2009 $[M+H]^+$, found: 284.2015.

**((3a*S*,4*R*,6*R*,7a*S*)-2-benzyl-5,5-dimethyloctahydro-3a*H*-4,6-methanoisoindol-3a-yl)methanol (3)**

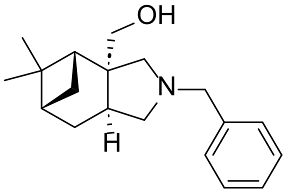

**3**

To a solution of **2** (1.0 g, 3.5 mmol, 1 eq.) in MeOH (10 mL) at 0°C $NaBH_4$ (173 mg, 4.5 mmol, 1.3 eq.) was added in portions. After stirring for 1 h, the reaction was quenched with acetone

(5 mL). The solvents were removed under reduced pressure and the residue was dissolved in water. The aqueous layer was extracted twice with ethyl acetate, the combined organic layers were dried over $Na_2SO_4$ and concentrated under reduced pressure. The crude product was purified by chromatography (PE/EtOAc 4:1) and the product was obtained as colourless oil (0.92 g, 3.22 mmol, 92 % yield)

**$^1$H NMR** (500 MHz, $CDCl_3$) δ 7.32 (d, *J* = 4.5 Hz, 4H), 7.30 – 7.25 (m, 1H), 3.81 (d, *J* = 9.9 Hz, 1H), 3.68 (t, *J* = 10.8 Hz, 2H), 3.39 (d, *J* = 9.9 Hz, 1H), 3.36 (d, *J* = 8.6 Hz, 1H), 2.86 (d, *J* = 9.3 Hz, 1H), 2.59 – 2.52 (m, 1H), 2.50 (d, *J* = 9.5 Hz, 1H), 2.23 – 2.17 (m, 1H), 2.18 – 2.11 (m, 1H), 2.11 (d, *J* = 13.9 Hz, 1H), 1.89 (dq, *J* = 5.9, 3.0 Hz, 1H), 1.75 (dd, *J* = 6.4, 4.9 Hz, 1H), 1.56 (d, *J* = 13.5 Hz, 1H), 1.40 (d, *J* = 10.4 Hz, 1H), 1.22 (s, 3H), 0.95 (s, 3H).
**$^{13}$C NMR** (126 MHz, $CDCl_3$) δ 137.4, 129.0, 128.7, 127.7, 71.4, 66.4, 64.2, 59.7, 51.0, 46.2, 41.3, 38.9, 32.9, 32.1, 27.6, 27.6, 24.4.

**HRMS** (ESI) for $C_{19}H_{28}ON$; calculated 286.2165 $[M+H]^+$, found 286.2167.

$[\alpha]^{20}_D$ = -53.7° ($CHCl_3$, *c* = 1.0 )

**((3a*S*,4*R*,6*R*,7a*S*)-2-benzyl-5,5-dimethyloctahydro-3a*H*-4,6-methanoisoindol-3a-yl)methyl *p*-tolylcarbamate (4)**

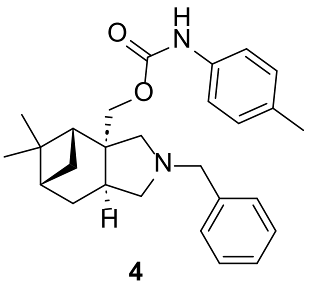

**4**

Following general procedure A: From **3** (100 mg, 0.350 mmol) and *p*-tolyl isocyanate. Purification by column chromatography (PE/EtOAc 9:1) afforded a colorless foam (122 mg, 0.291 mmol, 83 % yield).

**$^1$H NMR** ($CDCl_3$, 500 MHz): δ = 1.03 (s, 3 H, $CH_3$), 1.23 (s, 3 H, $CH_3$), 1.60 (dt, *J* = 3.5, 13.3 Hz, 1 H, CH), 1.82–1.89 (m, 2 H, 2 × CH), 1.95 (t, *J* = 5.7 Hz, 1 H, CH), 2.06–2.19 (m, 2 H, 2 × CH), 2.21–2.28 (m, 1 H, CH), 2.30 (s, 3 H, $CH_3$), 2.34 (d, *J* = 9.4 Hz, 1 H, CH), 2.46–2.52 (m, 1 H, CH), 2.55 (d, *J* = 9.5 Hz, 1 H, CH), ), 2.57–2.62 (m, 1 H, CH), 3.55 (d, *J* = 13.1 Hz, 1 H, $CH_A$), 3.63 (d, *J* = 13.3 Hz, 1 H, $CH_B$), 4.05 (d, *J* = 10.3 Hz, 1 H, $CH_{A'}$), 4.21 (d, *J* = 10.3 Hz, 1 H, $CH_{B'}$), 6.46 ($s_{br}$, 1 H, NH), 7.10 (d, *J* = 8.1 Hz, 2 H, 2 × Ar-H), 7.20–7.39 (m, 7 H, 7 × Ar-H).

$^{13}$**C NMR** ($CDCl_3$, 125 MHz): δ = 20.9, 23.6, 26.7, 27.1, 27.6, 29.9, 34.6, 34.9, 39.5, 40.6, 46.3, 50.5, 64.2, 71.0, 118.7, 126.8, 128.3, 128.6, 129.7, 133.01, 135.5, 139.9, 154.0.

**MS** (ESI) *m/z* (%) = 419.4 (100) $[M+H]^+$.
**HRMS** (ESI) for $C_{27}H_{35}N_2O_2$: calculated 419.2693 $[M+H]^+$, found 419.12692.

**((3a*S*,4*R*,6*R*,7a*S*)-5,5-dimethyloctahydro-3a*H*-4,6-methanoisoindol-3a-yl)methyl *p*-tolylcarbamate (5)**

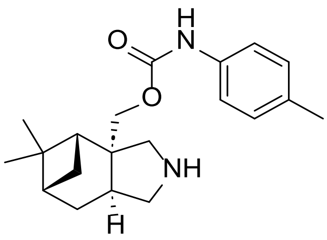

**5**

Following general procedure B1: From **4** (1.0 g, 2.39 mmol) afforded a colorless solid (745.5 mg, 2.27 mmol, 95 % yield).

$^1$**H NMR** ($CDCl_3$, 400 MHz): δ = 7.33 (d, *J* = 8.2 Hz, 2H), 7.08 (d, *J* = 8.2 Hz, 2H), 4.28 (d, *J* = 10.7 Hz, 1H), 4.07 (d, *J* = 10.7 Hz, 1H), 3.75 – 3.61 (m, 1H), 3.41 – 3.28 (m, 2H), 3.11 (d, *J* = 12.3 Hz, 1H), 2.95 (dd, *J* = 11.8, 7.5 Hz, 1H), 2.51 – 2.22 (m, 3H), 2.27 (s, 3H), 2.04 (t, *J* = 5.6 Hz, 1H), 1.96 (d, *J* = 5.4 Hz, 1H), 1.65 (dt, *J* = 13.8, 3.2 Hz, 1H), 1.28 (d, 3H), 1.06 (s, 3H).

**HRMS** (ESI) for $C_{20}H_{29}N_2O_2$: calculated 329,222 $[M+H]^+$, found: 329.224

**((3a*S*,4*R*,6*R*,7a*S*)-2-((4-(*tert*-butyl)phenyl)sulfonyl)-5,5-dimethyloctahydro-3a*H*-4,6-methanoisoindol-3a-yl)methyl *p*-tolylcarbamate(6) (iDeg-1)**

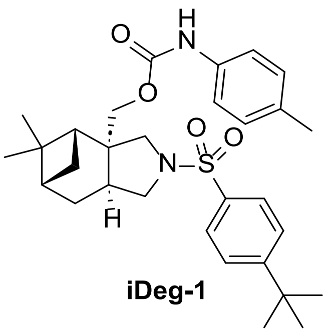

**iDeg-1**

Following general procedure B1: From **4** (110 mg, 0.263 mmol) afforded a colorless solid **5** (81.0 mg, 0.247 mmol, 94 %).

Following general procedure C: From amine **5** (81.0 mg, 0.247 mmol) and 4-*tert*-butylbenzenesulfonyl chloride. Purification by column chromatography (PE/EtOAc 8:1 → 4:1) afforded a colorless foam (115 mg, 0.219 mmol, 89 % yield).

**$^1$H NMR** (600 MHz, $CD_2Cl_2$) δ 7.71 (d, *J* = 8.5 Hz, 2H), 7.56 (d, *J* = 8.6 Hz, 2H), 7.19 (d, *J* = 7.6 Hz, 2H), 7.10 (d, *J* = 7.9 Hz, 2H), 6.36 (s, 1H), 4.06 (d, *J* = 10.7 Hz, 1H), 3.92 (d, *J* = 10.6 Hz, 1H), 3.15 (dd, *J* = 9.2, 7.1 Hz, 1H), 3.05 (d, *J* = 9.9 Hz, 1H), 3.02 (dd, *J* = 9.2, 3.1 Hz, 1H), 2.99 (d, *J* = 9.9 Hz, 1H), 2.33 (ddd, *J* = 10.5, 3.3, 2.1 Hz, 1H), 2.30 (s, 3H), 2.25 (tdd, *J* = 7.4, 4.2, 2.2 Hz, 1H), 2.20 (ddt, *J* = 8.4, 6.2, 3.1 Hz, 1H), 1.98 (dd, *J* = 6.5, 5.0 Hz, 1H), 1.87 (tt, *J* = 5.6, 3.0 Hz, 1H), 1.63 – 1.59 (m, 1H), 1.32 (d, *J* = 10.4 Hz, 1H), 1.30 (s, 9H), 1.22 (s, 3H), 1.01 (s, 3H).

**$^{13}$C NMR** (151 MHz, $CD_2Cl_2$) δ 157.2, 153.7, 135.6, 133.6, 132.4, 129.9, 128.3, 126.4, 119.2, 70.1, 58.5, 57.4, 54.2, 54.0, 53.8, 53.7, 53.5, 50.9, 46.7, 40.6, 39.4, 35.4, 34.7, 34.5, 31.2, 27.4, 27.3, 23.5, 20.8.

**MS** (ESI) *m/z* (%) = 525.2 (100) $[M+H]^+$.

**HRMS** (ESI) for $C_{30}H_{41}N_2O_4S$: calculated 525.2782 $[M+H]^+$, found 525.2785.

$[\alpha]^{20}_D$ = -16.0° ($CHCl_3$, *c* = 0.5)

**((3a*S*,4*R*,6*R*,7a*S*)-2-((4-(*tert*-butyl)phenyl)sulfonyl)-5,5-dimethyloctahydro-3a*H*-4,6-methanoisoindol-3a-yl)methanol (8)**

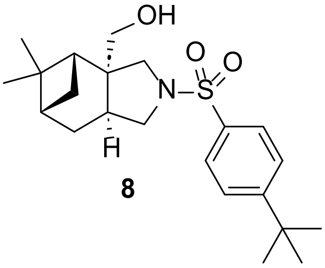

**8**

To a degassed solution of **3** (389 mg, 1.36 mmol) in MeOH (4 mL) was added Pd/C (10 wt%, 40 mg) and the mixture was stirred under a $H_2$ atmosphere for 16 h. The reaction mixture was filtered over Celite, washed with MeOH and the solvent was evaporated to give intermediate **7** a colorless solid (242 mg, 1.24 mmol, 91 % yield) which was used immediately for the following transformation.

To a solution of the solid **7** (242 mg, 1.24 mmol) in THF (10 mL) at 0 °C was added $NEt_3$ (198 µL, 1.49 mmol, 1.2 equiv.) and 4-*tert*-butylbenzenesulfonyl chloride (288 mg, 1.24 mmol, 1.0 equiv.) and the mixture was stirred at 0 °C for 2 h. The solvent was removed under reduced pressure. Column chromatography (PE/EtOAc 4:1 → 2:1) afforded a colorless solid 8 (415 mg, 1.06 mmol, 86 % yield).

**$^{1}$H NMR** (500 MHz, $CDCl_3$) δ 7.72 (d, *J* = 8.6 Hz, 2H), 7.53 (d, *J* = 8.6 Hz, 2H), 3.51 (d, *J* = 10.5 Hz, 1H), 3.42 (d, *J* = 10.6 Hz, 1H), 3.19 (dd, *J* = 9.1, 7.3 Hz, 1H), 3.06 (d, *J* = 9.7 Hz, 1H), 3.01 – 2.95 (m, 2H), 2.31 – 2.23 (m, 1H), 2.21 – 2.14 (m, 2H), 1.96 (dd, *J* = 6.5, 5.0 Hz, 1H), 1.86 (dq, *J* = 5.4, 2.8 Hz, 1H), 1.61 (ddd, *J* = 13.4, 4.4, 2.8 Hz, 1H), 1.35 (s, 9H), 1.31 (d, *J* = 10.4 Hz, 1H), 1.21 (s, 3H), 0.95 (s, 3H).

**$^{13}$C NMR** (126 MHz, $CDCl_3$) δ 156.6, 132.0, 128.1, 126.0, 68.2, 58.5, 56.9, 52.3, 45.8, 40.4, 39.2, 35.3, 34.2, 34.1, 31.2, 27.5, 27.2, 23.7.

**MS** (ESI) *m/z* (%) = 392.2 (100) $[M+H]^+$.
**HRMS** (ESI) for $C_{22}H_{34}NO_3S$: calculated 392.2254 $[M+H]^+$, found 392.2254.

$[\alpha]^{20}_{D}$ = - 25.3° ($CHCl_3$, *c* = 1.0).

**((3a*S*,4*R*,6*R*,7a*S*)-2-((4-(*tert*-butyl)phenyl)sulfonyl)-5,5-dimethyloctahydro-3a*H*-4,6-methanoisoindol-3a-yl)methyl (4-iodophenyl)carbamate (10) (iDeg-2)**

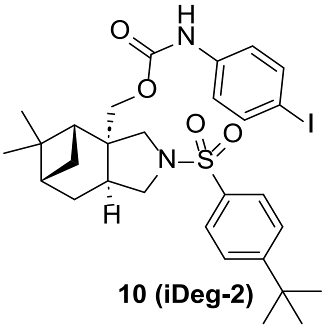

**10 (iDeg-2)**

Following general procedure D: From **8** (20.0 mg, 51.1 μmol) and 4-iodophenyl isocyanate (15 mg, 1.2 eq.). Purification by column chromatography (PE/EtOAc 4:1) afforded a colorless solid (28.0 mg, 44.0 μmol, 86 % yield).

**1H NMR** (500 MHz, $CD_2Cl_2$) δ 7.71 (d, J = 8.5 Hz, 2H), 7.61 (d, J = 8.8 Hz, 2H), 7.56 (d, J = 8.6 Hz, 2H), 7.15 (d, J = 8.3 Hz, 2H), 6.54 (s, 1H), 4.11 (d, J = 10.7 Hz, 1H), 3.93 (d, J = 10.7 Hz, 1H), 3.20 (dd, J = 9.3, 7.1 Hz, 1H), 3.04 – 2.96 (m, 3H), 2.37 – 2.28 (m, 1H), 2.28 – 2.17 (m, 2H), 1.98 (dd, J = 6.5, 4.9 Hz, 1H), 1.87 (s, 1H), 1.63 – 1.58 (m, 1H), 1.31 (s, 9H), 1.28 (s, 1H), 1.22 (s, 3H), 1.01 (s, 3H).

**$^{13}$C NMR** (126 MHz, $CD_2Cl_2$) δ 157.2, 153.4, 138.3, 138.2, 132.5, 128.3, 126.4, 120.9, 86.5, 70.4, 58.5, 57.3, 50.8, 46.6, 40.6, 39.4, 35.4, 34.7, 34.3, 31.2, 27.4, 27.2, 23.5.

**MS** (ESI) *m/z* (%) = 637.0 (100) $[M+H]^+$.
**HRMS** (ESI) for $C_{29}H_{38}IN_2O_4S$: calculated 637.1592 $[M+H]^+$, found 637.1601.$[\alpha]^{20}_D$ = -10.7° ($CHCl_3$, *c* = 1.0).

**((3a*S*,4*R*,6*R*,7a*S*)-2-((4-(*tert*-butyl)phenyl)sulfonyl)-5,5-dimethyloctahydro-3a*H*-4,6-methanoisoindol-3a-yl)methyl (4-ethynylphenyl)carbamate(12) (iDeg-3)**

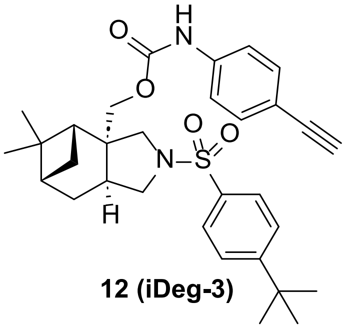

**12 (iDeg-3)**

To a degassed solution of **10** (60.0 mg, 94.3 µmol) in THF (0.5 mL) was added $NEt_3$ (65 µL, 0.471 mmol, 5.0 equiv.), trimethylsilylacetylene (26 µL, 0.189 mmol, 2.0 equiv.), CuI (3.6 mg, 18.9 µmol, 0.2 equiv.) and Pd $(PPh_3)_4$ (10.9 mg, 9.43 µmol, 0.1 equiv.) and the mixture was stirred at room temperature for 22 h. The reaction mixture was poured into sat. aq. NaCl solution, extracted with dichloromethane (2 × 20 mL), dried over $MgSO_4$ and the solvent was removed under reduced pressure. Column chromatography (PE/EtOAc 6:1) afforded a beige colored solid (50.0 mg).

To a solution of the solid in THF (0.5 mL) was added TBAF (1 M in THF, 99 µL, 98.9 µmol, 1.2 equiv.) and the mixture was stirred at room temperature for 1 h. The solvent was removed under reduced pressure. Column chromatography (PE/EtOAc 5:1) afforded a beige colored oil (25.0 mg, 46.8 µmol, 50 % yield over 2 steps).

**$^1$H NMR** (500 MHz, $CD_2Cl_2$) δ 7.72 (d, J = 8.6 Hz, 2H), 7.56 (d, J = 8.6 Hz, 2H), 7.43 (d, J = 8.7 Hz, 2H), 7.33 (d, J = 8.4 Hz, 2H), 6.63 (s, 1H), 4.12 (d, J = 10.7 Hz, 1H), 3.94 (d, J = 10.7 Hz, 1H), 3.21 (dd, J = 9.3, 7.1 Hz, 1H), 3.09 (s, 1H), 3.05 – 2.97 (m, 3H), 2.32 (tdd, J = 13.0, 3.1, 2.0 Hz, 1H), 2.26 (tt, J = 7.0, 3.6 Hz, 1H), 2.20 (dtd, J = 10.4, 6.3, 2.0 Hz, 1H), 1.98 (dd, J = 6.5, 5.0 Hz, 1H), 1.88 (tt, J = 5.7, 3.1 Hz, 1H), 1.63 – 1.59 (m, 1H), 1.31 (s, 9H), 1.28 (d, J = 9.9 Hz, 1H), 1.23 (s, 3H), 1.01 (s, 3H).

**$^{13}$C NMR** (126 MHz, $CD_2Cl_2$) δ 157.2, 153.3, 142.6, 138.9, 133.3, 132.5, 128.3, 126.4, 118.5, 117.1, 83.6, 76.8, 70.4, 58.5, 57.3, 50.8, 46.6, 40.6, 39.4, 35.5, 34.7, 34.3, 31.2, 27.4, 27.2, 23.5.

**MS** (ESI) *m/z* (%) = 535.2 (100) $[M+H]^+$.

**HRMS** (ESI) for $C_{31}H_{39}N_2O_4S$: calculated 535.2625 $[M+H]^+$, found 535.2627.

$[\alpha]^{20}_D$ = -13.3° ($CHCl_3$, *c* = 0.5).

**((3a*S*,4*R*,6*R*,7a*S*)-2-((4-(*tert*-butyl)-2-methylphenyl)sulfonyl)-5,5-dimethyloctahydro-3a*H*-4,6-methanoisoindol-3a-yl)methanol (9):**

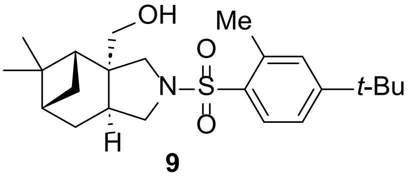

**9**

To a solution of the solid **7** (58.6 mg, 0.3 mmol) in THF (10 mL) at 0 °C was added $NEt_3$ (1.2 equiv.) and 4-(*tert*-butyl)-2-methylbenzenesulfonyl chloride (74.0 mg, 1.0 eq.) and the mixture was stirred at 0 °C for overnight. The solvent was removed under reduced pressure. Column chromatography (PE/EtOAc 4:1 → 2:1) afforded a colorless solid (80 mg, 0.2 mmol, 66 % yield).

**$^1$H NMR** (500 MHz, $CDCl_3$) δ 7.83 (d, J = 8.1 Hz, 1H), 7.33 – 7.28 (m, 2H), 3.57 (d, J = 10.6 Hz, 1H), 3.50 (d, J = 10.6 Hz, 1H), 3.42 (dd, J = 9.3, 7.0 Hz, 1H), 3.25 (d, J = 9.8 Hz, 1H), 3.03 (dd, J = 9.4, 3.7 Hz, 1H), 2.99 (d, J = 9.7 Hz, 1H), 2.66 (s, 3H), 2.27 (ddt, J = 12.1, 7.6, 2.1 Hz, 1H), 2.23 – 2.16 (m, 2H), 2.00 – 1.94 (m, 1H), 1.87 (dt, J = 5.4, 2.7 Hz, 1H), 1.59 (d, J = 3.2 Hz, 1H), 1.33 (s, 9H), 1.31 (d, J = 10.5 Hz, 1H), 1.22 (s, 3H), 0.97 (s, 3H).

**$^{13}$C NMR** (126 MHz, $CDCl_3$) δ 156.6, 138.0, 132.6, 130.4, 129.9, 123.2, 68.3, 58.0, 56.3, 52.4, 45.8, 40.5, 39.2, 35.0, 34.3, 34.1, 31.2, 27.6, 27.3, 23.8, 21.3.

**HRMS** (ESI) for $C_{23}H_{36}NO_3S$: calculated 406.2410 $[M+H]^+$, found 406.2408.

$[\alpha]^{20}_D$ = -88.3° ($CHCl_3$, *c* = 0.5)

**((3a*S*,4*R*,6*R*,7a*S*)-2-((4-(*tert*-butyl)-2-methylphenyl)sulfonyl)-5,5-dimethyloctahydro-3a*H*-4,6-methanoisoindol-3a-yl)methyl (4-iodophenyl)carbamate (11):**

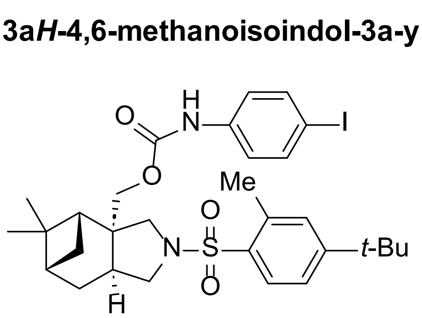

**11**

Following general procedure D: From **9** (60.84 mg, 0.15 mmol) and 1-iodo-4-isocyanatobenzene (73.5 mg, 1.5 eq.). Purification by column chromatography (PE/EtOAc 20:1 to 10:1) afforded a colorless solid **11** (69.6mg, 0.11 mmol, 71 % yield).

**1H NMR** (500 MHz, $CDCl_3$) δ 7.85 (d, J = 8.7 Hz, 1H), 7.60 (d, J = 8.8 Hz, 2H), 7.33 – 7.28 (m, 2H), 7.18 (d, J = 8.4 Hz, 2H), 6.90 (s, 1H), 4.18 (d, J = 10.7 Hz, 1H), 4.05 (d, J = 10.6 Hz, 1H), 3.48 (t, J = 8.4 Hz, 1H), 3.37 (d, J = 10.0 Hz, 1H), 3.04 (d, J = 10.0 Hz, 1H), 3.00 (dd, J = 9.5, 3.5 Hz, 1H), 2.66 (s, 3H), 2.35 (dt, J = 7.3, 3.5 Hz, 1H), 2.34 – 2.26 (m, 1H), 2.21 (dtd, J = 10.4, 6.3, 1.9 Hz, 1H), 1.95 – 1.87 (m, 2H), 1.57 – 1.53 (m, 1H), 1.31 (s, 9H), 1.29 (d, J = 8.5 Hz, 1H), 1.23 (s, 3H), 1.03 (s, 3H).

**$^{13}$C NMR** (176 MHz, $CD_2Cl_2$) δ 157.1, 153.5, 138.4, 138.3, 138.3, 133.0, 130.4, 130.3, 123.5, 120.8, 86.4, 70.6, 58.0, 56.6, 51.0, 46.7, 40.6, 39.3, 35.2, 34.8, 34.4, 31.1, 27.4, 27.4, 23.6, 21.2.

**HRMS** (ESI) for $C_{30}H_{39}IN_2O_4SNa$: calculated 651.1748 $[M+Na]^+$, found 651.1752.

$[\alpha]^{20}_D$ = +10.7° ($CHCl_3$, *c* = 0.5).

**((3a*S*,4*R*,6*R*,7a*S*)-2-((4-(*tert*-butyl)-2-methylphenyl)sulfonyl)-5,5-dimethyloctahydro-3a*H*-4,6-methanoisoindol-3a-yl)methyl (4-ethynylphenyl)carbamate 13 (iDeg-6):**

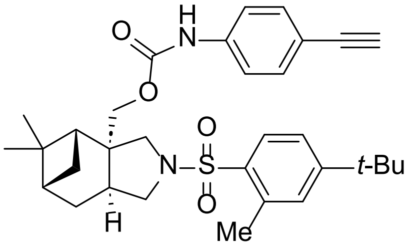

**13 (iDeg-6)**

To a degassed solution of **11** (89 mg, 0.14 mmol) in THF (1.4 mL) was added $NEt_3$ (98.3 µL, 0.7 mmol, 5.0 equiv.), Trimethylsilylacetylene (42 ul,0.28 mmol, 2.0 equiv.), CuI (5.3 mg, 0.2 equiv.) and $Pd(PPh_3)_4$ (16.2 mg, 0.014 mmol, 0.1 eq.), and the mixture was stirred at room temperature for 22 h. The reaction mixture was poured into sat. aq. NaCl solution, extracted with dichloromethane (2 ×), dried over $MgSO_4$ and the solvent was removed under reduced pressure. Column chromatography (PE/EtOAc 15:1 to 10:1 to 5:1) afforded a beige colored solid.

To a solution of the solid in THF (1.8 mL) was added TBAF (1 M in THF, 180 µL, 0.18 mmol, 1.2 equiv.) and the mixture was stirred at room temperature for 1 h. The solvent was removed under reduced pressure. Column chromatography (PE/EtOAc 15:1 to 10:1 to 5:1) afforded a light pink solid (53 % yield over 2 steps, >99% HPLC purity).

**$^1$H NMR** (600 MHz, $CD_2Cl_2$) δ 7.81 (d, *J* = 9.0 Hz, 1H), 7.45 – 7.41 (m, 2H), 7.36 (d, *J* = 8.4 Hz, 2H), 7.34 (d, *J* = 7.2 Hz, 2H), 6.95 (s, 1H), 4.10 (d, *J* = 2.7 Hz, 2H), 3.45 – 3.39 (m, 1H), 3.27 (d, *J* = 10.0 Hz, 1H), 3.09 (s, 1H), 3.02 (d, *J* = 10.0 Hz, 1H), 3.00 (dd, *J* = 10.3, 2.0 Hz,

1H), 2.64 (s, 3H), 2.32 (t, *J* = 3.1 Hz, 2H), 2.25 – 2.19 (m, 1H), 1.99 – 1.94 (m, 1H), 1.89 (dd, *J* = 5.5, 2.2 Hz, 1H), 1.30 (s, 9H), 1.29 (d, *J* = 5.7 Hz, 1H), 1.23 (s, 3H), 1.03 (s, 3H).
**$^{13}$C NMR** (151 MHz, $CD_2Cl_2$) δ 157.1, 153.4, 139.1, 138.3, 133.3, 133.0, 130.4, 130.3, 123.5, 118.5, 117.0, 83.6, 76.8, 70.6, 58.0, 56.6, 51.0, 46.7, 40.6, 39.3, 35.2, 34.9, 34.4, 31.1, 27.4, 27.4, 23.6, 21.2.

**HRMS** (ESI) for $C_{32}H_{40}N_2O_4SNa$: calculated: 571.2606 $[M+Na]^+$, found 571.2602.

$[\alpha]^{20}_D$ = +10.7° ($CHCl_3$, *c* = 0.5).

**General procedure for the synthesis of benzenesulfonyl chlorides (14):[20]**

Chlorosulfonic acid (5 equiv) was added dropwise to a solution of alkyl benzene (1 eq.) in chloroform at 0 °C. The resulting solution was stirred at room temperature for 1 h. The reaction mixture was poured onto ice, and the aqueous phase was extracted with chloroform twice. After the reaction was finished, the mixture was carefully poured over crushed ice and the aqueous layer was extracted with chloroform. The combined organic layers were then washed with brine, dried over sodium sulfate, and concentrated by rotary evaporation to give the crude product. The resulting oil was then purified by flash column chromatography (hexanes/ethyl acetate = 20:1, 35% yield).

**2-Methyl-4-(*tert*-butyl)-benzenesulfonyl chloride (14)**

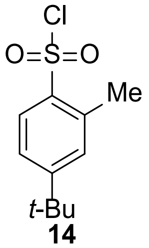

**14**

**$^1$H NMR** (500 MHz, $CDCl_3$) δ 7.98 (d, *J* = 9.1 Hz, 1H), 7.42 – 7.38 (m, 2H), 2.78 (s, 3H), 1.35 (s, 9H).
**$^{13}$C NMR** (126 MHz, $CDCl_3$) δ 159.6, 140.4, 137.9, 130.6, 128.9, 123.9, 35.4, 31.1, 20.7.
**LRMS** (EI) for $C_{11}H_{15}ClO_2S$: calculated: 246.05 [M], found 246.05.

**HPLC method to determine the stability of iDegs:**

| Time (min) | %A | %B |
|---|---|---|
| 0 | 90 | 10 |
| 1 | 90 | 10 |
| 3.3 | 50 | 50 |
| 6 | 5 | 95 |
| 8 | 5 | 95 |
| 10 | 90 | 10 |

A: Double distilled (dd) $H_2O$ + 0.1% (v/v) formic acid; B: acetonitrile + 0.1% (v/v) formic acid; flow rate: 0.25 mL/min. Instruments: Velos Pro (Thermo Fisher Scientific, US), Ultimate 3000 HPLC (Thermo Fisher Scientific, US).

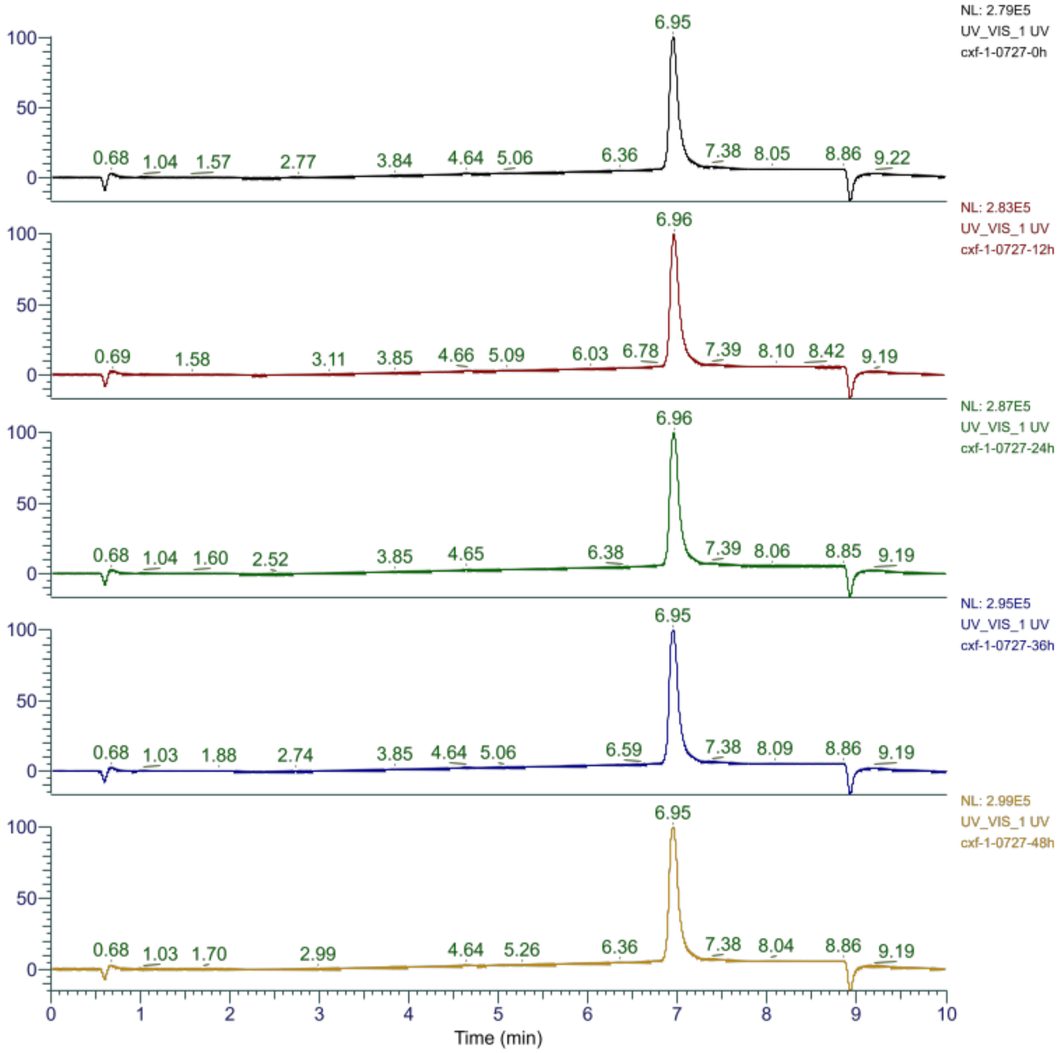

**Supporting Scheme 4**. Stability of iDeg-6. A 100 µM solution of iDeg-6 was incubated in $H_2O$/MeCN (v/v = 1/1) for up to 48 h at 37 °C and subsequently subjected to HPLC-ESI-MS analysis.

#### NMR-spectra

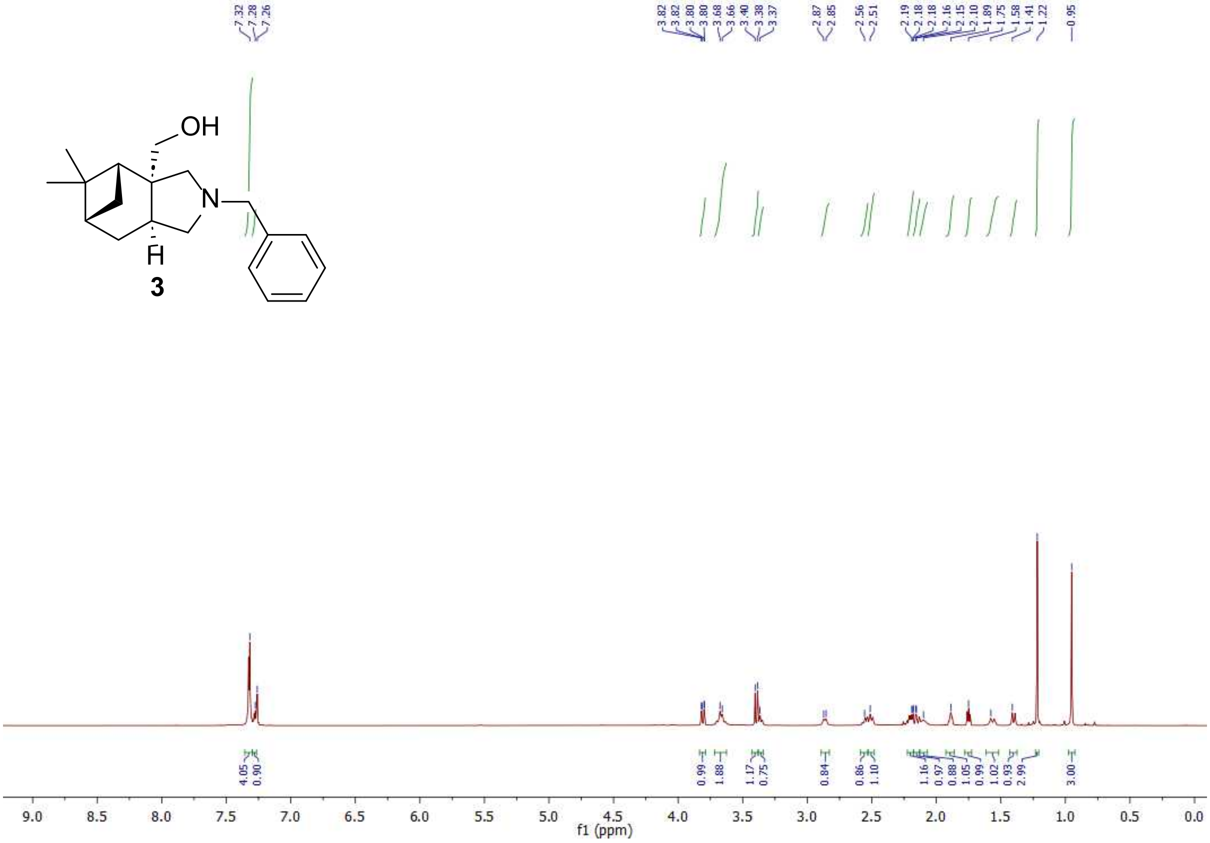

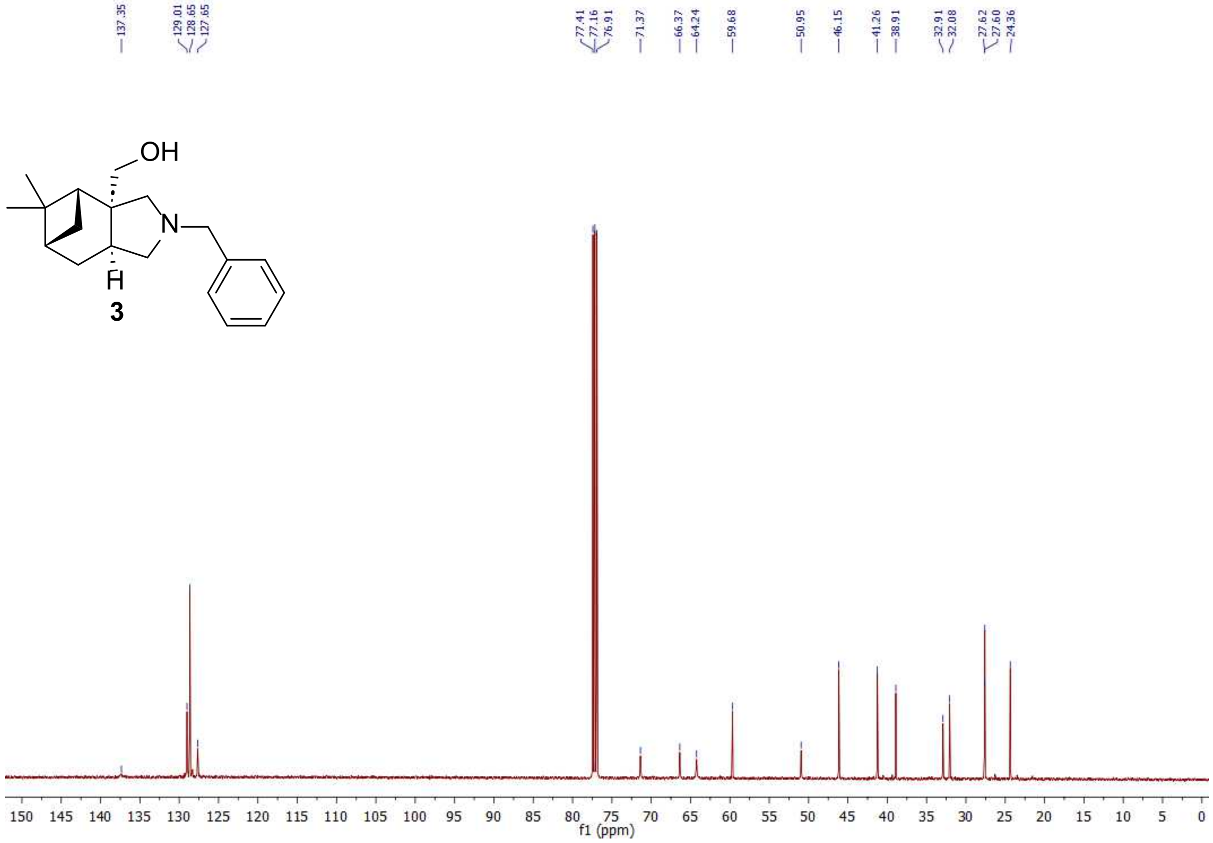

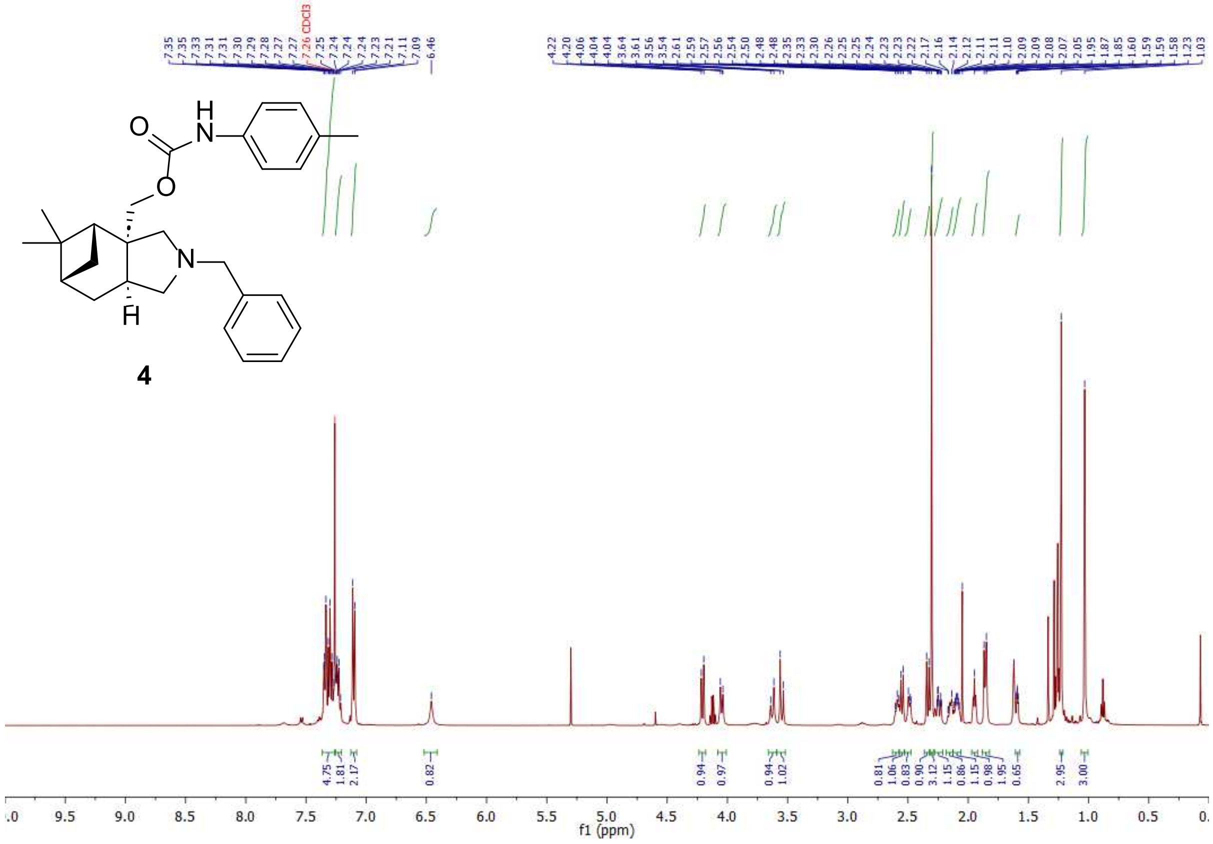
7.35 7.35 7.33 7.31 7.31 7.30 7.29 7.28 7.27 7.27 7.26 CDCl3 7.25 7.24 7.24 7.24 7.23 7.23 7.21 7.11 7.09 6.46
4.22 4.20 4.06 4.04 4.04 3.64 3.61 3.56 3.54 2.61 2.59 2.57 2.56 2.54 2.50 2.48 2.48 2.35 2.33 2.30 2.26 2.25 2.25 2.24 2.23 2.23 2.22 2.17 2.16 2.14 2.12 2.11 2.11 2.10 2.09 2.09 2.08 2.07 2.05 1.95 1.87 1.85 1.60 1.59 1.59 1.58 1.23 1.03
4
4.75 1.81 2.17 0.82 0.94 0.97 0.94 1.02 0.81 1.06 0.83 0.90 3.12 1.15 0.86 1.15 0.98 1.95 0.65 2.95 3.00
f1 (ppm)

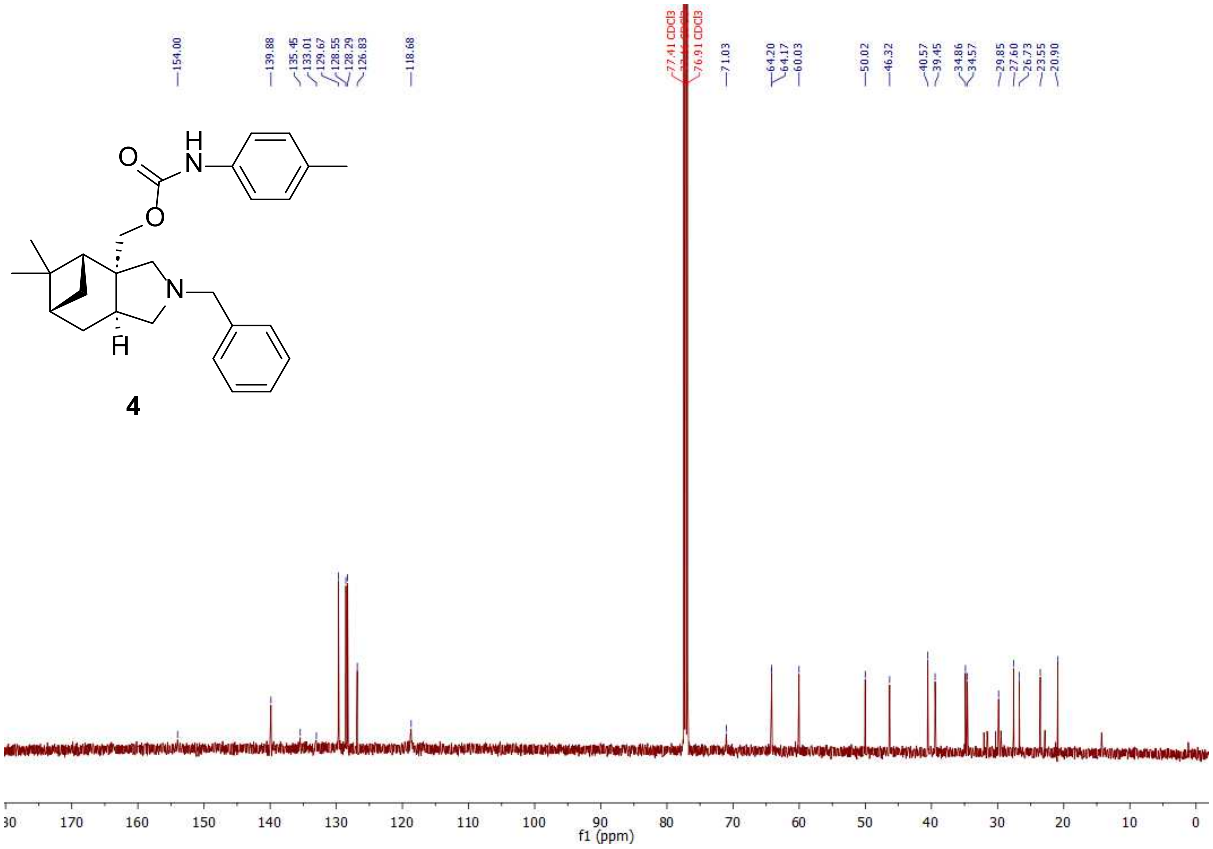
154.00 139.88 135.45 133.01 129.67 128.55 128.29 126.83 118.68
77.41 CDCl3 76.91 CDCl3
71.03 64.20 64.17 60.03 50.02 46.32 40.57 39.45 34.86 34.57 29.85 27.60 26.73 23.55 20.90
4
f1 (ppm)

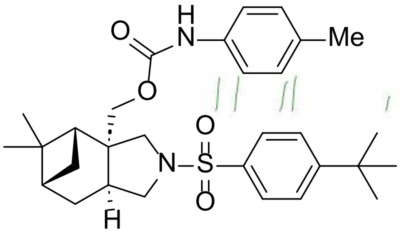

**iDeg-1**

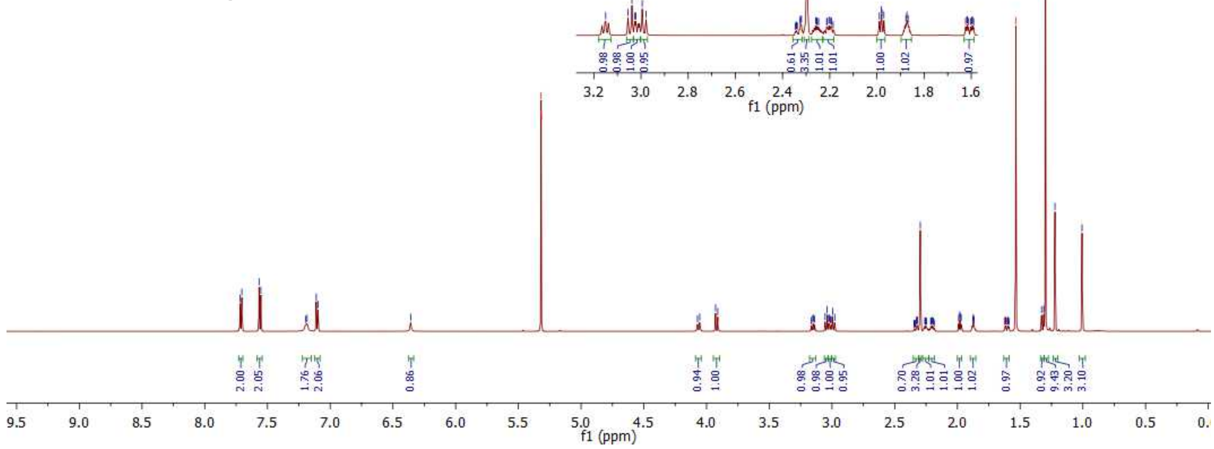

7.72 7.70 7.57 7.55 7.20 7.18 7.11 7.10 6.36
5.32 CD2Cl2 5.32 CD2Cl2 5.32 CD2Cl2
4.07 4.05 3.93 3.91 3.17 3.15 3.15 3.14 3.05 3.04 3.03 3.02 3.01 3.01 2.99 2.98 2.35 2.34 2.34 2.33 2.32 2.32 2.30 2.26 2.25 2.25 2.21 2.21 2.20 2.20 2.19 2.19 1.99 1.98 1.98 1.97 1.88 1.87 1.87
1.62 HDO
1.62 1.61 1.60 1.60 1.59 1.59
1.53 HDO
1.33 1.31 1.30 1.22 1.01
3.15 3.05 3.04 3.03 3.02 2.99 2.98 2.33 2.32 2.32 2.30 2.26 2.25 2.21 2.21 2.20 1.99 1.98 1.97 1.88 1.87 1.62 1.60 1.60 1.59
0.98 0.98 1.00 0.95 0.61 3.35 1.01 1.01 1.00 1.02 0.97
3.2 3.0 2.8 2.6 2.4 2.2 2.0 1.8 1.6
f1 (ppm)
2.00 2.05 1.76 2.06 0.86 0.94 1.00 0.98 0.98 1.00 0.95 0.70 3.28 1.01 1.01 1.00 1.02 0.97 0.92 9.43 3.20 3.10
9.5 9.0 8.5 8.0 7.5 7.0 6.5 6.0 5.5 5.0 4.5 4.0 3.5 3.0 2.5 2.0 1.5 1.0 0.5 0.
f1 (ppm)

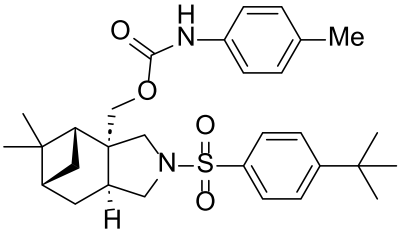

**iDeg-1**

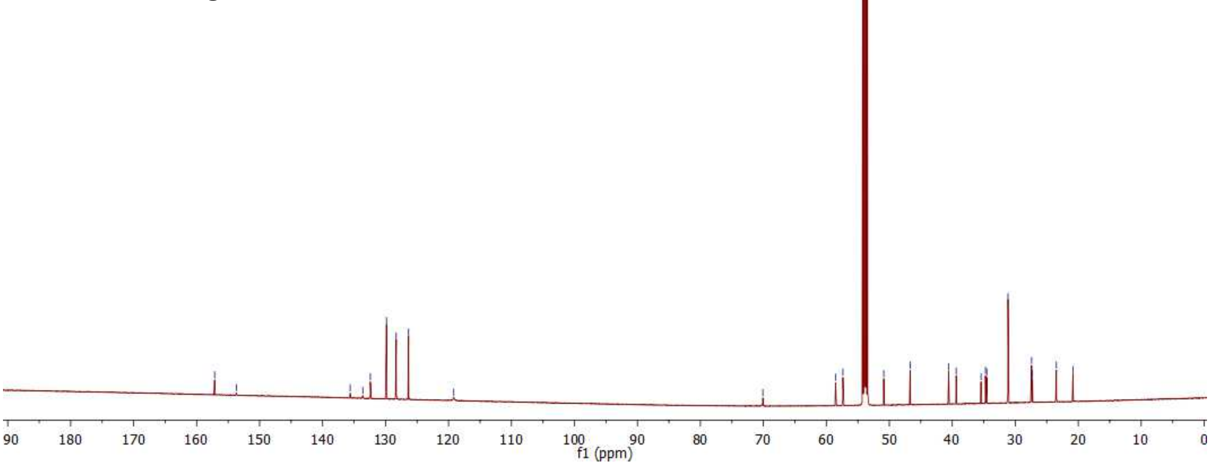

157.15 153.70 135.62 133.62 132.41 129.85 128.31 126.35 119.17 70.09 58.53 57.37 54.20 54.02 53.84 53.66 53.48 50.88 46.70 40.57 39.38 35.44 34.73 34.52 31.15 27.40 27.27 23.47 20.83
90 180 170 160 150 140 130 120 110 100 90 80 70 60 50 40 30 20 10 0
f1 (ppm)

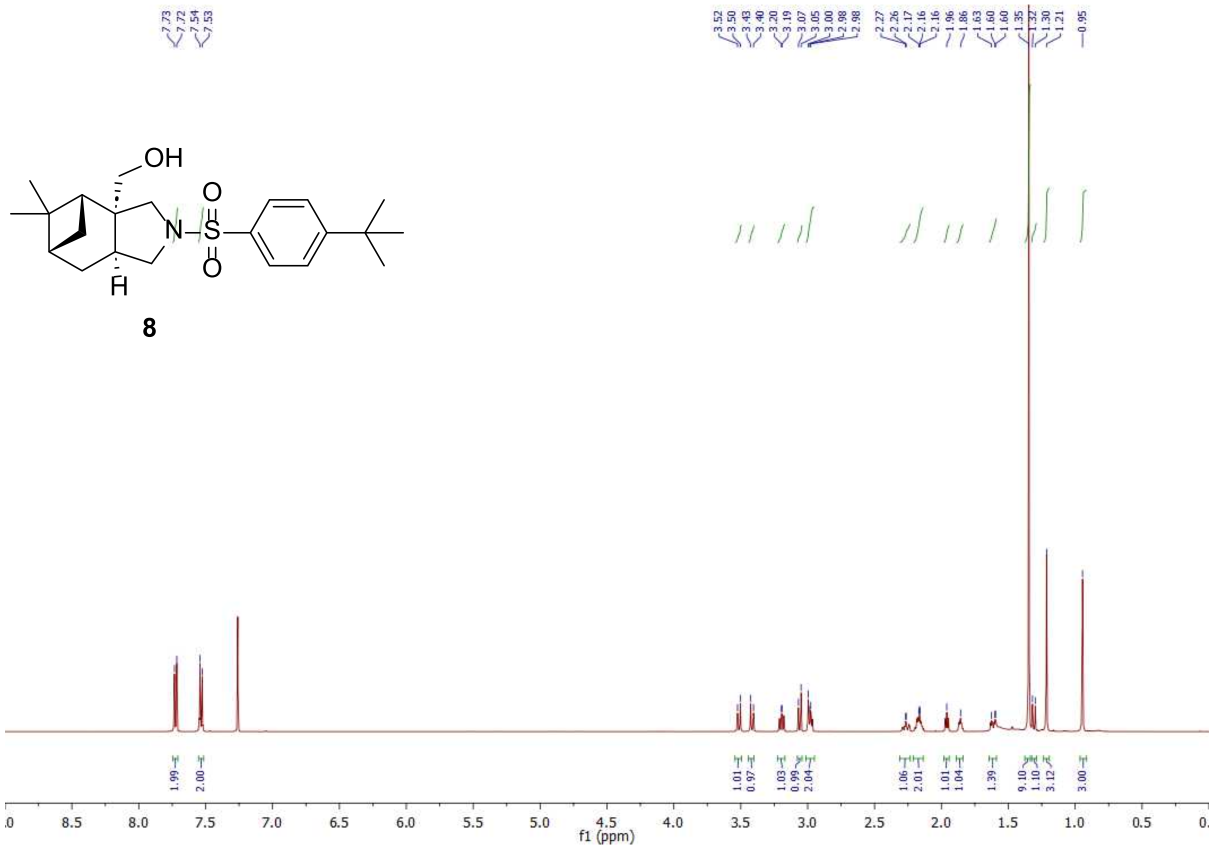
7.73
7.72
7.54
7.53
3.52
3.50
3.43
3.40
3.20
3.19
3.07
3.05
3.00
2.98
2.98
2.27
2.26
2.17
2.16
2.16
1.96
1.86
1.63
1.60
1.60
1.35
1.32
1.30
1.21
0.95
OH
O
N
S
O
H
8
1.99
2.00
1.01
0.97
1.03
0.99
2.04
1.06
2.01
1.01
1.04
1.39
9.10
1.10
3.12
3.00
f1 (ppm)

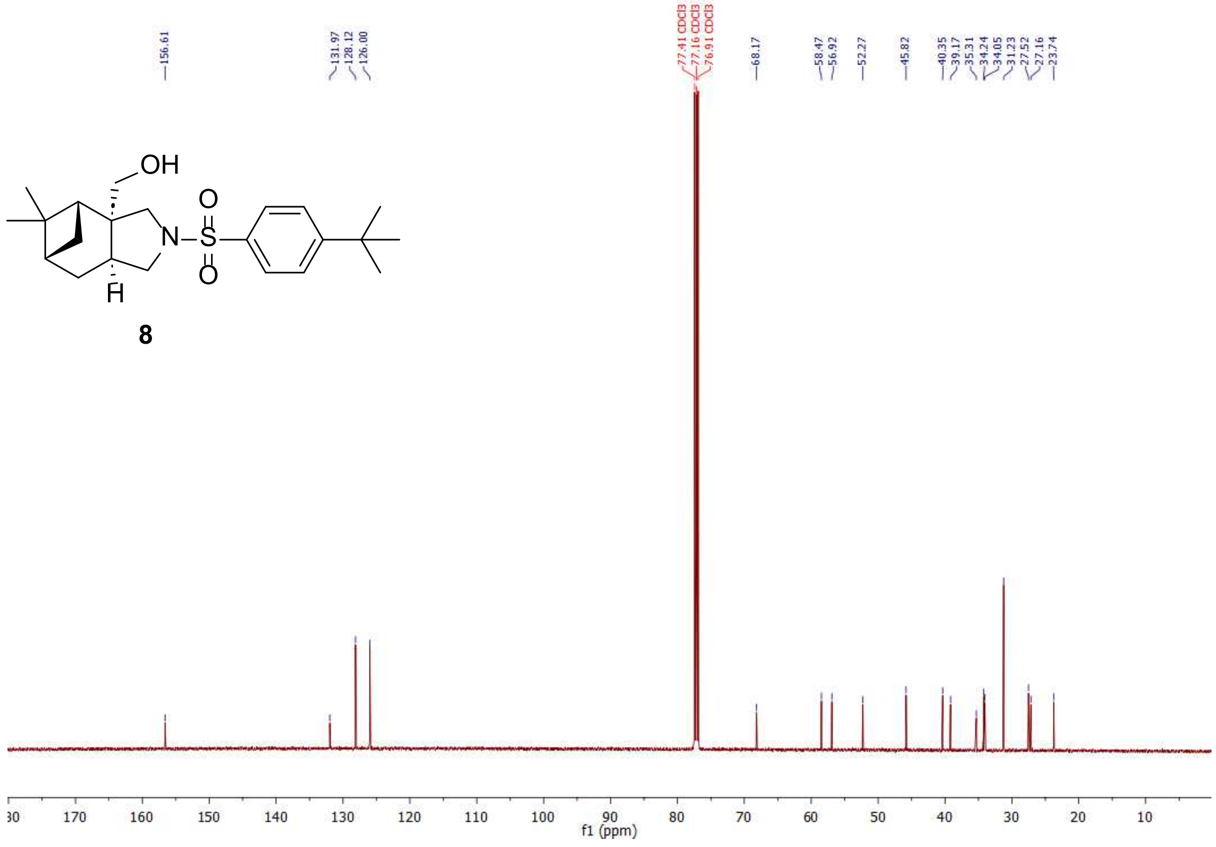
156.61
131.97
128.12
126.00
77.41 CDCl3
77.16 CDCl3
76.91 CDCl3
68.17
58.47
56.92
52.27
45.82
40.35
39.17
35.31
34.24
34.05
31.23
27.52
27.16
23.74
OH
O
N
S
O
H
8
f1 (ppm)

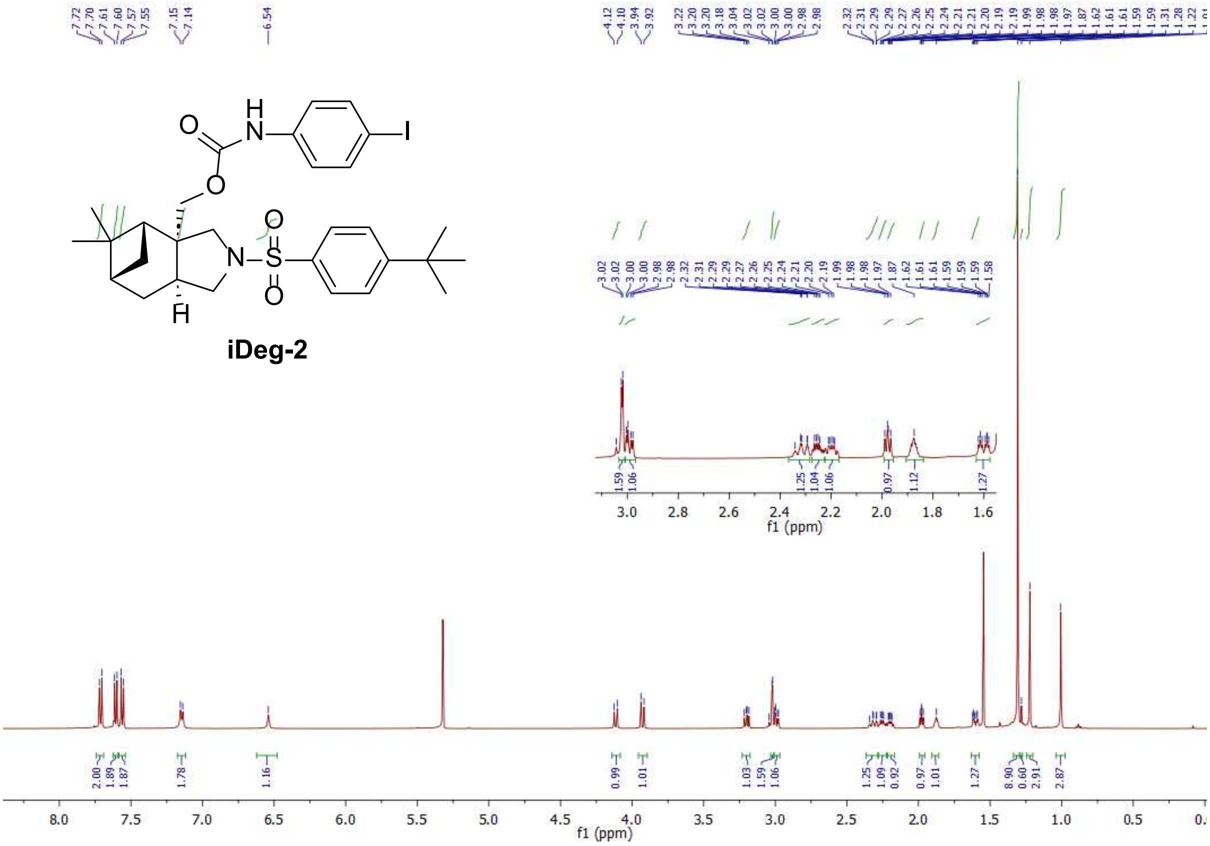
iDeg-2
f1 (ppm)

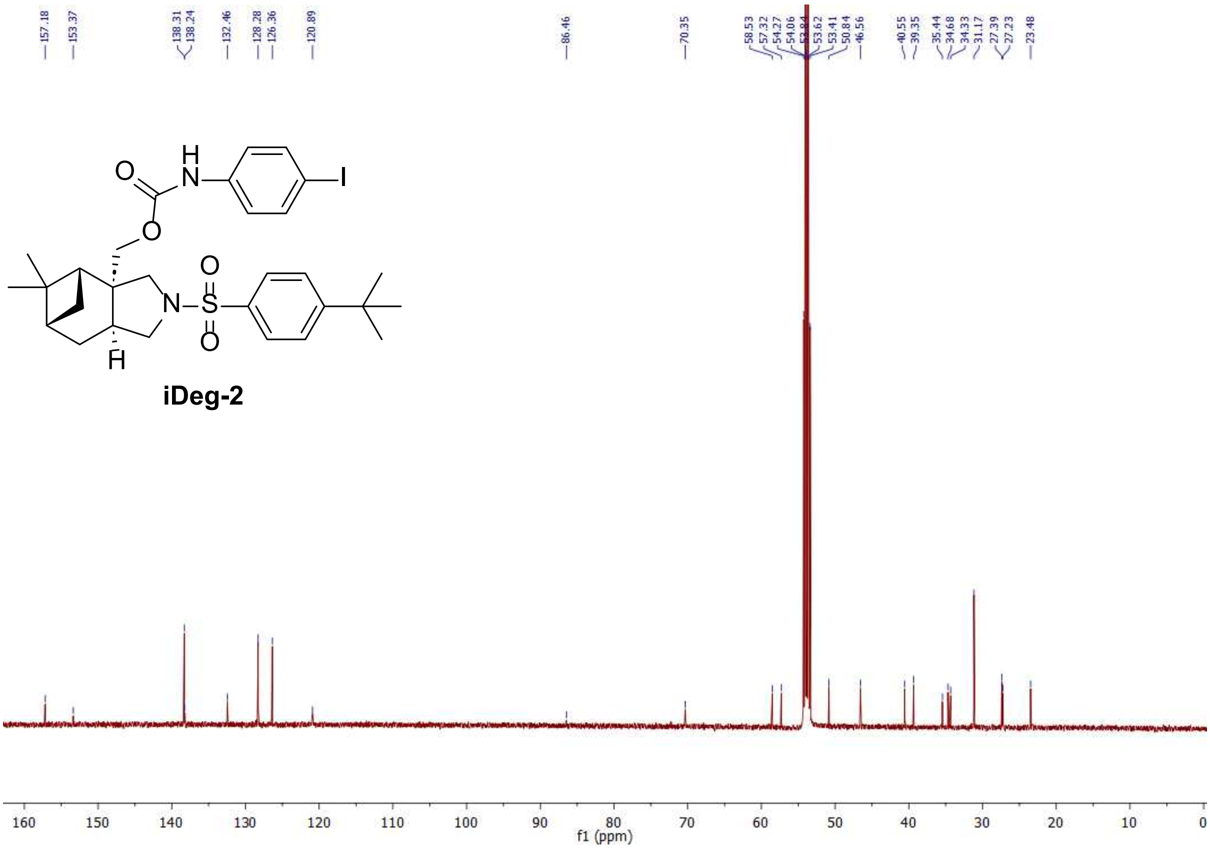
157.18
153.37
138.31
138.24
132.46
128.28
126.36
120.89
86.46
70.35
58.53
57.32
54.27
54.06
53.84
53.62
53.41
50.84
46.56
40.55
39.35
35.44
34.68
34.33
31.17
27.39
27.23
23.48
iDeg-2
f1 (ppm)

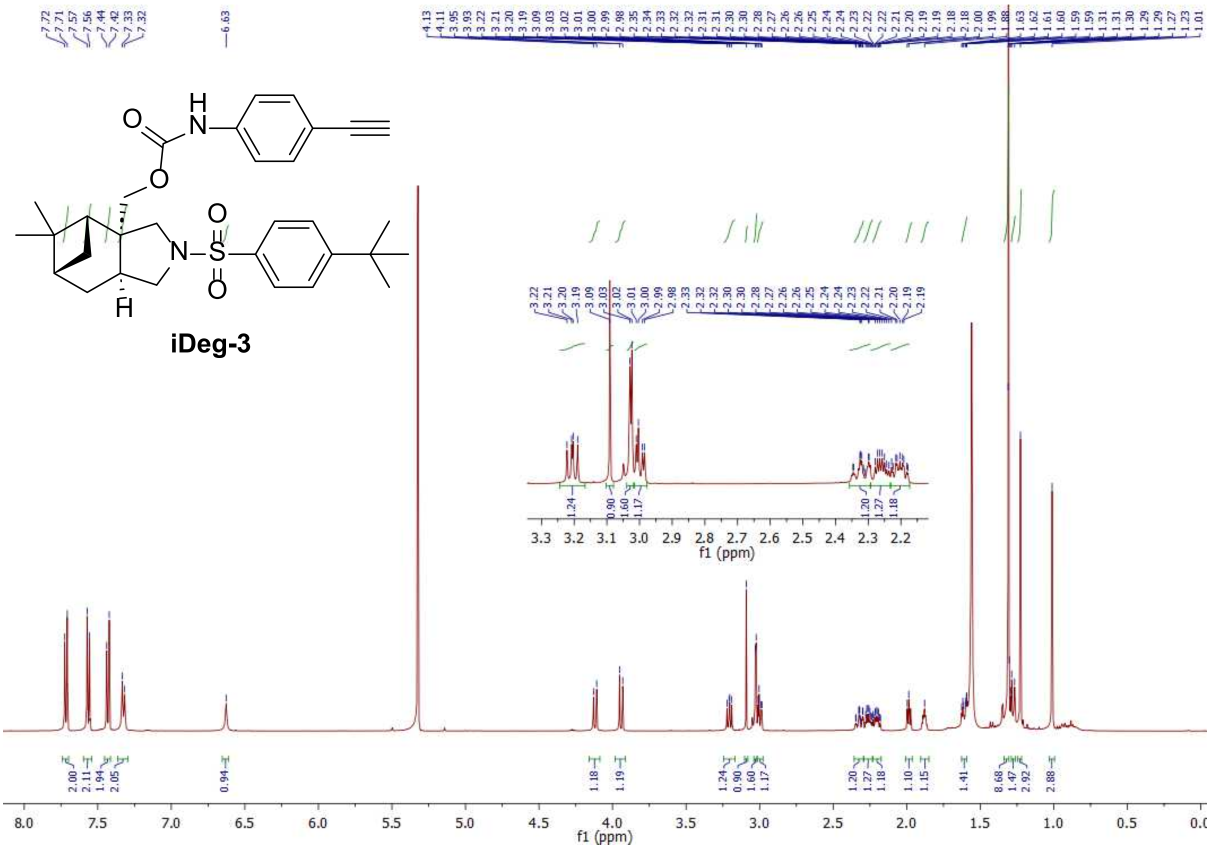
iDeg-3
f1 (ppm)

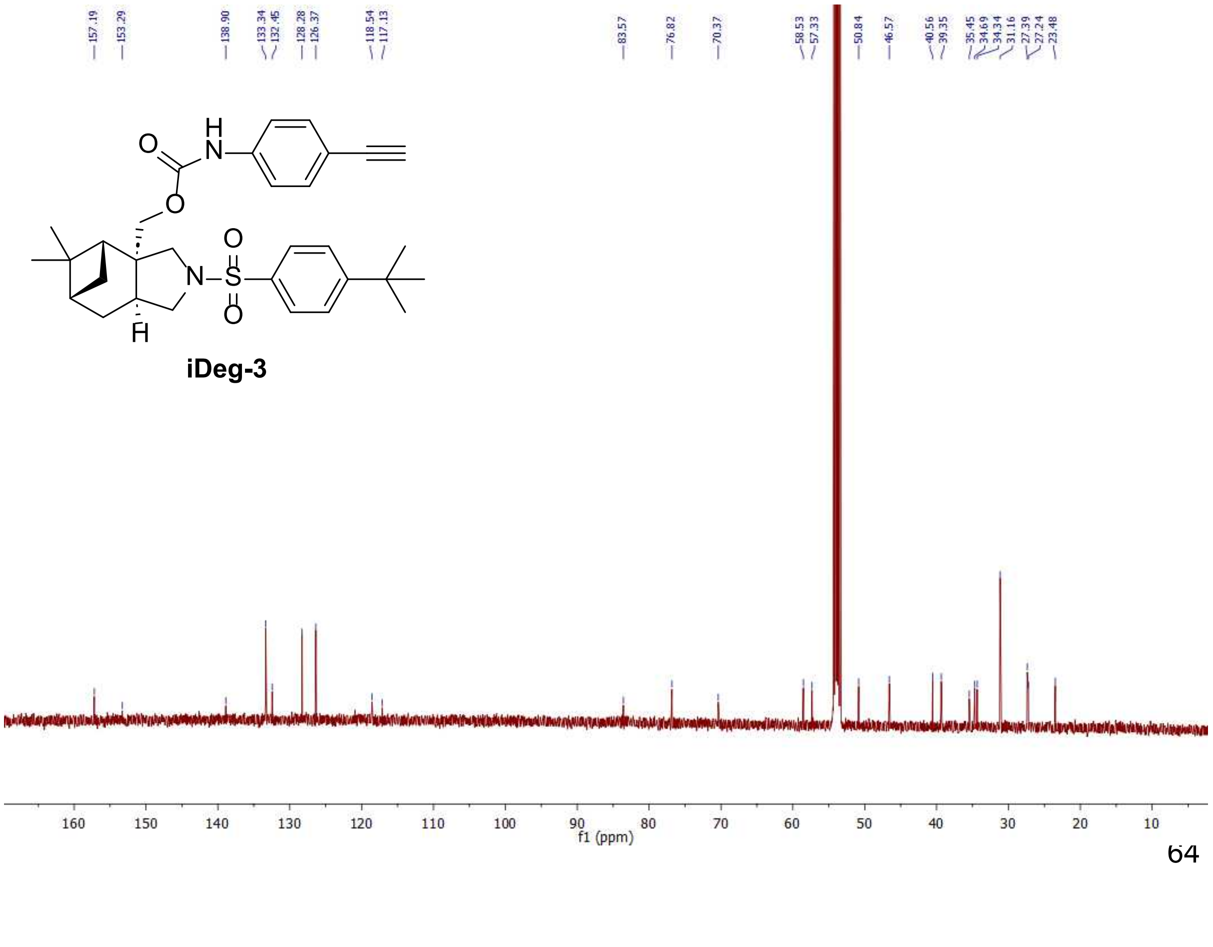
iDeg-3
f1 (ppm)

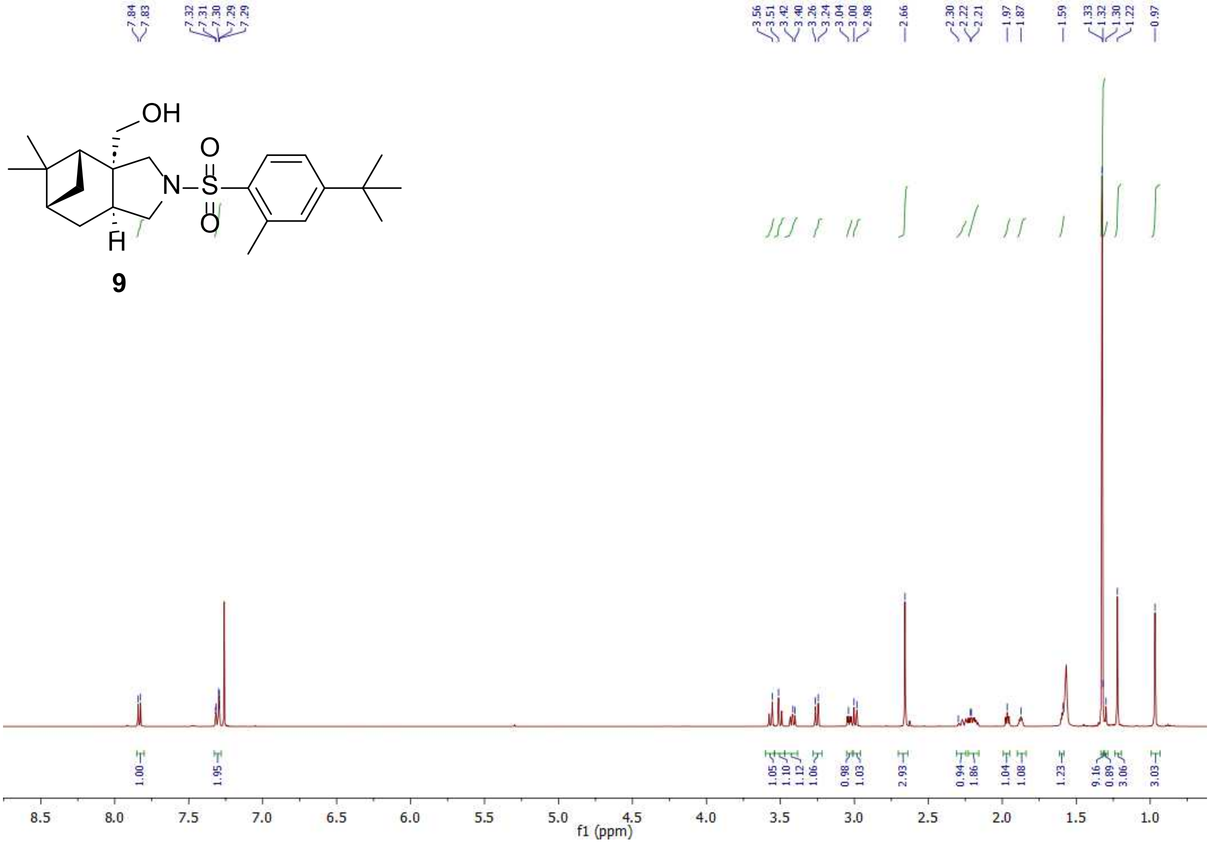
7.84
7.83
7.32
7.31
7.30
7.29
7.29
3.56
3.51
3.42
3.40
3.26
3.24
3.04
3.00
2.98
2.66
2.30
2.22
2.21
1.97
1.87
1.59
1.33
1.32
1.30
1.22
0.97
OH
O
S
N
O
H
9
1.00
1.95
1.05
1.10
1.12
1.06
0.98
1.03
2.93
0.94
1.86
1.04
1.08
1.23
9.16
0.89
3.06
3.03
f1 (ppm)

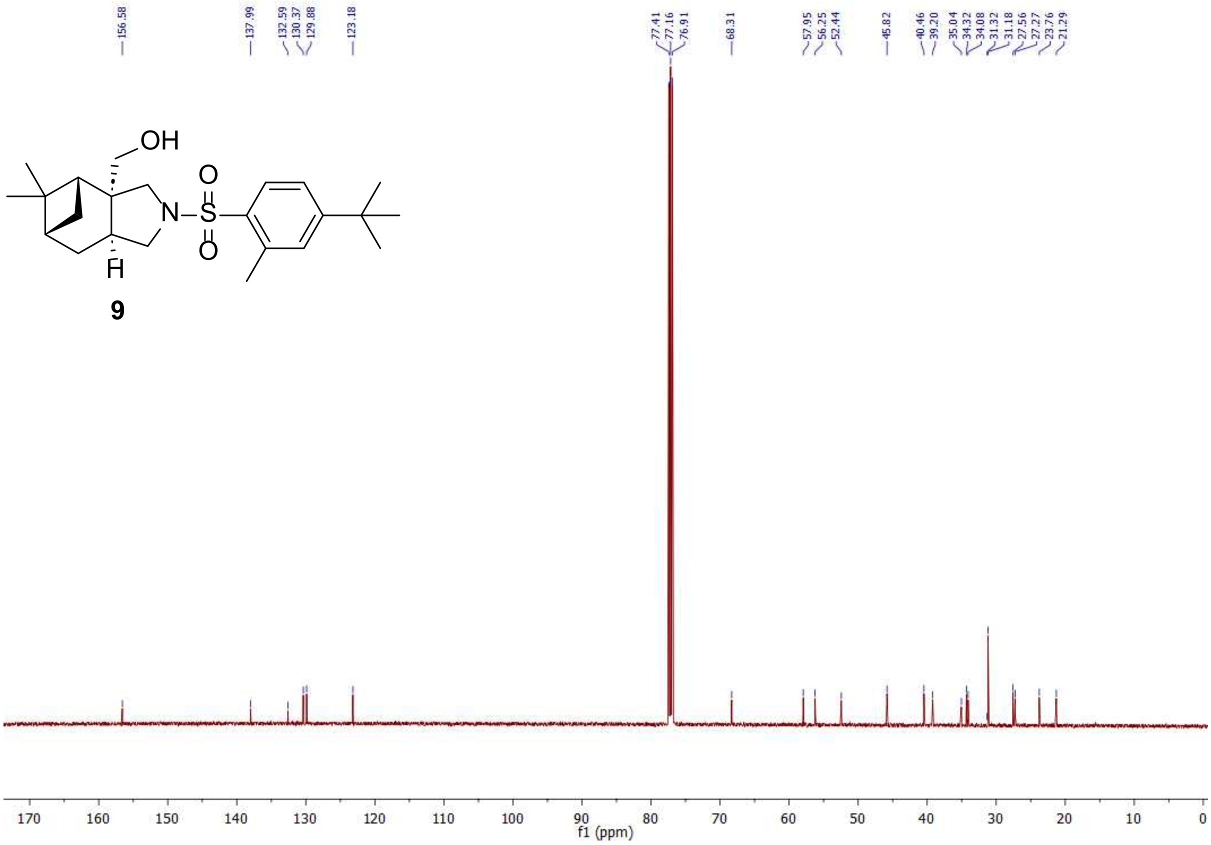
156.58
137.99
132.59
130.37
129.88
123.18
77.41
77.16
76.91
68.31
57.95
56.25
52.44
45.82
40.46
39.20
35.04
34.32
34.08
31.32
31.18
27.56
27.27
23.76
21.29
OH
O
S
N
O
H
9
f1 (ppm)

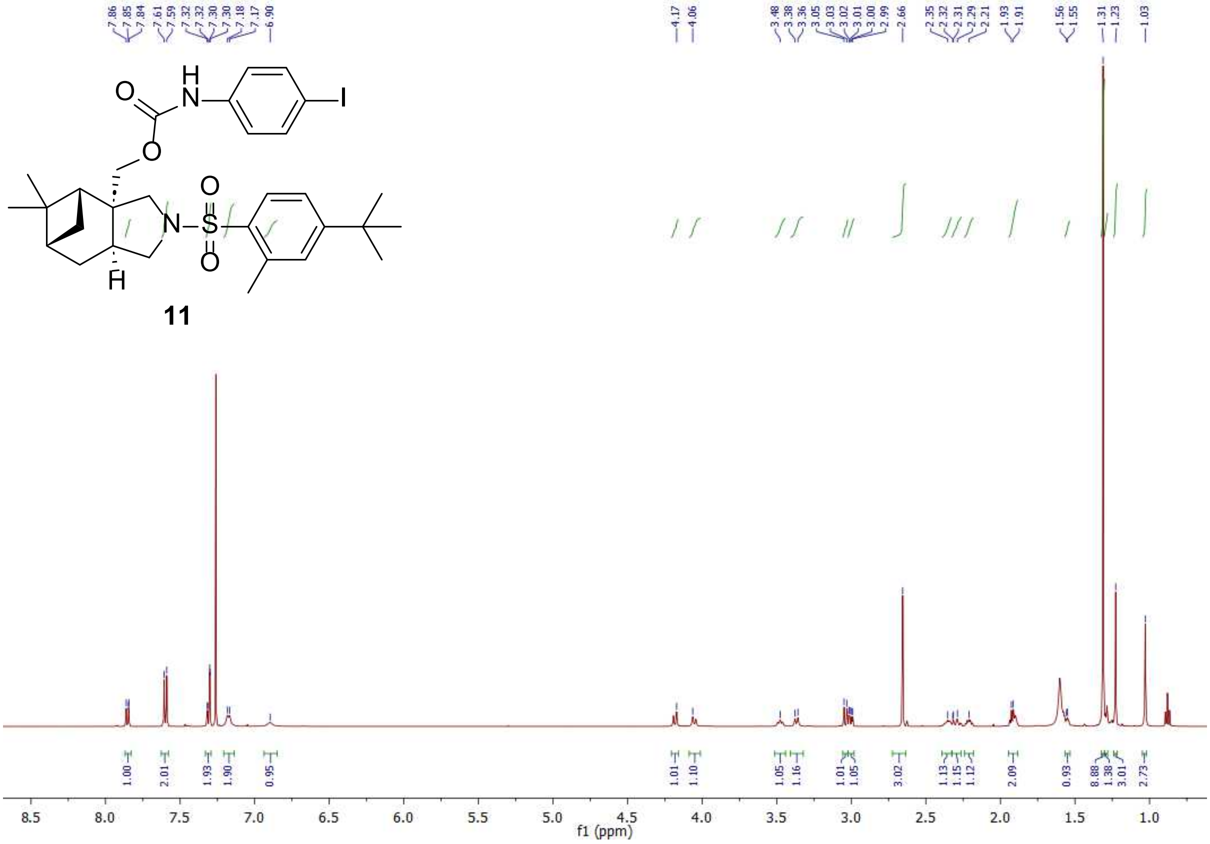
11
f1 (ppm)

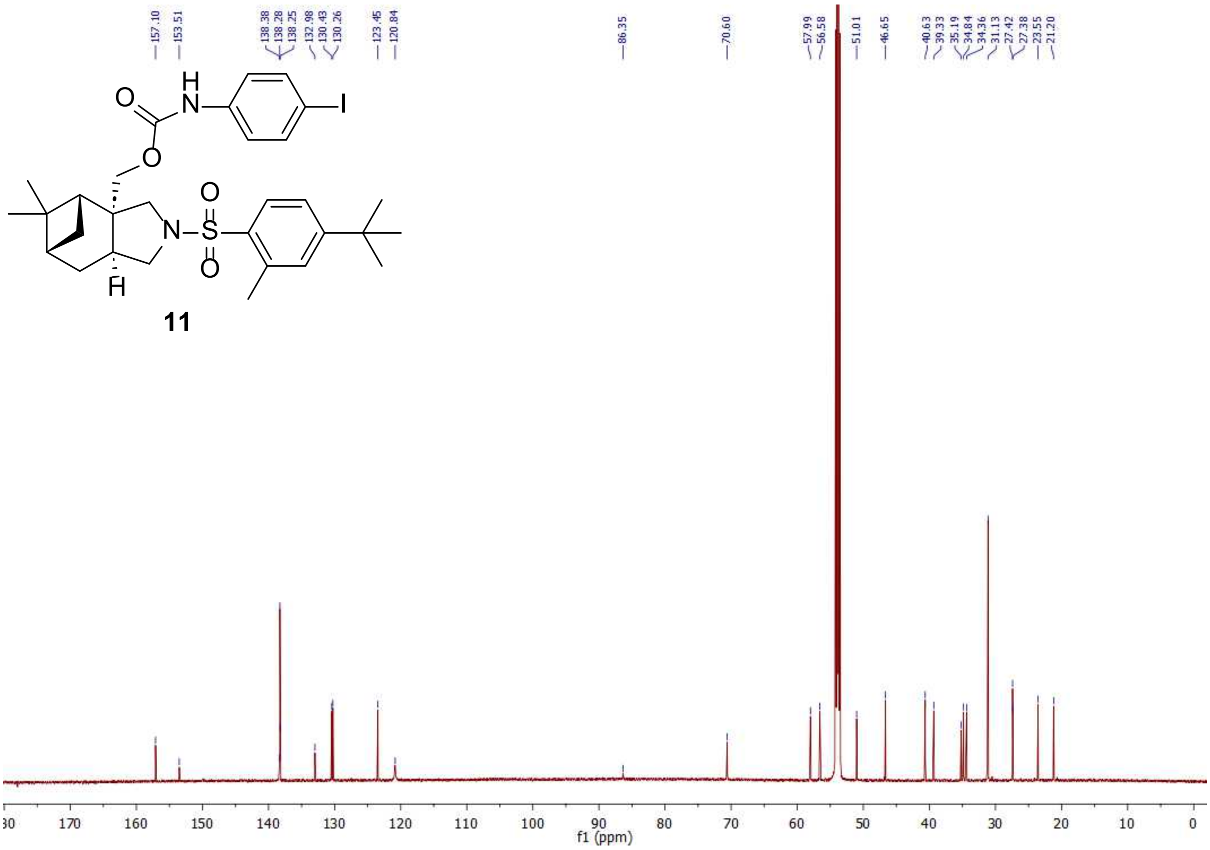
11
f1 (ppm)

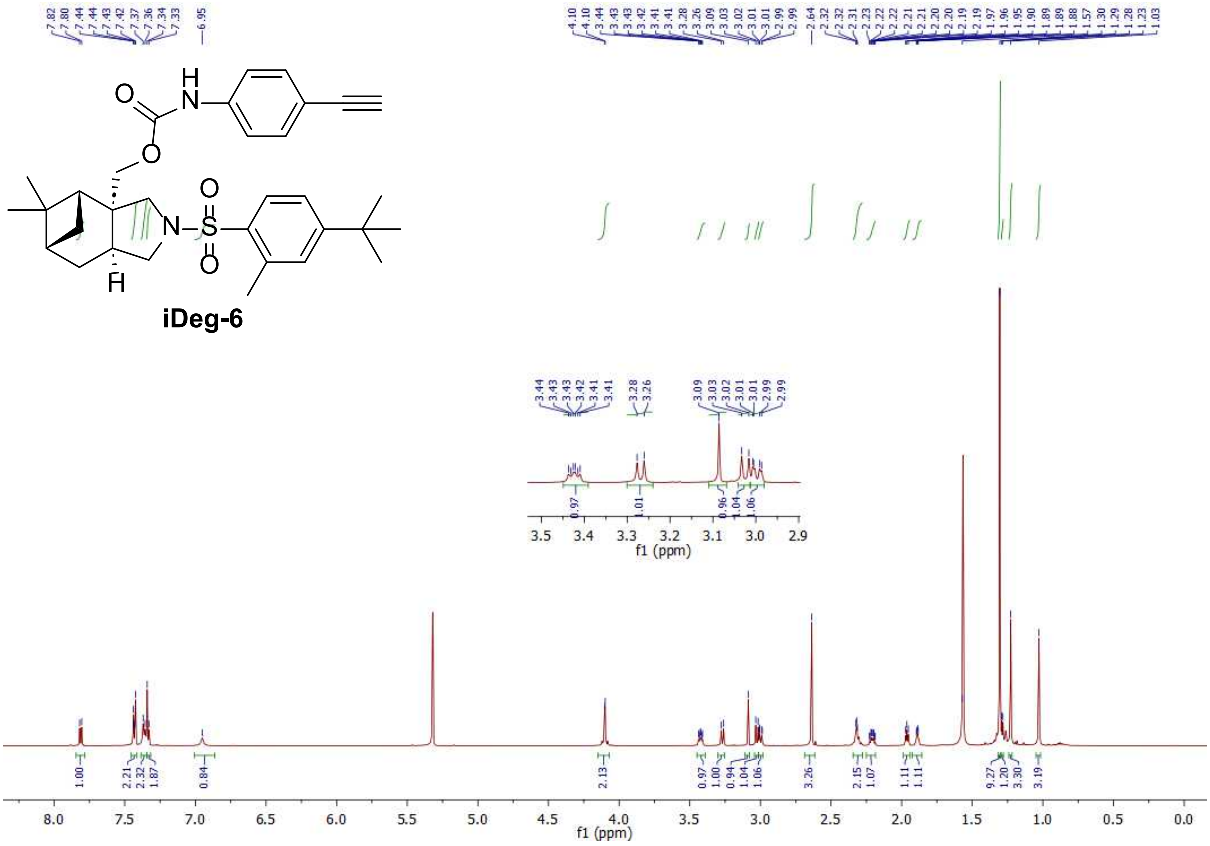
iDeg-6
f1 (ppm)

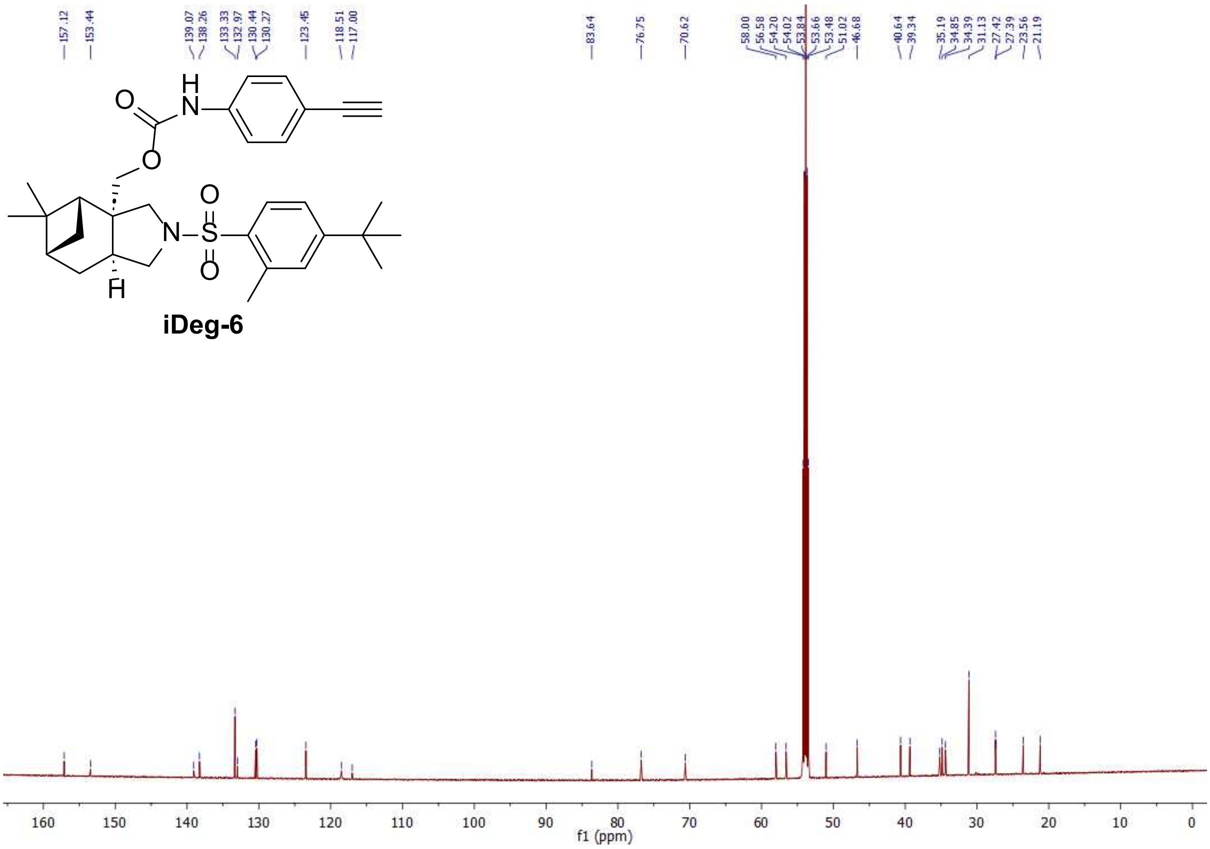
iDeg-6
f1 (ppm)

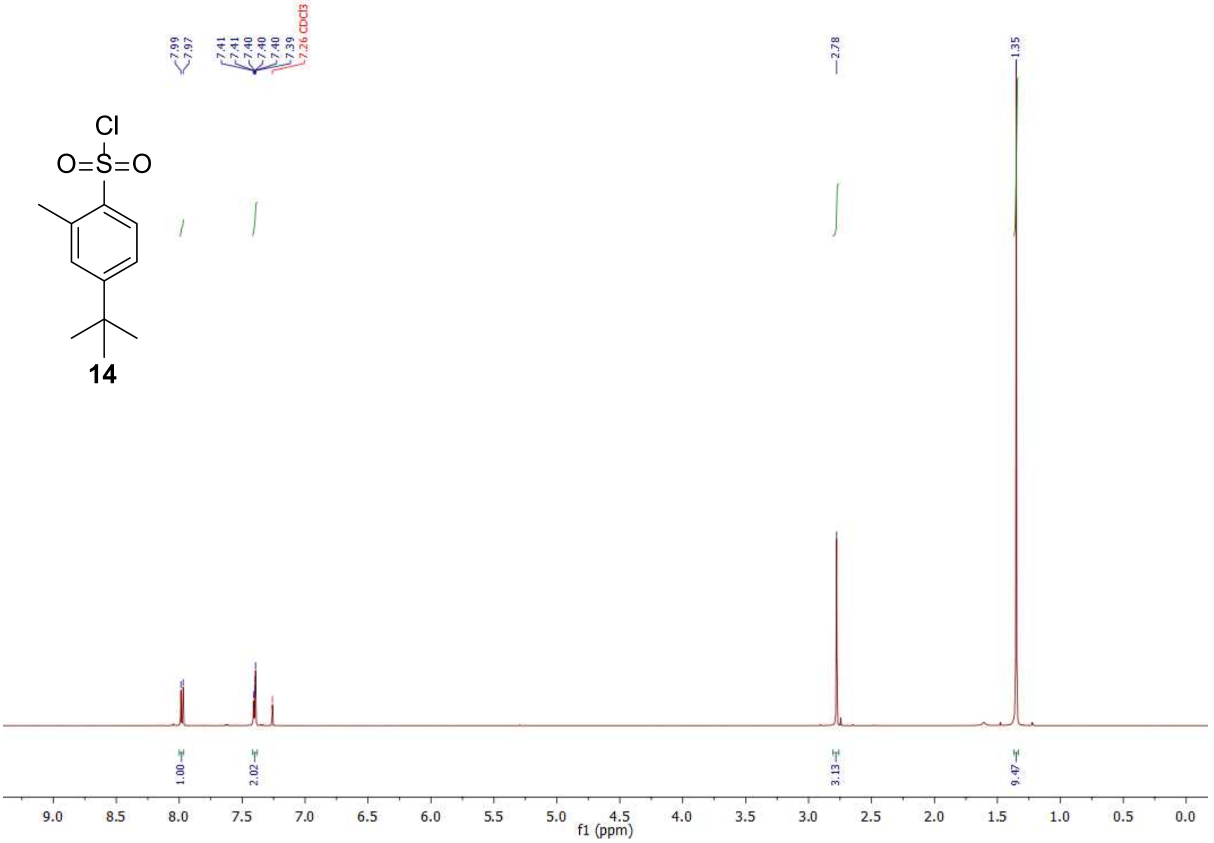
7.99
7.97
7.41
7.41
7.40
7.40
7.40
7.39
7.26 CDCl3
2.78
1.35
Cl
O=S=O
14
1.00
2.02
3.13
9.47
9.0 8.5 8.0 7.5 7.0 6.5 6.0 5.5 5.0 4.5 4.0 3.5 3.0 2.5 2.0 1.5 1.0 0.5 0.0
f1 (ppm)

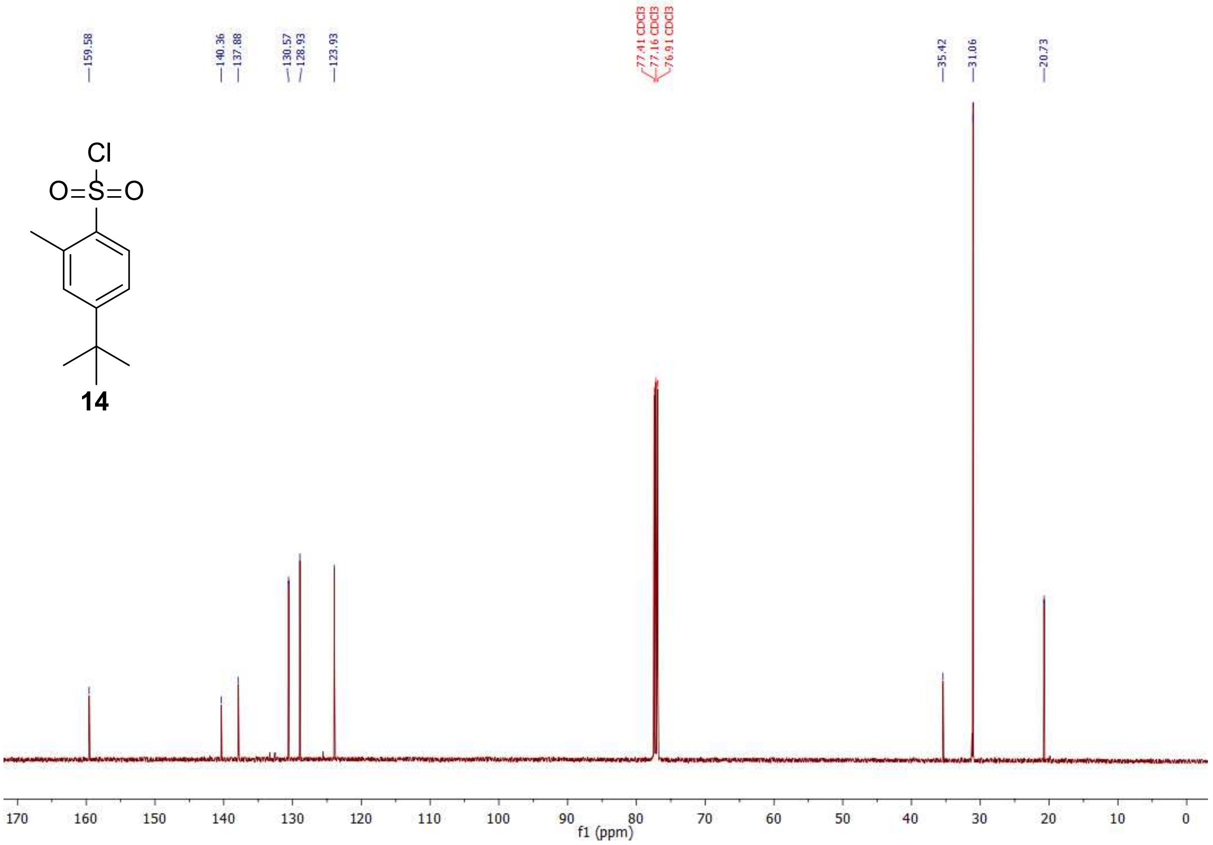
159.58
140.36
137.88
130.57
128.93
123.93
77.41 CDCl3
77.16 CDCl3
76.91 CDCl3
35.42
31.06
20.73
Cl
O=S=O
14
170 160 150 140 130 120 110 100 90 80 70 60 50 40 30 20 10 0
f1 (ppm)
